# Bacteriophage and Antibiotic Resistance Are Positively Associated across a Phylogenetically Diverse Set of Clinical *Pseudomonas aeruginosa* Isolates

**DOI:** 10.64898/2026.09.27.754834

**Authors:** Anna L. Rodden, Robert W. Targ, Zachary N. Flamholz, Paul L. Bollyky, Jessica C. Sacher

## Abstract

Co-administration of phages and antibiotics has been proposed as a therapeutic approach against antibiotic-resistant bacteria. The relationship, however, between antibiotic resistance and phage resistance in clinical isolates is unclear. Here, we examine associations between phage and antibiotic resistance profiles across a panel of *Pseudomonas aeruginosa* clinical isolates from the Centers for Disease Control (CDC) and Food and Drug Administration (FDA) Antimicrobial Resistance Isolate (ARI) Bank comprising 55 clinical strains with full genome sequences and antibiotic susceptibility testing (AST) data for 11 clinically relevant antibiotics. As phages in this study, we use three well-characterized, morphologically distinct phages, OMKO1, Luz19, and PAML31-1. We screen for phage resistance using a growth suppression assay, then conduct statistical analysis against antibiotic MIC (Minimum Inhibitory Concentration) data provided by the CDC to define association patterns across this dataset. We find multiple significant susceptibility correlations between pairs of antibiotics and phages, and a positive overall association between average phage resistance and antibiotic resistance across the 55 strains, even controlling for phylogenetic associations (*r*=0.358, p<0.005). We conclude that phage and antibiotic resistance are positively associated across this clinical isolate collection, suggesting that the two resistance phenotypes are not independent in *P. aeruginosa*. These findings have implications for the development of phage-antibiotic cocktails.

**Importance:** Antimicrobial resistance causes substantial and increasing global mortality. Understanding how bacteria develop resistance to different antimicrobial agents, particularly antibiotics and bacteriophages, is essential for identifying shared and distinct resistance mechanisms and developing more effective therapies.

## Introduction

Antimicrobial-resistant (AMR) infections pose an urgent threat to public health, with recent estimates that AMR directly contributed to the deaths of 1.27 million people and was associated with the deaths of nearly 5 million people globally in 2019 [1]. A recent study suggests that this annual death toll could increase by nearly 70% over the next two decades, resulting in 1.91 million directly attributable and 8.22 million associated fatalities in 2050 [2].

*Pseudomonas aeruginosa* is an opportunistic Gram-negative bacterium that exhibits high levels of AMR. Certain *Pseudomonas aeruginosa* lineages (“high-risk clones”) are disproportionately associated with multidrug resistance (MDR), extensive drug resistance (XDR), and even pan-drug resistance (PDR) [3, 4]. While antibiotic resistance can arise through a combination of intrinsic, acquired, and adaptive/evolutionary mechanisms, *P. aeruginosa* multidrug resistance typically emerges from the accumulation of multiple chromosomal mutations rather than acquisition of a single resistance plasmid [5, 6].

Given this extensive antibiotic resistance, *P. aeruginosa* is a frequent target of bacteriophage (phage) therapy: the use of lytic phages to target bacterial infections [7–10]. However, as with antibiotic resistance, bacterial resistance to phages also occurs via multiple mechanisms, including surface receptor variation [11], restriction-modification systems [12], CRISPR-Cas systems [13], and other anti-phage defense systems [14]. Certain successful clinical clones carry unusually large repertoires of defense systems, particularly prophage-encoded systems [15].

Efforts to counter both phage and antibiotic resistance have led to the development of ‘cocktails’ that combine both phages and antibiotics with the goal of applying orthogonal selection pressures to prevent emergence of resistant strains [11, 16–18]. The behavior of phage-antibiotic cocktails can be difficult to predict with both synergy and antagonism being possible [19–20]. Thus, is it essential to examine these interactions in a way that allows us to better predict combinatorial efficacy. There are emerging mechanistic investigations of phage-antibiotic interactions that may yield more predictable phage-antibiotic activity [10, 21–22].

While these and other studies have proactively examined phage-antibiotic interactions in vitro and in vivo [23–25], at present the prevalence and strength of associations between pre-existing phage and antibiotic resistance are unclear. Several studies examining *P. aeruginosa* isolates have found that many therapeutic phages retain activity against MDR strains [26–27], but these relationships have not been systematically evaluated, to our knowledge. Complicating matters, closely-related bacterial strains are known to share both phage host-range [28] and antibiotic resistance patterns [29], although there is also evidence of considerable within-lineage heterogeneity [30].

Here, we have examined the prevalence of phage and antibiotic co-resistance in a set of 55 well-characterized clinical isolate strains for which full genome sequences and antibiotic susceptibility testing (AST) data for a panel of 11 clinically relevant antibiotics were available. As phages in this study, we used three well-characterized, morphologically distinct phages with well-established receptor usage patterns and full genomes available. Using these phages and bacterial strains, we evaluated pairwise correlations between phage susceptibility and AST data and then examined the overall association between average phage resistance and antibiotic resistance whilst controlling for phylogeny. We also examined patterns of phage and antibiotic susceptibility by strain source, and finally tested for associations between particular resistance genes and phage-antibiotic cross-resistance.

## Results

### Selection of phages and bacterial strains

As clinical isolates in this study, we used a set of *P. aeruginosa* strains isolated from antibiotic resistant infections in clinics which we obtained from the CDC & FDA Antimicrobial Resistance Isolate Bank. The strains vary in level of resistance, with strains falling into the internationally recognized categories of MDR, XDR, and PDR, as well as none of the above (**Table 1**). The CDC & FDA Antimicrobial Resistance Isolate Bank provided AST data in the form of MIC values, with susceptible/intermediate/resistant breakpoints adapted from the CLSI 2025 standards. In downstream testing, we assigned the S/I/R classifications to numerical values, with 0 meaning resistant, 0.5 meaning intermediate, and 1 meaning susceptible. Directionality was assigned in order to match that of the phage AST values (suppression index data, as described below).

As phages in this study, we selected three phages which each target distinct *P. aeruginosa* receptors, and are thus members of three different ‘receptor complementarity groups’ [11, 31–33]. These phages represent different morphologies and have differently sized genomes. OMKO1 has a 281,755-bp genome [34] and has been shown to use the porin OprM and flagella as receptors [34–35, 11]. PAML31-1 has a 66,117 bp genome and binds lipopolysaccharide [11]. Luz19 has a 43,548-bp genome and binds the Type IV pilus [36, 37] (**Table 2**). These three phages have been extensively characterized.

### More closely related strains display more similar resistance profiles to antibiotics and phages

We next examined the susceptibility of our set of 55 clinical isolates to these three phages using a spectrophotometer to provide a robust, quantitative measurement of bacterial growth by recording the optical density at a wave-length of 600 nm (OD600) over time. From this data we generated a suppression index as described previously by taking the area under the curve (AUC) of the growth curves under each condition and subtracting the AUC under the phage-treated condition from the AUC under the control condition, then dividing this value by the AUC of the control condition [11]. We expressed this suppression index over a scale of <0-1, with negative values representing strains for which the phage-treated condition grew more than the control, values closer to 0 representing strains for which the phage-treated condition grew similarly or only slightly less than the control (indicating less suppression/more resistance), and values closer to 1 representing strains for which the phage-treated condition grew significantly less than the control or didn’t grow at all (indicating high suppression/more susceptibility).

We then performed a set of pairwise comparisons of these phage susceptibility data and the bacterial AST data for these clinical isolates. A similar strategy has been employed in *E. coli*, where specific pairwise associations were shown to predict uncharacterized resistance mechanisms [38], demonstrating the more direct clinical applications of this approach.

It is important to consider the role of genetic relatedness in these associations. To examine this, we constructed phylogenetic trees using the allele profiles for each of the 55 strains given by Pasteur Multi-Locus Sequencing Types (MLST scheme)(**Supplemental Table 2**). We chose this approach given that our strains were under strong evolutionary pressure from antibiotic treatments in the clinic. Core genome MLST sequencing, which utilizes alleles universally shared in the bacterial species, has been shown to be more robust than phylogenetic trees built using genomic sequences in bacterial populations affected by high numbers of small DNA segment homologous recombination events and presence/absence of accessory genes, both of which would apply in competitive evolution scenarios [39]. Using this approach, we observed that the bacterial isolates we tested had various resistance phenotypes for both phages and antibiotics. (**Fig. 1**)

**Fig. 1.**
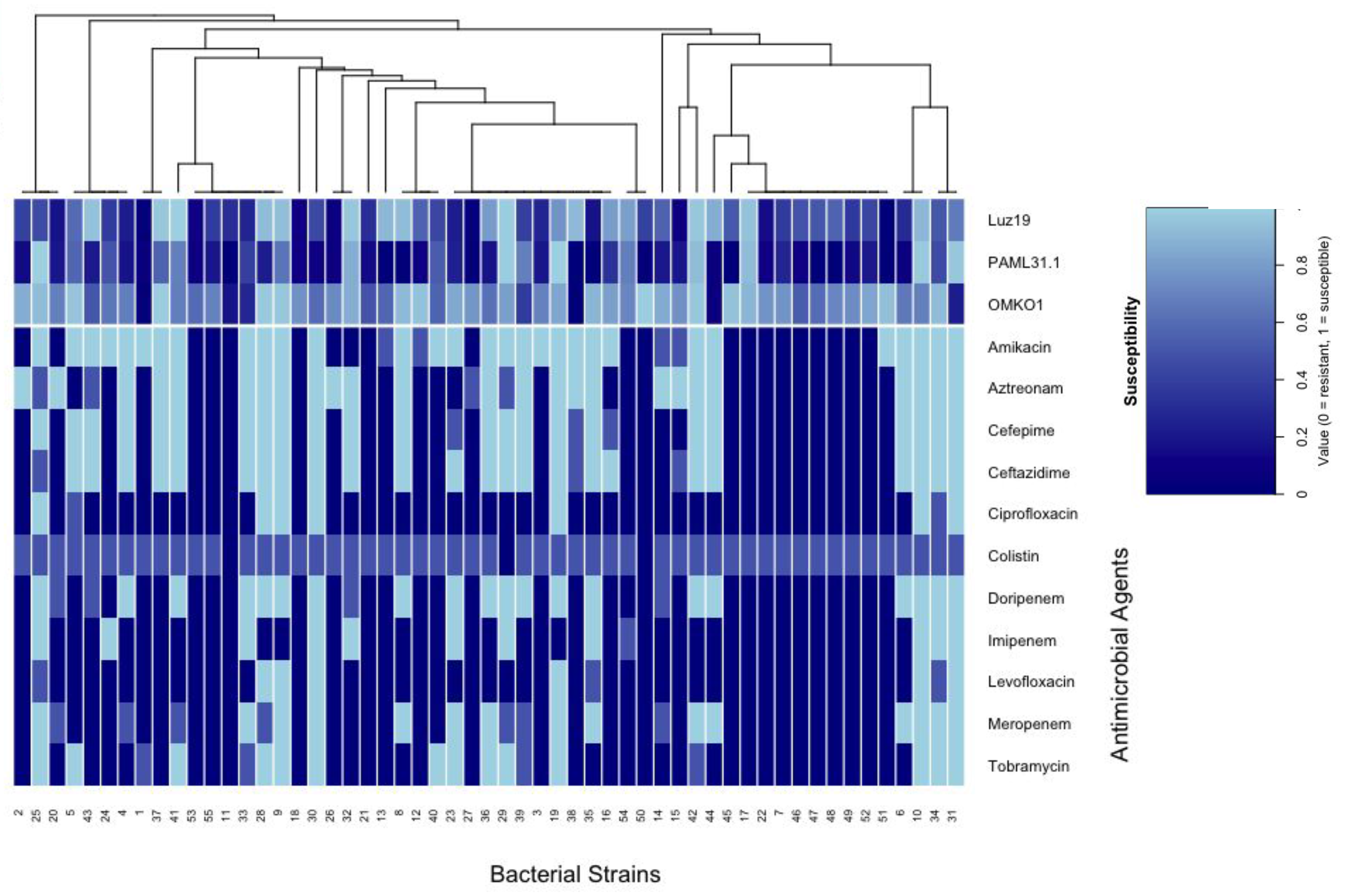
Phage and antibiotic susceptibility patterns in a set of 55 clinical isolates of *P. aeruginosa*. Phage and antibiotic susceptibility, scaled from 0 (least susceptible, dark blue) to 1 (most susceptible, light blue). Strains (x axis) are clustered by phylogeny and antimicrobials are grouped by type (phages at the top, antibiotics at the bottom). Brackets above the heatmap indicate the overall phylogenetic tree constructed based on differences in MLST allele profiles. Full OD @ 600 nm readings can be found at **Supplemental Table 3.** Growth curves can be found at **Supplemental Figure 1**

The prevalence of antibiotic resistance phenotypes was unequally distributed within this group, with 56.7% of all isolate-antibiotic interactions showing antibiotic resistance, 14.4% showing intermediate antibiotic resistance, and 28.9% showing antibiotic susceptibility (ꭓ² = 167.78, P < 0.001)**(Fig. 2a**).

**Fig. 2.**
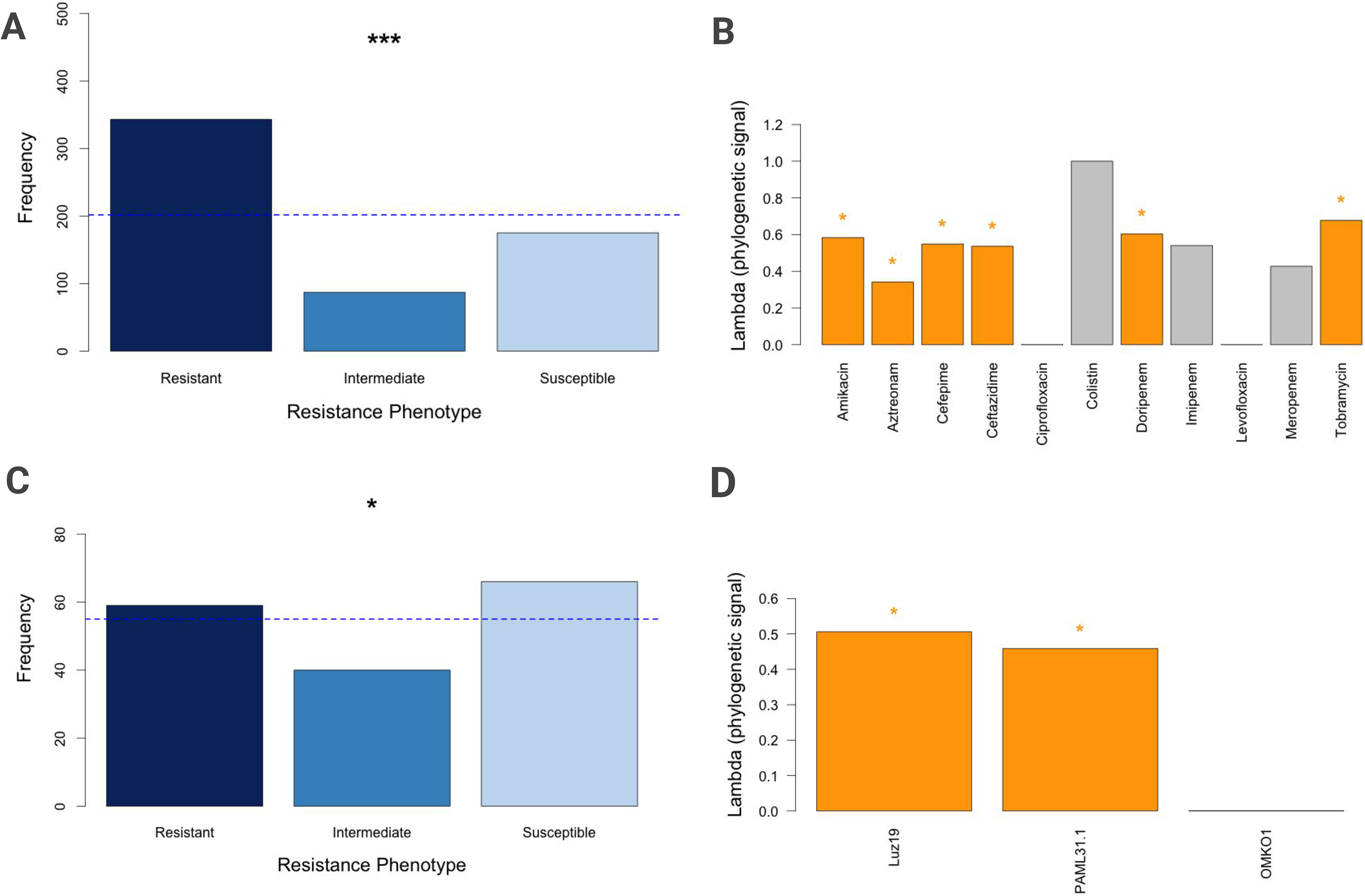
Phage and antibiotic resistance phenotype distributions and phylogenetic associations in a set of 55 clinical isolates of *P. aeruginosa*. **A)** Overall antibiotic resistance distribution, pooling interactions between all 55 strains and all 11 antibiotics. Categories determined by MIC values and CLSI 2025 standard clinical breakpoints. **B)** Association between antibiotic resistance profiles and phylogenetic distance, measured by Pagel’s lambda scores of each individual antibiotic. **C)** Overall phage resistance distribution, pooling interactions between all 55 strains and all 3 phages. Unequal distribution measured by Chi square test. Categories determined by suppression index value (x): x < 0.33, resistant; 0.33 < x < 0.66, intermediate; 0.66 < x, susceptible. **D)** Association between phage resistance profiles and phylogenetic distance, measured by Pagel’s lambda scores of each individual phage. * p<0.05; ** p<0.01; *** p < 0.001. Exact P values, Confidence Intervals, and Q values can be found at **Supplemental Tables 4 and 5**

We further observed that antibiotic resistance phenotypes were unequally distributed depending on phylogenetic similarity, with more related antibiotics displaying more similar resistance profiles. Pagel’s Lambda test showed that for 6 out of the 11 individual antibiotics, including amikacin, aztreonam, cefepime, ceftazidime, doripenem, and tobramycin, resistance profiles were significantly phylogenetically conserved across the panel of isolates ( λ = 0.34-0.68, P < 0.05)(**Fig. 2b**). For another 3 antibiotics, including colistin, imipenem, and meropenem, Lambda values showed strong phylogenetic conservation but were not significant (λ = 0.43-0.99, P > 0.05). The remaining 2 antibiotics, ciprofloxacin and levofloxacin, did not show a notable phylogenetic signal. When measuring resistance as an average across all antibiotics rather than examining the phylogenetic association of individual antibiotics, there was a significant, strong phylogenetic signal (λ = 0.473, P < 0.001). .

The prevalence of phage resistance profiles was also unequally distributed across our panel of 55 clinical isolates, with 35.8% of all isolate-phage interactions showing phage resistance, 24.2% showing intermediate phage resistance, and 40% showing susceptibility to phages (ꭓ² = 6.58, P < 0.05)(**Fig. 2c**). These categories were determined by splitting suppression index data into thirds by value, since this data is continuous and represents the complete phenotypic distribution.

We also observed that for the most part, phage resistance phenotypes were unequally distributed based on phylogenetic similarity. Pagel’s Lambda test showed that for 2 out of the 3 individual phages (Luz19 and PAML31-1), resistance profiles were significantly phylogenetically conserved across the panel of isolates (Pagel’s λ = 0.459-0.506, P < 0.005)(**Fig. 2d**). When measuring resistance as an average across all phages, there was a significant, strong phylogenetic signal (λ = 0.479, P < 0.01).

Together, these data indicate that for any given pair of strains in our clinical isolate panel, having a shared bacterial lineage increases the probability of having similar resistance levels to specific phages and antibiotics. This provides justification for controlling for phylogeny in downstream association tests by providing evidence that associations between resistance profiles might in part be due to shared phylogenetic lineage rather than acquired resistance.

### Many phage-antibiotic pairs show significant positive susceptibility associations

For phage-antibiotic pairs, 16 out of 33 pairs were statistically significant by a partial Kendall’s tau association test, and showed weak to moderate associations (τ = 0.2-0.45). The strongest phage-antibiotic pairing was PAML31-1 paired with tobramycin (τ = 0.45, p < 0.05), with Luz19’s cephalosporin pairings close behind (cefepime τ = 0.40, p < 0.05; ceftazidime τ = 0.38, p < 0.05). Notably, the phage OMKO1 and the antibiotic colistin showed no statistically significant associations with either antimicrobial type (**Fig. 3a**). There were no significant negative associations.

**Fig. 3.**
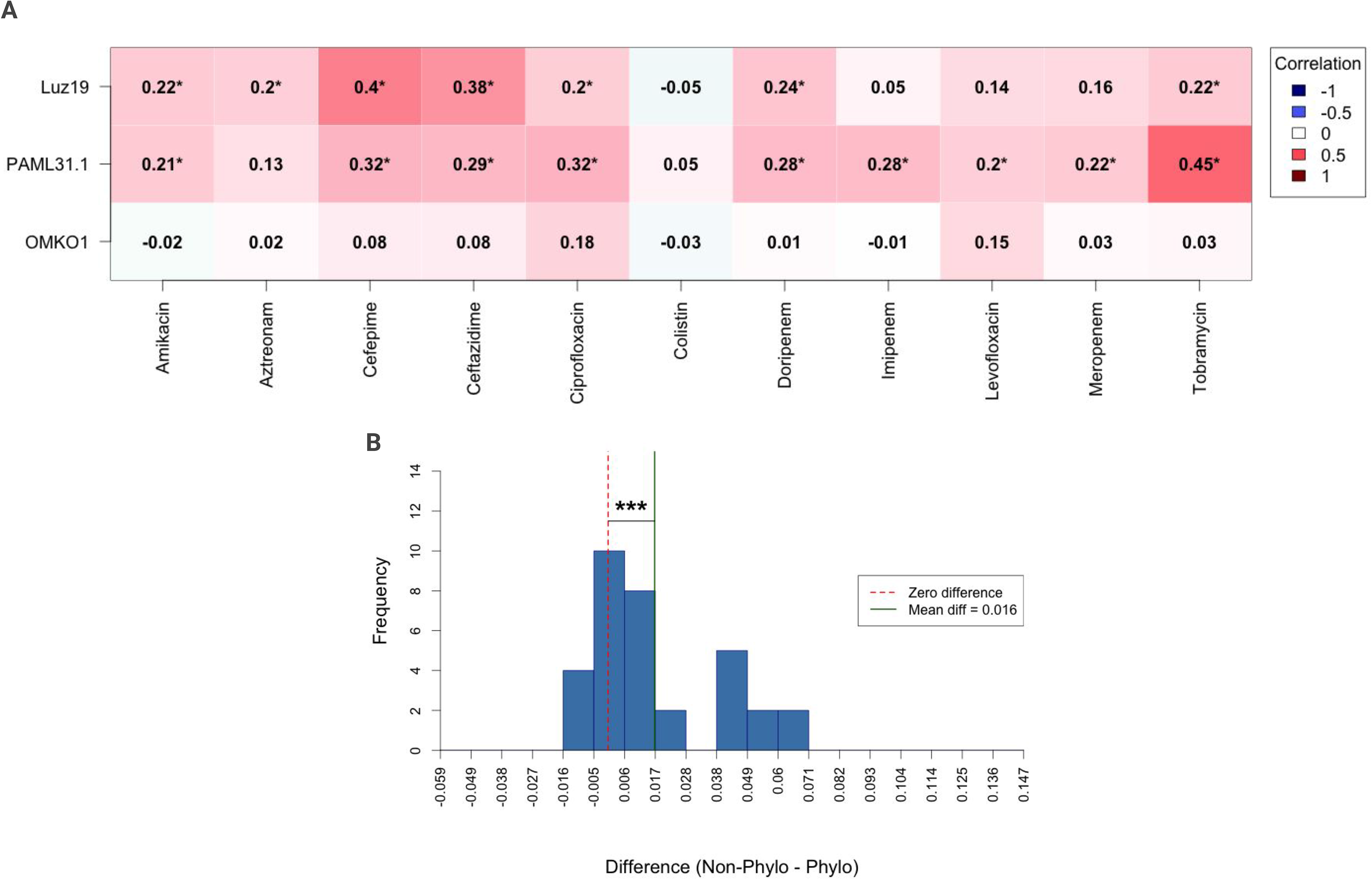
Phage-antibiotic pairwise susceptibility associations. **A)** A phylogenetically-controlled (partial) Kendall’s Tau correlation matrix for bacterial susceptibility profiles to the 11 antibiotics and phages in our data, displaying strength of association for phage-antibiotic pairings. Association strength is displayed by color, with dark red corresponding to strongest positive association and dark blue corresponding to strongest negative association. **B)** Histogram showing the distribution of the differences between uncorrected and phylogenetically corrected association values for phage-antibiotic pairings. For **A)** and **B)** * p<0.05; ** p<0.01; *** p<0.001 The raw data values for these data are available as **Supplemental Table 1**. The complete phylogenetically corrected correlation values are available as **Supplemental Table 6.** The complete raw (uncorrected) correlation values are available as **Supplemental Table 7**.

We then asked whether bacterial lineage effects could explain these associations. To this end, we compared the distribution of phylogenetically corrected pairwise correlations to uncorrected pairwise correlations for phage-antibiotic pairs. We found that uncorrected pairwise associations were systematically higher than phylogenetically corrected pairwise associations, indicating again that raw associations are amplified by phylogenetic signal but this effect can be removed by controlling for phylogeny (**Fig. 3b**).

Together, these data demonstrate that for phage-antibiotic pairings, susceptibility to one antimicrobial agent sometimes predicts susceptibility to another antimicrobial agent. The manner of these associations was only partially dependent on phylogeny, potentially indicating a mechanistic tie between resistance to certain phages and antibiotics. This unexplained mechanism driving associations between specific phage-antibiotic pairings warrants further investigation.

### There are few associations between phage-phage pairs

We next examined whether susceptibility to individual phages was associated with susceptibility to other individual phages. For phage-phage pairs, only one pairing (Luz19 x PAML31-1) was moderately associated (τ = 0.31, p < 0.05). We did not observe other significant phage-phage susceptibility associations. There were no negative associations (**Fig. 4a**).

**Fig. 4.**
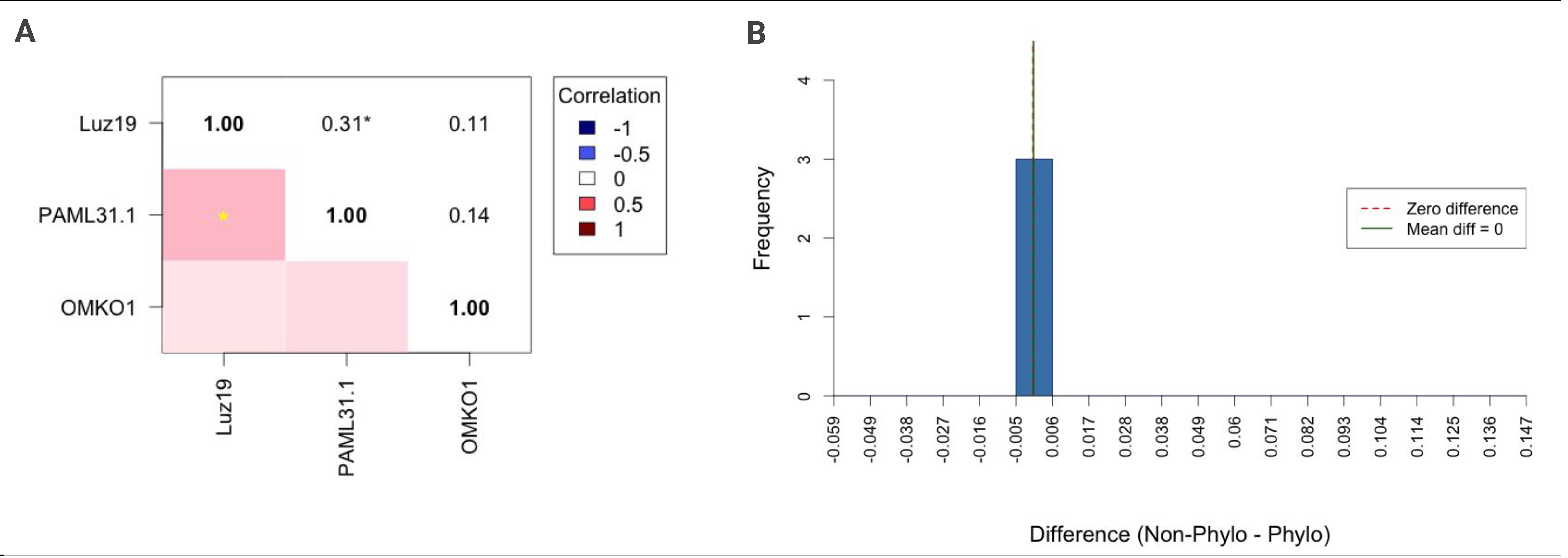
Phage-phage pairwise susceptibility associations. **A)** A phylogenetically-controlled (partial) Kendall’s Tau correlation matrix for bacterial susceptibility profiles to the three phages in our data, displaying strength of association for phage-phage pairings. Association strength is displayed by color, with dark red corresponding to strongest positive association and dark blue corresponding to strongest negative association. The upper triangle of the heatmap displays the association values while the lower triangle displays the corresponding colors. **B)** Histogram showing the distribution of the differences between uncorrected and phylogenetically corrected association values for phage-phage pairings. For **A)** and **B)** * p<0.05; ** p<0.01; *** p<0.001 The raw data values for these data are available as **Supplementary Table 1**. The complete phylogenetically corrected correlation values are available as **Supplemental Table 6**. The complete raw (uncorrected) correlation values are available as **Supplemental Table 7**.

There was no significant difference between phylogenetically corrected and uncorrected phage-phage associations, indicating that the raw association was not amplified by phylogenetic signal. This absence of a bacterial lineage effect to explain the phage-phage association could potentially indicate shared mechanisms of resistance that were evolved independently (**Fig. 4b**).

Together, these data demonstrate that susceptibilities to the different phages in our study were not strongly associated. The one moderate association that did occur contained no significant element of phylogenetic signal, potentially indicating independent evolution of resistance to the phages in this dataset.

### Many antibiotic-antibiotic pairs show significant positive susceptibility associations

For antibiotic-antibiotic pairs, 44 out of 55 pairings were statistically significant by Kendall’s tau partial association test, displaying a range of association strength (τ = 0.2-0.93, p < 0.05)(**Fig. 5a).** Susceptibility to imipenem and colistin was modestly negatively associated (τ = -0.1).

**Fig. 5.**
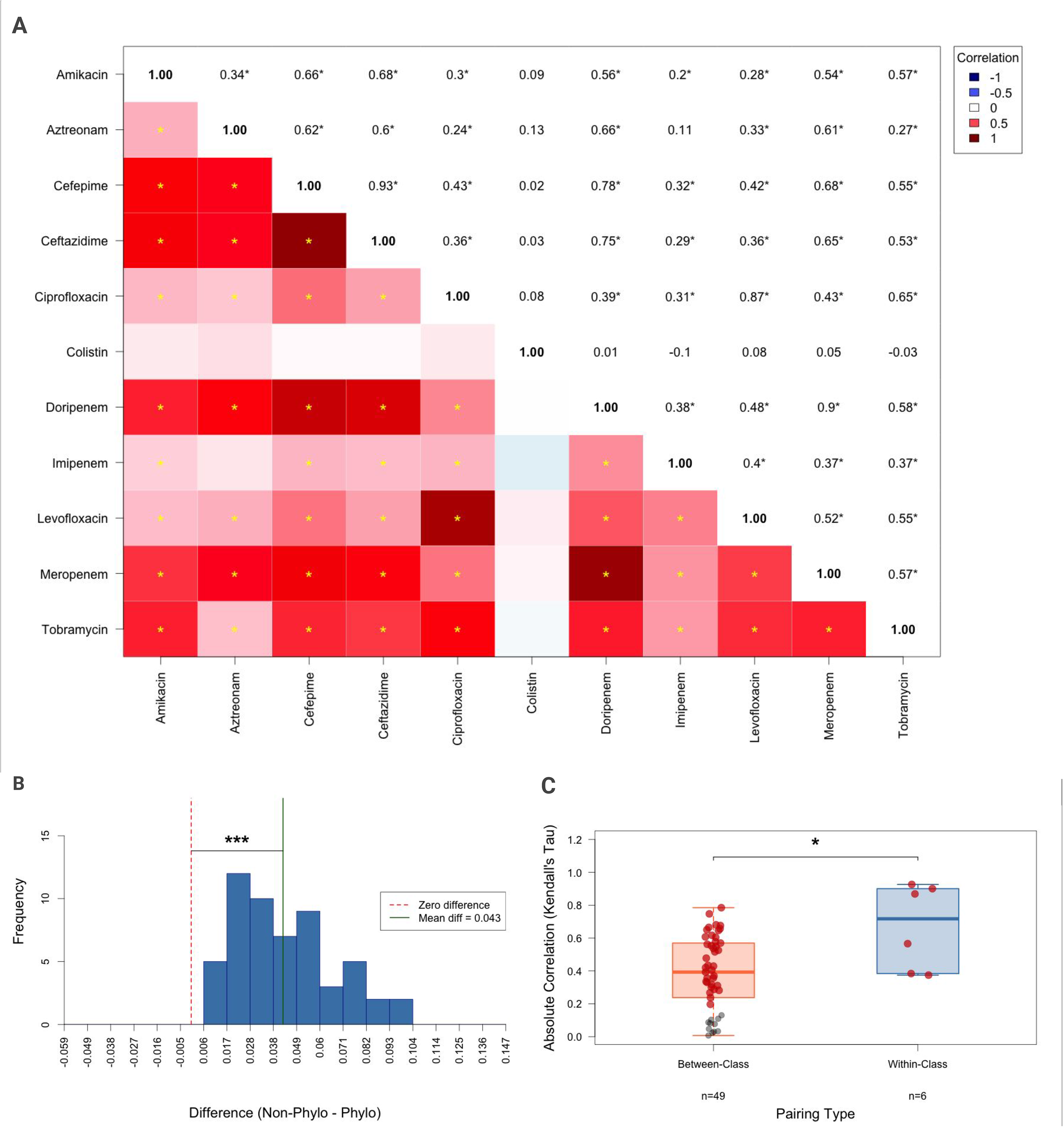
Antibiotic-antibiotic pairwise susceptibility associations. **A)** A phylogenetically-controlled (partial) Kendall’s Tau correlation matrix for bacterial susceptibility profiles to the 11 antibiotics in our data, displaying strength of association for antibiotic-antibiotic pairings. Association strength is displayed by color, with dark red corresponding to strongest positive association and dark blue corresponding to strongest negative association. The upper triangle of the heatmap displays the association values while the lower triangle displays the corresponding colors. **B)** Histogram showing the distribution of the differences between uncorrected and phylogenetically corrected association values for antibiotic-antibiotic pairings. **C)** Strength of antibiotic-antibiotic associations within-antibiotic class versus between-antibiotic class. Each dot represents a particular pairing, mapped by its absolute correlation as determined by the partial Kendall’s tau test. Red dots indicate p<0.05. For **A)**, **B)**, and **C)** * p<0.05; ** p<0.01; *** p<0.001 The raw data values for these data are available as **Supplemental Table 1**. The complete phylogenetically corrected correlation values are available as **Supplemental Table 6.** The complete raw (uncorrected) correlation values are available as **Supplemental Table 7**.

We then asked whether bacterial lineage effects could explain these associations. To this end we compared the distribution of phylogenetically corrected pairwise correlations to uncorrected pairwise correlations for antibiotic-antibiotic pairs. We found that uncorrected pairwise associations were systematically higher than phylogenetically corrected pairwise associations, indicating that raw associations are amplified by phylogenetic signal, an effect that can be removed via phylogenetic control (Wilcoxon p < 0.001)(**Fig. 5b**). Individual associations that remain significant despite phylogenetic control might suggest a meaningful link to known mechanisms of antibiotic cross-resistance, such as pleiotropic mutations or co-inheritance of resistance genes via mobile genetic elements [40–41]. This link is supported by within-class antibiotic associations varying significantly from between-class associations (Mann-Whitney U W = 63, p < 0.05), with within-class associations (n = 6, mean = 0.67) being stronger than between-class (n = 49, mean = 0.38)(**Fig. 5c**).

Together, these data indicate that for pairs of antibiotics, antibiotic susceptibility patterns predicted susceptibility to other antibiotics in a manner only partially dependent on phylogeny. The strength of this effect was also dependent on the relation between the classes of the two antibiotics.

### Comparing antimicrobial types reveals stronger phylogenetic signal for antibiotic-antibiotic pairings than phage-antibiotic pairings

Next, we asked whether resistance phenotypes were more strongly associated on average within or between the two types of antimicrobial agents. We compared the distributions of association strength across antibiotic-antibiotic (n=55, mean=0.418), phage-phage (n=3, mean=0.189), and antibiotic-phage (n=33, mean=0.171) pairing groups. Kruskal-Wallis analysis revealed significant differences across pairing types (*χ*² (2) = 47.158, p < 0.001), and post-hoc analysis showed that antibiotic-antibiotic correlations were statistically stronger than phage-phage correlations (p < 0.05)(**Fig. 6a**).

**Fig. 6.**
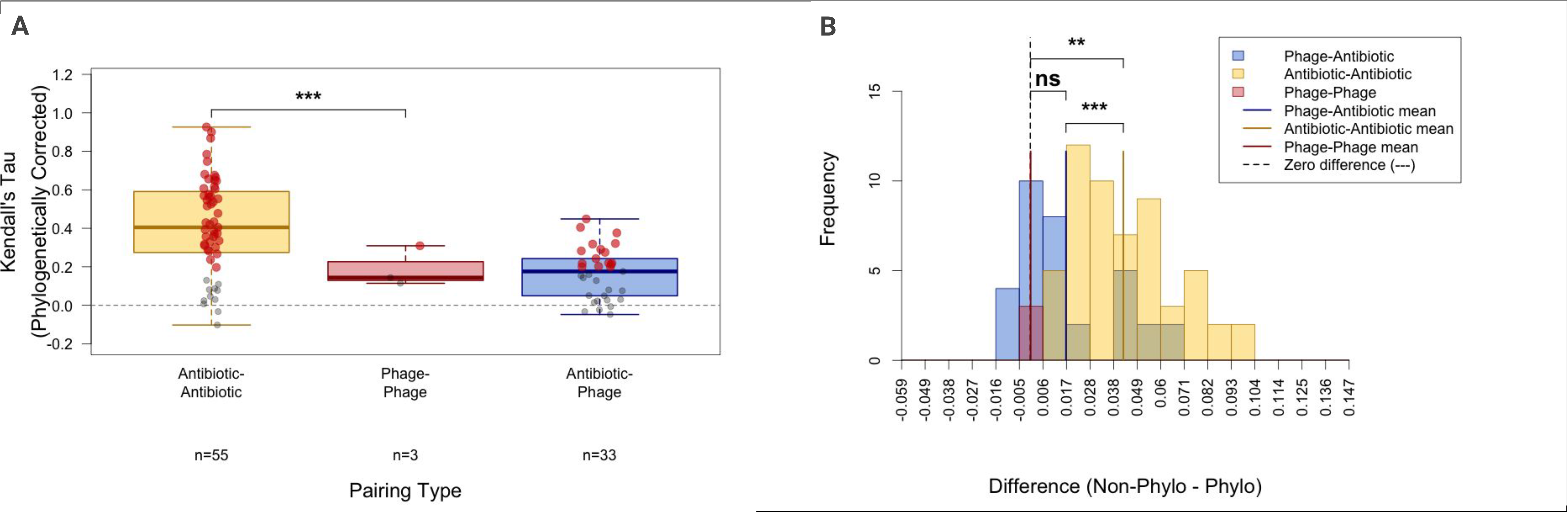
Comparing Association Strength and Bacterial Lineage Effect by Antimicrobial Combination. **A)** Strength of association by pairings amongst different categories of antimicrobials (antibiotic-antibiotic, phage-phage, phage-antibiotic). Each dot represents a particular pairing, mapped by its absolute correlation as determined by the partial Kendall’s tau test. Red dots indicate p<0.05. B) Histogram comparing the distributions of differences between uncorrected and phylogenetically corrected association values for pairings amongst different categories of antimicrobials (antibiotic-antibiotic, phage-phage, phage-antibiotic). For **A)** and **B)** * p<0.05; ** p<0.01; *** p<0.001 The raw data values for these data are available as **Supplementary Table 1**. The complete phylogenetically corrected correlation values are available as **Supplemental Table 6.** The complete raw (uncorrected) correlation values are available as **Supplemental Table 7**.

We then examined the differences between the size of bacterial lineage effects across pairing types. While examination of bacterial lineage effects within individual pairings showed both phage-antibiotic and antibiotic-antibiotic pairing types to be significant, this cross-pairing type comparison showed the bacterial lineage effect to be larger for antibiotic-antibiotic pairings than phage-antibiotic pairings (**Fig. 6b**).

Together, these data suggest a largely orthogonal development of resistance to the diverse phages in our panel. On the other hand, there is a strong phylogenetic signal that drives similar resistance profiles to pairs of antibiotics among different strains in our panel. While there were also numerous cases in which many strains displayed similar resistance profiles to certain pairs of phages and antibiotics (with no significant difference between the mean strength of association for antibiotic-antibiotic and antibiotic-phage pairings), these associations were not explained by shared phylogenetic lineage to the same extent as the associations between two antibiotics.

### Nonrandom positive association between overall patterns of phage and antibiotic resistance

Having examined individual phage-antibiotic pairings in Fig. 1, we next asked whether the overall frequency of bacterial susceptibility to both phages and antibiotics was correlated in this sample set. We found a moderately positive overall association between average phage and antibiotic resistance for each strain when controlling for phylogeny using the MLST scheme. A partial Spearman test accounting for 49% of phylogenetic variance using PC1 showed the strongest positive association (Partial Spearman r = 0.358, P < 0.05), while a partial Mantel test and a phylogenetic least squares (PGLS) test, both accounting for 100% of the phylogenetic structure, remain significantly positive but weaker (partial Mantel r = 0.18, P = 0.001; PGLS β = 0.22, P < 0.05) (**Fig. 7a**) (**Supplemental Table 8**).

**Fig. 7.**
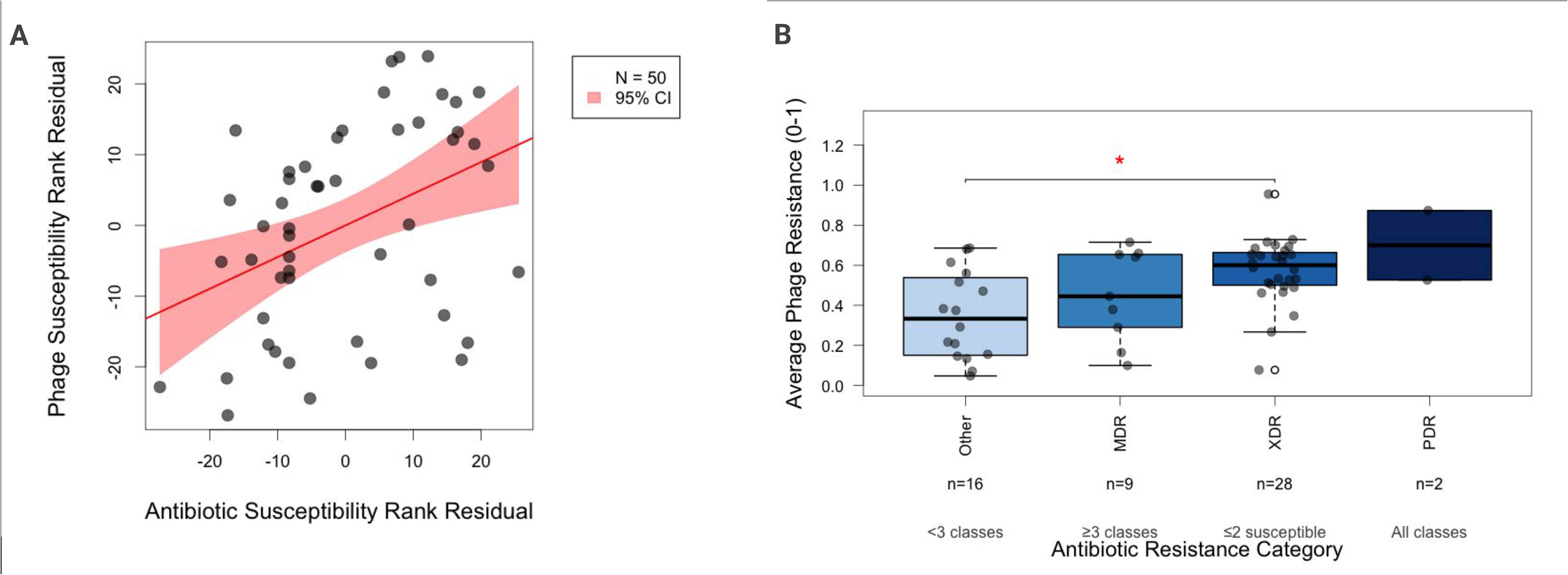
Association Between Overall Patterns of Phage and Antibiotic Resistance in these samples. **A)** Strains mapped by susceptibility residuals obtained by regressing average resistance across all 11 antibiotics and average resistance across all 3 phages on PC1 from the phylogenetic distance matrix calculated based on the MLST allele scheme of each strain. The trend line seen in red was calculated by the ordinary least squares method, and 2 outliers identified via the standardized residual method were removed. The parametric 95% confidence interval (CI) of this regression is shaded in red. Various overall antibiotic and phage resistance correlation test results and their P-values and CIs are reported in **Supplemental Table 8**. **B)** Average phage resistance compared between strains grouped by antibiotic resistance levels determined by the number of classes of antibiotics the strain in question is resistant to. For **A)** and **B)**, each dot corresponds to one strain. Average antibiotic resistance was calculated using the scaled susceptibility values based on MIC data. Average phage resistance was calculated using suppression assay data.

When grouping strains by the clinically relevant MDR classification [15], which is determined by the number of antibiotic classes the strains are resistant to, a similar pattern was observed. Kruskal-Wallis analysis showed a significant difference between the means of the categories “Other” (not classified in the MDR scale; resistant to <3 antibiotic classes), MDR (multidrug-resistant; resistant to ≥ 3 antibiotic classes), XDR (extensive drug resistance; susceptible to ≤ 2 antibiotic classes), and PDR (pandrug-resistant; resistant to all antibiotic classes in the panel), and the mean phage resistance increased with each category, respectively (Kruskall-Wallis *χ*² (3) = 10.489, p < 0.05). Post-hoc analysis with multiple hypothesis testing correction showed a significant difference between the XDR and Other groups (**Figure 7c**).

Together, these data indicate that in a panel of *P. aeruginosa* strains isolated from the clinic, resistance to antibiotics and phages tends to increase in tandem.

### Presence of resistance genes and isolate source alone do not explain patterns of cross-resistance

In order to gather more information about the demonstrated co-resistance to phages and antibiotics in this panel of clinical strains, we analyzed how certain properties of the individual strains influenced both individual pairwise associations and overall patterns of phage and antibiotic resistance.

First, we asked whether certain clinical strain sources displayed particular patterns of phage and antibiotic resistance. Isolates sourced from aspirate were on average significantly more susceptible to antibiotics than the average across all 55 strains (p < 0.05), and their average antibiotic susceptibility also varied significantly from their average phage susceptibility, with a mean antibiotic susceptibility of 0.73 and a mean phage susceptibility of 0.43 (p < 0.05). Bronchial washings, urine, and strains without a source listed were on average more susceptible to our phages than they were to antibiotics (bronchial washings and urine: p < 0.005; unknown source: p < 0.05)(**Fig. 8a**, **Fig. 8b**).

**Fig. 8.**
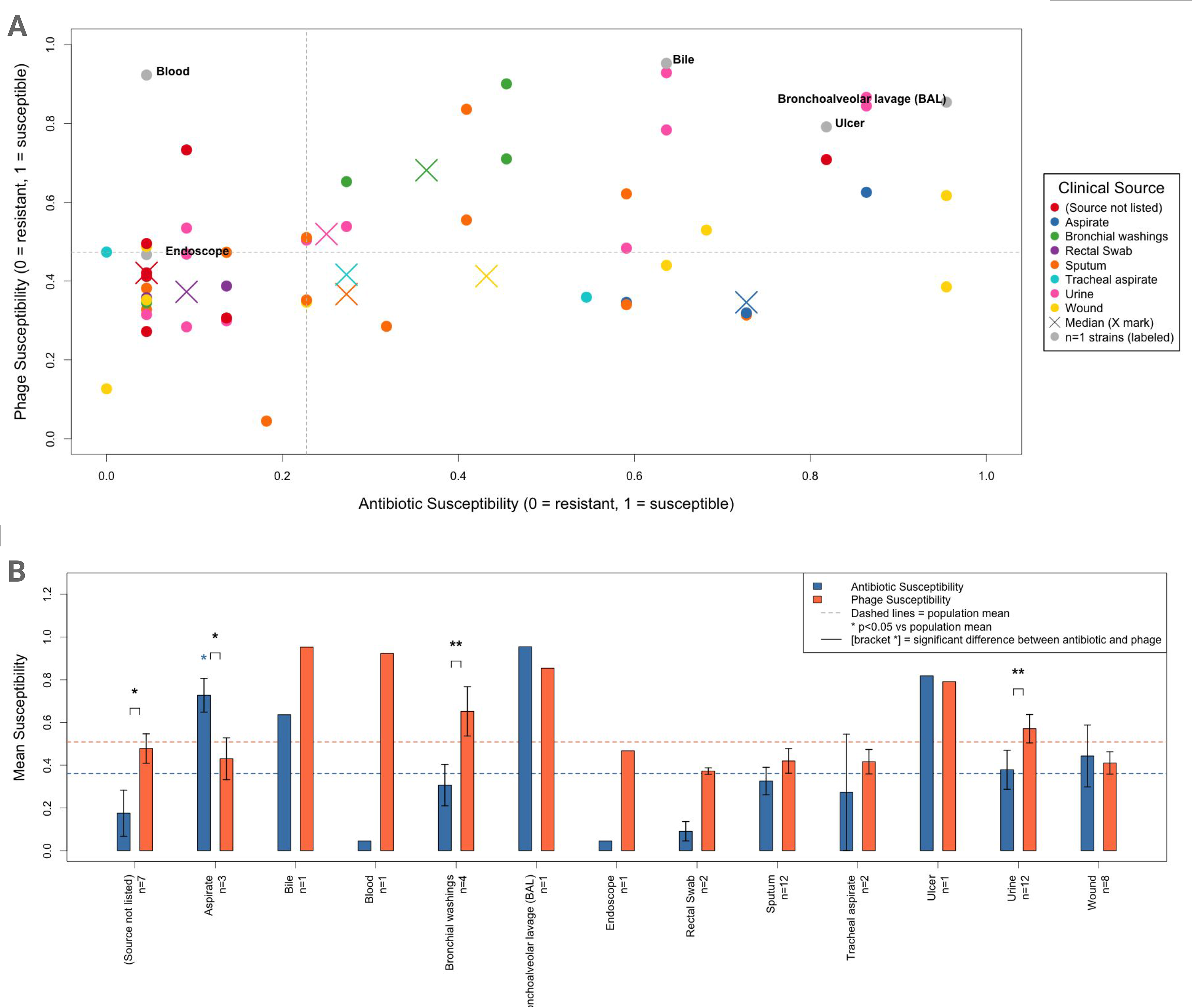
Susceptibility Patterns by Clinical Source. A) Resistance profiles by clinical source. Dots indicate individual strains, exes indicate the mean phage and antibiotic susceptibility within a source group. Grey dots indicate source groups with n=1 strains, labeled by source. B) Variance from the population means for phage and antibiotic resistance. Blue asterisks indicate significant (p < 0.05) variance from the population mean for a particular antimicrobial agent, brackets indicate a significant (* p < 0.05; ** p < 0.01) difference between average phage and antibiotic resistance.

Next, we tested for enrichment or depletion of previously characterized resistance genes in phage-resistant strains among strains resistant to antibiotics to see whether these genes might control instances of cross-resistance. The CDC performed whole genome sequencing and provided the known resistance genes carried by individual strains, which we used for this analysis. We defined phage-resistant strains by a suppression index value in the lower third (<0.33), and only tested phage-antibiotic pairings that were significantly associated via Kendall’s Tau. After controlling for phylogeny by fitting a logistic mixed model using phylogenetic PC1 and PC2 as well as an intercept for lineage, no genes remained significant (**Supplemental Table 9**). We then controlled for phylogeny instead by comparing gene presence only among groups of strains within each lineage, which also yielded no significant results.

Together, these data indicate that the phage-antibiotic co-resistance in this dataset cannot be explained by source of infection or presence of previously characterized resistance genes alone. This indicates that further examination of the factors driving cross-resistance, genetic and otherwise, is necessary in future studies.

## Discussion

We report that resistance to phages and antibiotics was frequently associated across this collection of clinical *P. aeruginosa* isolates, with numerous strains displaying similar resistance profiles to particular phage-antibiotic pairs. More broadly, we observed a positive association between aggregate phage resistance and antibiotic resistance across the strain collection: isolates that were more resistant to antibiotics were, on average, also more resistant to the phages tested. This pattern was also apparent when isolates were stratified by conventional antibiotic resistance categories (non-MDR, MDR, XDR, and PDR), with mean phage resistance increasing with increasing multidrug resistance. These findings suggest that the highly antibiotic-resistant clinical isolates for which alternative antimicrobial approaches may be most needed may also, on average, be less susceptible to available phages. Our results therefore caution against assuming that phage and antibiotic susceptibility are necessarily independent or orthogonal properties of clinical isolates.

Genetic relatedness contributed to, but did not fully account for, these associations. Comparison of unadjusted associations with those controlling for phylogeny revealed the largest contribution of shared lineage to antibiotic-antibiotic associations, a more modest contribution to phage-antibiotic associations, and little apparent contribution to phage-phage associations. Among phylogeny-controlled antibiotic-antibiotic comparisons, within-class associations were stronger than between-class associations. This provides internal validation of the analytical approach, as antibiotics within the same class frequently share mechanisms of action and corresponding bacterial resistance mechanisms [42]. Importantly, the persistence of an overall association between phage and antibiotic resistance after controlling for phylogeny suggests that shared ancestry alone does not explain the observed relationship.

The biological basis for this association remains unclear. Phages and antibiotics generally target very different bacterial structures and processes, and our data do not establish that common resistance mechanisms directly confer resistance to both. Instead, the observed association could arise from multiple mechanisms. Bacterial traits associated with multidrug resistance—including alterations in cell-surface structures, envelope composition, efflux and global regulatory pathways, growth state, biofilm-associated phenotypes, or the accumulation of anti-phage defense systems—could independently influence susceptibility to both classes of antimicrobial agents. Alternatively, repeated antimicrobial exposure or other ecological pressures in clinical settings could select for combinations of traits that favor resistance to both antibiotics and phages. Distinguishing among these possibilities will require mechanistic investigation.

Our study has several limitations. First, it is exploratory and observational and therefore cannot establish causal mechanisms underlying the observed associations. Phages and antibiotics were tested independently, so these data also cannot predict synergy, antagonism, or efficacy when particular phage-antibiotic combinations are administered simultaneously. Second, our analysis included only three phages. Although these phages are well characterized and differ in morphology and receptor usage, they cannot capture the diversity of phage-host interactions found across the broader *P. aeruginosa* phageome. Likewise, the 55 isolates in this study represent a useful and phylogenetically diverse clinical collection but may not reflect the full diversity of *P. aeruginosa* encountered across different infection types, geographic regions, or patient populations. Finally, we did not experimentally identify the genetic or physiological mechanisms responsible for the observed associations.

Future studies should therefore determine which bacterial traits account for the association between phage and antibiotic resistance. This could be done via a genome-wide association study (GWAS) on particular clinical and lab-generated strains which display high levels of resistance to phages and antibiotics, using a method similar to what has been previously done to characterize antibiotic co-resistance [43–45]. It will be important to recapitulate co-resistance using experimental evolution, and validate causal genetic elements flagged by GWAS via genetic knock-outs and knock-ins. .

Comprehensive testing has been done to study co-resistance between pairs of antibiotics, but our study sets the stage for further research into patterns and mechanisms of co-resistance between pairs of phages and antibiotics [46–48]. Research in this area is essential to better understand and predict the efficacy of therapeutic options using phages and/or antibiotics to resolve persistent clinical infections.

In summary, we find that phage and antibiotic resistance are positively associated across a phylogenetically diverse collection of clinical *P. aeruginosa* isolates and that this relationship persists after accounting for bacterial phylogeny. These results challenge the assumption that susceptibility to phages and antibiotics is necessarily independent and raise the question of which bacterial traits underlie their association. Understanding these relationships may improve both our understanding of antimicrobial resistance biology and the rational development of phage-based therapies for multidrug-resistant infections.

## Methods

### Bacteria used in this study

The PSA (*P. aeruginosa*) panel of 55 diverse clinical isolate strains was obtained from the CDC Antimicrobial Resistance Isolate Bank [49] (**Table 1**). Full genomes for each strain are available at Biosample Accession Numbers SAMN04901619 through SAMN04901662, SAMN15646438, SAMN15646440, SAMN15646443 through SAMN15646445, SAMN11954021, and SAMN11954023 through SAMN11954027.

MIC results for each of the 11 antibiotics against each strain were obtained by the CDC by broth microdilution. Clinically relevant classifications of “Resistant,” “Intermediate,” and “Susceptible” based on these MICs for particular antibiotics were assigned one of 3 numeric values–0 for resistant, 0.5 for intermediate, and 1 for susceptible. Whenever reported as “resistance” rather than “susceptibility” (in figures or for data analysis purposes), these values were subtracted from 1 (1 for resistant, 0.5 for intermediate, 0 for susceptible).

### Phages used in this study

As phages in this study, we selected three phages which each target distinct *P. aeruginosa* receptors, and are thus members of three different ‘receptor complementarity groups’, as we previously described in Kim *et al.*, 2024 [11, 31–33]. These phages represent different phage phylogenies and have different sized genomes (**Table 2**).

### Bacterial Cultures

Cultures of bacteria were grown as follows. Bacteria was streaked on agar plates and incubated overnight (12-16 hours) at 37°C. Approximately 6 colonies were then inoculated in 3 mL of LB Miller broth and incubated at 37°C with constant shaking at 250 rpm to an OD600 reading of 0.4-0.8 (3-4 hours).

### Phage Production and Purification

Phage purification was performed as previously reported [50–51]. In brief, the titre of phage lysates was determined by spotting 10-fold serial dilutions of phage lysate up to a dilution factor of 1e10 on solidified top agar mixed with bacterial inoculum. After incubation overnight at 37°C, PFU/mL was calculated for the phage batch based on plaque counts.

Appropriate dilutions to ensure substantially separated plaques were mixed with molten top agar and bacterial inoculum, then plated and let dry before incubating overnight (12-16 hours). Plaques were selected based on morphology and separation. These plaques were picked and expelled into an eppendorf tube containing 1 mL of SM buffer (100 mM NaCl, 8 mM MgSO4·7H2O, 50 mM Tris-Cl (pH 7.5), and 0.01% (w/v) gelatin in H2O). These solutions were then incubated at room temperature for 10 minutes before being diluted and plated as described above. The process was repeated three times, or until plaques showed completely uniform and expected morphologies.

Phage propagation was performed as follows. Phage lysate was added to host culture at a final concentration of 3e5 PFU/mL, followed by incubation at 37°C with constant shaking at 250 rpm for 2 hours. The entirety of the contents of this culture was then added to prewarmed salt-supplemented LB media (10 mM MgSO₄, 10 mM CaCl₂) and incubated overnight (12-16 hours). The resulting lysate was syringe-filtered using PES 0.2-mm filters and stored at 4°C.

Phage identity and purity was confirmed by whole genome sequence analysis. Filtered phage lysate was treated with DNAase (2 U/mL), RNAse A (10 µg/mL), and proteinase K (30 µg/mL) (New England Biolabs, Ipswich, MA, USA) and extracted using a QIAGEN DNeasy kit (Qiagen, Germantown, MD, USA). Standard Microbial WGS libraries were prepared by Novogene and sequenced on an Illumina NovaSeq X Plus Series (PE150) (Novogene, Sacramento, CA, USA). Genome assembly and annotation were done according to Shen and Millard (2021) [52].

To further ensure purity of the desired phage, all lytic phage propagations and purifications were performed using a mutant *P. aeruginosa* PAO1 strain with prophage pf4 and pf6 deletions (mPAO1Δpf4Δpf6) as the host (Schmidt, *et. al*, 2024 [53]).

### Phage Suppression Assay

The phage suppression assay was adapted from the protocol used by Kim *et. al*, 2024 [11]. Host cultures were grown overnight, then diluted 1:200 in LB and regrown for 2.5 hours (to OD600 0.1-0.2), then diluted again 1:100 (1e6 CFU/mL confirmed by CFU counts). Phage stocks were prepared at the appropriate dilutions to reach multiplicities of infection (MOIs) of 1 and 100 (1e6 and 1e8 PFU/mL), and 75 uL were added to each well of a 96-well plate, excluding growth control wells and sterility controls/blanks, with each of the 3 phages occupying 2 rows (1 for each MOI). 75 uL bacteria were then added to each well, with 1 strain in each column (12 strains per plate). The plates were covered with polyurethane Breath-Easy® sealing membranes (Sigma-Aldrich, St. Louis, MO, USA) then incubated with constant shaking at 37°C for 20 hours in an automated spectrophotometer (Biotek Synergy2 plate reader), during which time OD at 600 nm was measured every 20 minutes. Area under the curve (AUC) for resultant growth curves was calculated using Graphpad Prism Version 9.0 software, and growth curves were generated using R (**Supplemental Figure 1**). All of this was performed for 3 biological replicates. The suppression index was calculated using the following formula:

(*AUC growt*ℎ *control condition* − *AUC p*ℎ*age treated condition*) / *AUC growt*ℎ *control condition*

Whenever this suppression index metric was reported as “resistance” rather than “susceptibility” (in figures or for data analysis purposes), these values were subtracted from 1.

### Statistical Analysis

All statistical analyses were performed using R. The necessity of controlling for phylogenetic distance when comparing phenotypic relationships via regression or correlation is explained by Felsenstein, 1985 [54] and supported by our demonstration of phylogenetic association between strain resistance phenotypes using Pagel’s λ test via the fitContinuous() function for continuous phage susceptibility data and the fitDiscrete() function using the equal rates (ER) model for ordinal antibiotic susceptibility data, both from the geiger R package [55]. Whenever antibiotic susceptibility data was collapsed into one average per strain, values within the same antibiotic class were averaged first before generating one average of all the classes in order to prevent classes which were overrepresented in the panel from disproportionately affecting the results. Rather than Phylogenetically Independent Contrasts (PICs), which calculate differences between two descendant lineages at each node of the phylogenetic tree [52], we use a partial correlation approach that is analogous to Phylogenetic Eigenvector Regression (PVR), which uses PCA analysis on phylogenetic distance matrices to calculate residuals used for correlation tests [56].

To obtain our phylogenetic distance matrix, we utilized allelic differences given by MLST profiles contained in the CDC ARI Bank Data [49]. Calculating phylogenetic distance using MLST schemes rather than whole-genome analysis is a traditional method [57], but the two have been shown to produce similar results [39]. After calculating the phylogenetic distance matrix, we performed a principle component analysis and used the dominant PC1 as a representation of the gradient of evolutionary divergence.

We then used the pcor.test() function from the ppcor R package [58] to perform a partial correlation by regressing both phage and antibiotic resistance data onto the phylogenetic PC1 values to remove variation associated with phylogeny and correlated the residuals to identify an independent association between antimicrobial resistance profiles. This partial correlation (multiple regression) approach has been shown to be more effective than stepwise regression with downstream testing approaches [59]. We chose Kendall’s τ as the nonparametric test for our partial pairwise correlations because of its robustness to outliers and tied ranks [60]. For our overall association test comparing average phage and antibiotic resistance across all strains, we chose the Spearman ρ test because it is the standard for Mantel tests [61] and used the same phylogenetically controlled partial correlation approach (partial Spearman ρ), using PC1 of the MLST-based distance matrix. The averaging of the ordinal data in the Mantel test compared to the pairwise associations greatly decreases ties, hence our use of Spearman in this case and Kendall in the other. Since PC1 only accounts for part of the phylogenetic variance, we then conducted a partial Mantel test in which the Spearman statistic is calculated by correlating Euclidean antibiotic-susceptibility and phage-susceptibility distance matrices while partialing out the full pairwise phylogenetic distance matrix. Finally, an alternative phylogenetic generalized least squares approach was taken. In this analysis, a hierarchical clustering tree was built from the MLST-based distance matrix, and phage susceptibility was regressed on antibiotic susceptibility using the generalized least squares method with a Pagel’s λ correlation structure fit to the tree. This was done using the gls() function from the nlme R package and the corPagel function from the ape R package. When testing for resistance gene enrichment in cross resistant strains, we used a mixed-effects logistic regression (gene presence/absence + PC1 + PC2 + (1 | clonal lineage)) fit with the bglmer() function from the blme R package per candidate gene. All other analyses were performed using base R and standard nonparametric tests. All unspecified multiple testing corrections were calculated via FDR.

## Supplemental Materials

All supplemental materials can be found at https://docs.google.com/document/d/1CJh6ljp8tXY6iZDsHE4fGxW073YjkJyjIizK_vBDflY/edit?usp=sharing

## Acknowledgments

P.L.B. discloses support from National Institutes of Health grants R01 HL148184-01; R01 AI12492093; R01 DC019965, K24 AI166718, P01AI1960471, and R01 EB038154 and support from the Cystic Fibrosis Foundation and the CFRI.

---

**Supplemental Table 1.** Data matrix displaying suppression index values for each phage against each strain, as well as scaled susceptibility values based on MIC using S/I/R breakpoints adapted from CLSI 2025 M100 S35. S/I/R converted to numerical scale by assigning S=1, I=0.5, R=0 to match directionality of the suppression index used for phage AST. AMK: amikacin; AZM: aztreonam; CPM: cefepime; CAZ: ceftazidime; CIP: ciprofloxacin; COL: colistin; DOR: doripenem; IPM: imipenem; LVX: levofloxacin; MEM: meropenem; TOB: tobramycin confounded by the lineage effect.

| strain | Luz19 | PAML31-1 | OMKO1 | AMK | AZM | CPM | CAZ | CIP | COL | DOR | IPM | LVX | MEM | TOB |
| --- | --- | --- | --- | --- | --- | --- | --- | --- | --- | --- | --- | --- | --- | --- |
| 1 | 0.03322575 | 0.06615145 | 0.034703267 | 1 | 0 | 0 | 0 | 0 | 0.5 | 0 | 0 | 0 | 0 | 0.5 |
| 2 | 0.406899905 | 0.14330975 | 0.868423802 | 0 | 1 | 0 | 0 | 0 | 0.5 | 0 | 0 | 0 | 0 | 0 |
| 3 | 0.268204978 | 0.199540241 | 0.694667436 | 1 | 0 | 0 | 0 | 0 | 0.5 | 0 | 0 | 0 | 0 | 0 |
| 4 | 0.224367874 | 0.17519317 | 0.677634695 | 1 | 1 | 1 | 1 | 0 | 0.5 | 1 | 0 | 0 | 0.5 | 0 |
| 5 | 0.59316481 | 0.576949599 | 0.95998133 | 1 | 0 | 1 | 1 | 0.5 | 0.5 | 0 | 0 | 0 | 0 | 1 |
| 6 | 0.292022908 | 0.089573829 | 0.656464779 | 1 | 1 | 1 | 1 | 0 | 0.5 | 1 | 0 | 0 | 1 | 0 |
| 7 | 0.37524173 | 0.266297298 | 0.760057508 | 0 | 0 | 0 | 0 | 0 | 0.5 | 0 | 0 | 0 | 0 | 0 |
| 8 | 0.926133757 | 0.024862455 | 0.913019283 | 1 | 1 | 1 | 1 | 0 | 0.5 | 1 | 0 | 0 | 1 | 0 |
| 9 | 0.931565961 | 0.634721076 | 0.967057715 | 1 | 1 | 1 | 1 | 1 | 0.5 | 1 | 0 | 1 | 1 | 1 |
| 10 | 0.904847543 | 0.971932478 | 0.684887966 | 1 | 1 | 1 | 1 | 1 | 0.5 | 1 | 1 | 1 | 1 | 1 |
| 11 | 0.315853607 | -0.111989965 | 0.176068709 | 0 | 0 | 0 | 0 | 0 | 0 | 0 | 0 | 0 | 0 | 0 |
| 12 | 0.557465897 | 0.101633413 | 0.944153326 | 0.5 | 0 | 0 | 0 | 0 | 0.5 | 0 | 0 | 0 | 0 | 0 |
| 13 | 0.908341296 | -0.080202084 | 0.577253352 | 0.5 | 0 | 0 | 0 | 0 | 0.5 | 0 | 0 | 0 | 0 | 0 |
| 14 | 0.559267233 | 0.203601209 | 0.852512745 | 0.5 | 1 | 0 | 0 | 0 | 0.5 | 0.5 | 0 | 0 | 0.5 | 0 |
| 15 | 0.093815828 | 0.193875391 | 0.751570195 | 0.5 | 1 | 0 | 0.5 | 0 | 0.5 | 0 | 0 | 0 | 0 | 0 |
| 16 | 0.806382819 | 0.331439876 | 0.819277069 | 1 | 0 | 0.5 | 1 | 0 | 0.5 | 0 | 0 | 0 | 0 | 0 |
| 17 | 0.942353364 | 0.926143239 | 0.899983781 | 0 | 0 | 0 | 0 | 0 | 0.5 | 0 | 0 | 0 | 0 | 0 |
| 18 | 0.110507403 | 0.113528651 | 0.758269702 | 0 | 0 | 0 | 0 | 0 | 0.5 | 0 | 0 | 0 | 0 | 0 |
| 19 | 0.764195736 | 0.996200929 | 0.837979296 | 1 | 1 | 1 | 1 | 1 | 0.5 | 1 | 0 | 1 | 1 | 1 |
| 20 | 0.174686654 | 0.195070726 | 0.685939105 | 0 | 1 | 0 | 0 | 0 | 0.5 | 0.5 | 0 | 0 | 0.5 | 0 |
| 21 | 0.311085355 | 0.141793708 | 0.492081783 | 0 | 0 | 0 | 0 | 0 | 0.5 | 0 | 0 | 0 | 0 | 0 |
| 22 | 0.151897108 | 0.113748982 | 0.769622243 | 0 | 0 | 0 | 0 | 0 | 0.5 | 0 | 0 | 0 | 0 | 0 |
| 23 | 0.229200202 | 0.249717714 | 0.840349251 | 1 | 0 | 0.5 | 1 | 0 | 0.5 | 1 | 1 | 0 | 1 | 1 |
| 24 | 0.372827635 | 0.499400004 | 0.640270817 | 1 | 0 | 0 | 0 | 0 | 0.5 | 0 | 1 | 0 | 0 | 0 |
| 25 | 0.450820252 | 0.994386832 | 0.928739979 | 1 | 0.5 | 1 | 0.5 | 1 | 0.5 | 1 | 1 | 0.5 | 1 | 1 |
| 26 | 0.080546343 | 0.093779274 | 0.723865916 | 0 | 1 | 0 | 0 | 0 | 0.5 | 0 | 0 | 0 | 0 | 0 |
| 27 | 0.0457542 | 0.031414063 | 0.774068377 | 0 | 0.5 | 0 | 0 | 0 | 0.5 | 0 | 0 | 0 | 0 | 0 |
| 28 | 0.924775077 | 0.228884211 | 0.971262267 | 1 | 1 | 1 | 1 | 1 | 0.5 | 1 | 0 | 1 | 0.5 | 1 |
| 29 | 0.986550408 | 0.985638416 | 0.814151272 | 1 | 0.5 | 1 | 1 | 0 | 0 | 1 | 1 | 0 | 0.5 | 1 |
| 30 | 0.443776628 | 0.120165346 | 0.591942351 | 1 | 1 | 1 | 1 | 1 | 0.5 | 1 | 1 | 1 | 1 | 1 |
| 31 | 0.666245403 | 0.960790926 | 0.223943725 | 1 | 1 | 1 | 1 | 1 | 0.5 | 1 | 1 | 1 | 1 | 1 |
| 32 | 0.968341667 | 0.865542332 | 0.867506047 | 0 | 1 | 1 | 1 | 0 | 0.5 | 0.5 | 1 | 0 | 0 | 0 |
| 33 | 0.263120491 | 0.399762143 | 0.279131268 | 1 | 1 | 1 | 1 | 0 | 0.5 | 1 | 1 | 0 | 1 | 0.5 |
| 34 | 0.50191201 | 0.452945441 | 0.920975222 | 1 | 1 | 1 | 1 | 0.5 | 0.5 | 1 | 1 | 0.5 | 1 | 1 |
| 35 | 0.177210704 | -0.116995407 | 0.89757692 | 1 | 1 | 1 | 1 | 0 | 0.5 | 1 | 1 | 0.5 | 1 | 0 |
| 36 | 0.796737271 | 0.178104758 | 0.613100552 | 1 | 1 | 1 | 1 | 0 | 0.5 | 1 | 0 | 0 | 1 | 1 |
| 37 | 0.964597011 | 0.556078628 | 0.987016969 | 1 | 1 | 1 | 1 | 0 | 0.5 | 0 | 0 | 0 | 0 | 0 |
| 38 | 0.916986798 | -0.086974663 | 0.025428001 | 1 | 1 | 0.5 | 0.5 | 0 | 0.5 | 0 | 0 | 0 | 0 | 0 |
| 39 | 0.391814237 | 0.683598914 | 0.37508583 | 1 | 1 | 1 | 1 | 0 | 0.5 | 1 | 0 | 0 | 0.5 | 0.5 |
| 40 | 0.432593005 | 0.535664875 | 0.562915686 | 1 | 0 | 0 | 0 | 0 | 0.5 | 0 | 0 | 0 | 0 | 1 |
| 41 | 0.994990329 | 0.680086569 | 0.675633282 | 1 | 1 | 1 | 1 | 0 | 0.5 | 1 | 0 | 0 | 0.5 | 1 |
| 42 | 0.966251062 | 0.927353804 | 0.963920948 | 1 | 1 | 1 | 1 | 0 | 0.5 | 1 | 0 | 0 | 1 | 0.5 |
| 43 | 0.941954 | 0.215881383 | 0.507045798 | 1 | 0.5 | 1 | 1 | 0 | 0.5 | 0.5 | 0 | 0 | 0 | 0 |
| 44 | 0.862125373 | 0.079362299 | 0.078902863 | 1 | 1 | 1 | 1 | 0 | 0.5 | 1 | 0 | 0 | 1 | 0 |
| 45 | 0.515228066 | 0.000389522 | 0.947146537 | 0 | 0 | 0 | 0 | 0 | 0.5 | 0 | 0 | 0 | 0 | 0 |
| 46 | 0.494019082 | 0.11379871 | 0.537610947 | 0 | 0 | 0 | 0 | 0 | 0.5 | 0 | 0 | 0 | 0 | 0 |
| 47 | 0.506786384 | -0.039588922 | 0.60640882 | 0 | 0 | 0 | 0 | 0 | 0.5 | 0 | 0 | 0 | 0 | 0 |
| 48 | 0.569385375 | -0.232279593 | 0.719300375 | 0 | 0 | 0 | 0 | 0 | 0.5 | 0 | 0 | 0 | 0 | 0 |
| 49 | 0.36287104 | 0.13990727 | 0.732637606 | 0 | 0 | 0 | 0 | 0 | 0.5 | 0 | 0 | 0 | 0 | 0 |
| 50 | 0.40002337 | 0.03758239 | 0.9831453 | 0 | 0 | 0 | 0 | 0 | 0 | 0 | 0 | 0 | 0 | 0 |
| 51 | 0.005505534 | -0.022218187 | 0.935821188 | 1 | 0 | 0 | 0 | 0 | 0.5 | 0 | 0 | 0 | 0 | 0 |
| 52 | 0.43352058 | 0.213772613 | 0.838249455 | 0 | 0 | 0 | 0 | 0 | 0.5 | 0 | 0 | 0 | 0 | 0 |
| 53 | 0.123107078 | 0.094425763 | 0.598118963 | 0 | 0 | 0 | 0 | 0 | 0.5 | 0 | 0 | 0 | 0 | 0 |
| 54 | 0.812321128 | 0.67260405 | 0.713498684 | 0 | 0 | 0 | 0 | 0 | 0.5 | 0 | 0.5 | 0 | 0 | 0 |
| 55 | 0.374314769 | 0.194272775 | 0.693668639 | 0 | 0 | 0 | 0 | 0 | 0.5 | 0 | 0 | 0 | 0 | 0 |

**Supplemental Table 2.** MLST allele profiles for each strain, derived from the Pasteur MLST scheme.

| strain | acsA | aroE | guaA | mutL | nuoD | ppsA | trpE | MLST |
| --- | --- | --- | --- | --- | --- | --- | --- | --- |
| 1 |  |  |  |  |  |  |  | Unknown |
| 2 | 16 | 5 | 30 | 11 | 4 | 31 | 41 | 233(Pasteur) |
| 3 | 6 | 5 | 11 | 7 | 3 | 12 | 19 | 282(Pasteur) |
| 4 | 38 | 11 | 3 | 13 | 1 | 2 | 4 | 235(Pasteur) |
| 5 | 6 | 5 | 6 | 7 | 4 | 6 | 7 | 27(Pasteur) |
| 6 | 38 | 11 | 3 | 13 | 1 | 2 | 4 | 235(Pasteur) |
| 7 | 2 | 4 | 5 | 3 | 1 | 6 | 11 | 357(Pasteur) |
| 8 | 18 | 4 | 5 | 3 | 1 | 17 | 13 | 446(Pasteur) |
| 9 | 6 | 5 | 6 | 7 | 4 | 6 | 7 | 27(Pasteur) |
| 10 | 15 | 5 | 11 | 72 | 4 | 15 | 3 | 2012(Pasteur) |
| 11 | 13 | 8 | 9 | 3 | 1 | 6 | 9 | 316(Pasteur) |
| 12 | 38 | 11 | 3 | 13 | 1 | 2 | 4 | 235(Pasteur) |
| 13 | 17 | 5 | 5 | 4 | 4 | 4 | 3 | 111(Pasteur) |
| 14 | 16 | 5 | 30 | 11 | 4 | 31 | 41 | 233(Pasteur) |
| 15 | 16 | 5 | 30 | 11 | 4 | 31 | 41 | 233(Pasteur) |
| 16 | 47 | 8 | 7 | 6 | 8 | 11 | 40 | 313(Pasteur) |
| 17 | 18 | 4 | 5 | 3 | 1 | 17 | 13 | 446(Pasteur) |
| 18 |  |  |  |  |  |  |  | Unknown |
| 19 | 17 | 3 | 5 | 4 | 4 | 4 | 3 | 966(Pasteur) |
| 20 | 16 | 5 | 30 | 11 | 4 | 31 | 41 | 233(Pasteur) |
| 21 | 16 | 5 | 30 | 11 | 4 | 31 | 41 | 233(Pasteur) |
| 22 | 2 | 4 | 5 | 3 | 1 | 6 | 11 | 357(Pasteur) |
| 23 | 16 | 5 | 19 | 3 | 4 | 13 | 7 | 1600(Pasteur) |
| 24 | 38 | 11 | 3 | 13 | 1 | 2 | 4 | 235(Pasteur) |
| 25 | 17 | 3 | 5 | 4 | 4 | 4 | 3 | 966(Pasteur) |
| 26 |  |  |  |  |  |  |  | Unknown |
| 27 | 16 | 5 | 30 | 11 | 4 | 31 | 41 | 233(Pasteur) |
| 28 | 11 | 5 | 1 | 7 | 9 | 4 | 7 | 17(Pasteur) |
| 29 | 36 | 27 | 28 | 3 | 4 | 13 | 7 | 179(Pasteur) |
| 30 | 47 | 8 | 7 | 6 | 8 | 11 | 40 | 313(Pasteur) |
| 31 | 15 | 5 | 36 | 11 | 27 | 4 | 2 | 360(Pasteur) |
| 32 | 17 | 22 | 5 | 3 | 1 | 14 | 3 | 389(Pasteur) |
| 33 | 4 | 4 | 16 | 12 | 1 | 6 | 3 | 253(Pasteur) |
| 34 | 6 | 28 | 4 | 3 | 3 | 4 | 7 | 252(Pasteur) |
| 35 | 47 | 8 | 7 | 6 | 8 | 11 | 40 | 313(Pasteur) |
| 36 |  |  |  |  |  |  |  | Unknown |
| 37 | 28 | 22 | 5 | 3 | 3 | 14 | 19 | 175(Pasteur) |
| 38 | 17 | 3 | 5 | 4 | 4 | 4 | 3 | 966(Pasteur) |
| 39 | 11 | 5 | 1 | 7 | 9 | 4 | 7 | 17(Pasteur) |
| 40 | 6 | 28 | 4 | 3 | 3 | 4 | 7 | 252(Pasteur) |
| 41 | 17 | 3 | 5 | 4 | 4 | 4 | 3 | 966(Pasteur) |
| 42 | 16 | 5 | 19 | 3 | 4 | 13 | 7 | 1600(Pasteur) |
| 43 | 17 | 3 | 5 | 4 | 4 | 4 | 3 | 966(Pasteur) |
| 44 | 17 | 3 | 5 | 4 | 4 | 4 | 3 | 966(Pasteur) |
| 45 | 13 | 8 | 9 | 3 | 1 | 17 | 15 | 309(Pasteur) |
| 46 | 13 | 8 | 9 | 3 | 1 | 17 | 15 | 309(Pasteur) |
| 47 | 13 | 8 | 9 | 3 | 1 | 17 | 15 | 309(Pasteur) |
| 48 | 13 | 8 | 9 | 3 | 1 | 17 | 15 | 309(Pasteur) |
| 49 | 13 | 8 | 9 | 3 | 1 | 17 | 15 | 309(Pasteur) |
| 50 | 13 | 8 | 9 | 3 | 1 | 17 | 241 | 2775(Pasteur) |
| 51 | 13 | 8 | 9 | 3 | 1 | 17 | 15 | 309(Pasteur) |
| 52 | 13 | 8 | 9 | 3 | 1 | 17 | 15 | 309(Pasteur) |
| 53 | 13 | 8 | 9 | 3 | 1 | 17 | 15 | 309(Pasteur) |
| 54 | 6 | 5 | 6 | 7 | 4 | 6 | 7 | 27(Pasteur) |
| 55 | 13 | 8 | 9 | 3 | 1 | 17 | 15 | 309(Pasteur) |

**Supplemental Table 4.** Full statistics for phage Pagel’s Lambda results.

| Type | Trait | N | N_unique_values | Lambda | CI_lower | CI_upper | LogLik_full | LogLik_lambda_0 | LR_statistic | P_value | Q_value_BH | Significant_raw_p05 | Significant_FD_R_q05 |
| --- | --- | --- | --- | --- | --- | --- | --- | --- | --- | --- | --- | --- | --- |
| Phage | Luz19 | 55 | 55 | 0.506148821444281 | 0.185 | 0.755 | -7.5053472804163 | -13.1532460738195 | 11.2957975868064 | 0.000776827224694518 | 0.00217511622914465 | TRUE | TRUE |
| Phage | PAML31.1 | 55 | 55 | 0.458751267545214 | 0.115 | 0.745 | -14.3261015713367 | -18.327601794445 | 8.00300044621665 | 0.00466999024613501 | 0.00933998049227001 | TRUE | TRUE |
| Phage | OMKO1 | 55 | 55 | 0.000545076806112702 | 0 | 0.33 | -0.565063825576864 | -0.565063825576864 | 0 | 1 | 1 | FALSE | FALSE |

**Supplemental Table 5.** Full statistics for antibiotic Pagel’s Lambda results.

| Type | Trait | N | N_uniq<br>ue_valu<br>es | Lambda | CI_low<br>er | CI_up<br>per | LogLik<br>_full | LogLik<br>_lambd<br>a0 | LR_stat<br>istic | P_value | Q_value<br>_BH | Signific<br>ant_raw<br>_p05 | Signific<br>ant_FD<br>R_q05 |
| --- | --- | --- | --- | --- | --- | --- | --- | --- | --- | --- | --- | --- | --- |
| Antibiotic | Amikacin | 51 | 3 | 0.58362<br>3655815<br>233 | 0.29107<br>3117705<br>318 | 1 | -<br>36.1318<br>7711092<br>13 | -<br>48.9478<br>9199807<br>13 | 25.6320<br>297743 | 4.13124<br>8108706<br>68E-07 | 2.75057<br>6781702<br>83E-06 | TRUE | TRUE |
| Antibiotic | Aztreonam | 51 | 3 | 0.34136<br>1215222<br>115 | 0.08442<br>8956716<br>4309 | 1 | -<br>46.2180<br>8326022<br>07 | -<br>54.1415<br>9238788<br>54 | 15.8470<br>1825532<br>93 | 6.86746<br>6156396<br>19E-05 | 0.00024<br>0361315<br>473867 | TRUE | TRUE |
| Antibiotic | Cefepime | 51 | 3 | 0.54846<br>4101550<br>168 | 0.26418<br>3744512<br>241 | 1 | -<br>40.2512<br>7213413<br>41 | -<br>52.7245<br>6315718<br>29 | 24.9465<br>8204609<br>76 | 5.89409<br>3103648<br>92E-07 | 2.75057<br>6781702<br>83E-06 | TRUE | TRUE |
| Antibiotic | Ceftazidime | 51 | 3 | 0.53663<br>5784670<br>613 | 0.25390<br>4540043<br>245 | 1 | -<br>40.3447<br>3943031<br>49 | -<br>53.5353<br>7844869<br>61 | 26.3812<br>7803676<br>25 | 2.80243<br>7171755<br>1E-07 | 2.75057<br>6781702<br>83E-06 | TRUE | TRUE |
| Antibiotic | Ciprofloxacin | 51 | 3 | 2.09186<br>6208614<br>07E-15 | 6.38673<br>8024949<br>09E-16 | 6.85154<br>8032262<br>22E-15 | -<br>30.0042<br>8673005<br>06 | -<br>30.0043<br>7395158<br>62 | 0.00017<br>4443071<br>252028 | 0.98946<br>2094560<br>771 | 1 | FALSE | FALSE |
| Antibiotic | Colistin | 51 | 2 | 0.99999<br>9998591<br>375 | NA | NA | -<br>10.4030<br>9035743<br>61 | -<br>11.4096<br>1822337<br>46 | 2.01305<br>5731877<br>06 | 0.15595<br>0931016<br>245 | 0.24259<br>0337136<br>381 | FALSE | FALSE |
| Antibiotic | Doripenem | 51 | 3 | 0.60310<br>8440584<br>871 | 0.30567<br>3930700<br>863 | 1 | -<br>47.8637<br>5567197<br>34 | -<br>51.0477<br>0956154<br>17 | 6.36790<br>7779136<br>73 | 0.01162<br>0252077<br>6147 | 0.02033<br>5441135<br>8256 | TRUE | TRUE |
| Antibiotic | Imipenem | 51 | 3 | 0.54031<br>0732936<br>18 | 0.35891<br>9837412<br>98 | 0.81337<br>2953220<br>54 | -<br>35.7117<br>5300223<br>9 | -<br>36.1429<br>8921262<br>57 | 0.86247<br>2420773<br>344 | 0.35304<br>7725948<br>238 | 0.46001<br>3765440<br>368 | FALSE | FALSE |
| Antibiotic | Levofloxacin | 51 | 3 | 2.08091<br>2840005<br>05E-93 | 9.00651<br>6133782<br>2E-94 | 4.80785<br>0431151<br>64E-93 | -<br>30.0042<br>8673004<br>97 | -<br>30.0043<br>6716221<br>86 | 0.00016<br>0864337<br>857447 | 0.98988<br>0517453<br>254 | 1 | FALSE | FALSE |
| Antibiotic | Meropenem | 51 | 3 | 0.42742<br>9926201<br>871 | NA | NA | -<br>48.6917<br>2465843<br>25 | -<br>49.1081<br>6757214<br>85 | 0.83288<br>5827432<br>136 | 0.36143<br>9387131<br>718 | 0.46001<br>3765440<br>368 | FALSE | FALSE |
| Antibiotic | Tobramycin | 51 | 3 | 0.67717<br>1643675<br>166 | 0.15926<br>7620696<br>128 | 1 | -<br>39.5261<br>8454628<br>35 | -<br>42.8142<br>2591668<br>35 | 6.57608<br>2740800<br>06 | 0.01033<br>5810945<br>1815 | 0.02033<br>5441135<br>8256 | TRUE | TRUE |

**Supplemental Table 6.** Kendall’s Tau correlation values by pairing, with P values, phylogenetically controlled (partial Kendall’s Tau)

| Antimicrobial 1 | Antimicrobial 2 | Kendall Tau | P Value |
| --- | --- | --- | --- |
| Luz19 | PAML31.1 | 0.309165934351616 | 0.000964767846046001 |
| Luz19 | OMKO1 | 0.114274037489596 | 0.222475959443205 |
| Luz19 | Amikacin | 0.217317895472959 | 0.0203381738672905 |
| Luz19 | Aztreonam | 0.199465009672516 | 0.0332165914958712 |
| Luz19 | Cefepime | 0.40472762873157 | 1.55460763325196E-05 |
| Luz19 | Ceftazidime | 0.376137413806579 | 5.92995429048305E-05 |
| Luz19 | Ciprofloxacin | 0.202083314735213 | 0.0309738799553664 |
| Luz19 | Colistin | -0.0479002093303253 | 0.609088687786896 |
| Luz19 | Doripenem | 0.242820765365502 | 0.00953326030752416 |
| Luz19 | Imipenem | 0.0491086764496132 | 0.600086501503812 |
| Luz19 | Levofloxacin | 0.143041403789618 | 0.126739524762672 |
| Luz19 | Meropenem | 0.159691964841854 | 0.088223325997386 |
| Luz19 | Tobramycin | 0.21906842319661 | 0.0193489804835644 |
| PAML31.1 | OMKO1 | 0.143816192295581 | 0.12469593203811 |
| PAML31.1 | Amikacin | 0.211705706062045 | 0.0238128629199807 |
| PAML31.1 | Aztreonam | 0.128355444117926 | 0.170592794721401 |
| PAML31.1 | Cefepime | 0.318005819509644 | 0.000686342071815776 |
| PAML31.1 | Ceftazidime | 0.290683590822869 | 0.00191381984191577 |
| PAML31.1 | Ciprofloxacin | 0.321063695436567 | 0.000608890369593415 |
| PAML31.1 | Colistin | 0.0504771189570708 | 0.589965939224602 |
| PAML31.1 | Doripenem | 0.282525040940641 | 0.00255972218837948 |
| PAML31.1 | Imipenem | 0.275102018879302 | 0.00331464229965603 |
| PAML31.1 | Levofloxacin | 0.199678298273253 | 0.0330288393170906 |
| PAML31.1 | Meropenem | 0.219779320613198 | 0.0189594275274181 |
| PAML31.1 | Tobramycin | 0.44859811632647 | 1.67481375608889E-06 |
| OMKO1 | Amikacin | -0.0237524575605046 | 0.799822123178491 |
| OMKO1 | Aztreonam | 0.0225092679631529 | 0.810093641581864 |
| OMKO1 | Cefepime | 0.0794038496407648 | 0.396603734387335 |
| OMKO1 | Ceftazidime | 0.0753732234424847 | 0.421008928808586 |
| OMKO1 | Ciprofloxacin | 0.175900767440951 | 0.0603965490487416 |
| OMKO1 | Colistin | -0.03299275200676 | 0.724669398929661 |
| OMKO1 | Doripenem | 0.0145971321084693 | 0.876161841892514 |
| OMKO1 | Imipenem | -0.00701478250049314 | 0.940303299226186 |
| OMKO1 | Levofloxacin | 0.154236504322655 | 0.0996395789327826 |
| OMKO1 | Meropenem | 0.0265384807583467 | 0.776931210292703 |
| OMKO1 | Tobramycin | 0.0301707676602921 | 0.747378351055097 |
| Amikacin | Aztreonam | 0.335392989012174 | 0.000342805868377063 |
| Amikacin | Cefepime | 0.664907772981135 | 1.26167808563825E-12 |
| Amikacin | Ceftazidime | 0.676081501414128 | 5.28607896372736E-13 |
| Amikacin | Ciprofloxacin | 0.301405748558831 | 0.00129196766275523 |
| Amikacin | Colistin | 0.0867186029869962 | 0.354553310812364 |
| Amikacin | Doripenem | 0.562237662385647 | 1.94474150764206E-09 |
| Amikacin | Imipenem | 0.197004688755137 | 0.0354492045889953 |
| Amikacin | Levofloxacin | 0.281861749948314 | 0.00262014651328336 |
| Amikacin | Meropenem | 0.536784610639398 | 1.00061911309326E-08 |
| Amikacin | Tobramycin | 0.56588980654397 | 1.52833686890107E-09 |
| Aztreonam | Cefepime | 0.616833702394945 | 4.54228271749576E-11 |
| Aztreonam | Ceftazidime | 0.604442872707195 | 1.09706354511814E-10 |
| Aztreonam | Ciprofloxacin | 0.237558041993191 | 0.0112084975599151 |
| Aztreonam | Colistin | 0.130132943390392 | 0.164748285221523 |
| Aztreonam | Doripenem | 0.656674234953913 | 2.37390331282182E-12 |
| Aztreonam | Imipenem | 0.109165894206356 | 0.243842120980373 |
| Aztreonam | Levofloxacin | 0.330561335889942 | 0.000417104358314489 |
| Aztreonam | Meropenem | 0.607227137741822 | 9.01214129389772E-11 |
| Aztreonam | Tobramycin | 0.267051415680937 | 0.00435824039611277 |
| Cefepime | Ceftazidime | 0.926430130435923 | 4.58124125372512E-23 |
| Cefepime | Ciprofloxacin | 0.433835208189132 | 3.62932945232018E-06 |
| Cefepime | Colistin | 0.0238707849774563 | 0.798846246949169 |
| Cefepime | Doripenem | 0.784858981695852 | 5.33679719549428E-17 |
| Cefepime | Imipenem | 0.316893492699464 | 0.000716718467994659 |
| Cefepime | Levofloxacin | 0.420224424880861 | 7.2488840724223E-06 |
| Cefepime | Meropenem | 0.680721312660851 | 3.6682960290962E-13 |
| Cefepime | Tobramycin | 0.553521204739291 | 3.43562414690904E-09 |
| Ceftazidime | Ciprofloxacin | 0.357128831042333 | 0.000137485602841555 |
| Ceftazidime | Colistin | 0.0299282677379648 | 0.749340381293581 |
| Ceftazidime | Doripenem | 0.747344514788569 | 1.48090291333006E-15 |
| Ceftazidime | Imipenem | 0.28800103180418 | 0.00210748672986299 |
| Ceftazidime | Levofloxacin | 0.356131255256431 | 0.000143533600883017 |
| Ceftazidime | Meropenem | 0.646180656781688 | 5.25478243668303E-12 |
| Ceftazidime | Tobramycin | 0.52575558386889 | 1.98987114411017E-08 |
| Ciprofloxacin | Colistin | 0.0810413043354187 | 0.38693803235154 |
| Ciprofloxacin | Doripenem | 0.392989944602977 | 2.72275361859152E-05 |
| Ciprofloxacin | Imipenem | 0.310534596722036 | 0.000915723064104513 |
| Ciprofloxacin | Levofloxacin | 0.868653411984192 | 1.79966125365605E-20 |
| Ciprofloxacin | Meropenem | 0.429176244670938 | 4.60958173332059E-06 |
| Ciprofloxacin | Tobramycin | 0.650330810278615 | 3.84337116111E-12 |
| Colistin | Doripenem | 0.00683234445522147 | 0.941853082220288 |
| Colistin | Imipenem | -0.102665401266671 | 0.273061761686651 |
| Colistin | Levofloxacin | 0.0769095305730228 | 0.411604430559186 |
| Colistin | Meropenem | 0.0466635648256965 | 0.618362452982349 |
| Colistin | Tobramycin | -0.0328449247749715 | 0.725853195403791 |
| Doripenem | Imipenem | 0.383939903438346 | 4.15155936583245E-05 |
| Doripenem | Levofloxacin | 0.477651100446476 | 3.40868108408526E-07 |
| Doripenem | Meropenem | 0.900579918988044 | 6.94846974030866E-22 |
| Doripenem | Tobramycin | 0.577643785254357 | 6.9672725017119E-10 |
| Imipenem | Levofloxacin | 0.404957468913703 | 1.53741004797603E-05 |
| Imipenem | Meropenem | 0.374531484500686 | 6.37623351680851E-05 |
| Imipenem | Tobramycin | 0.3696154669445 | 7.94829693938726E-05 |
| Levofloxacin | Meropenem | 0.51622891501109 | 3.56446751802703E-08 |
| Levofloxacin | Tobramycin | 0.546467662843488 | 5.41145883541978E-09 |
| Meropenem | Tobramycin | 0.569571133922189 | 1.1969825404119E-09 |

**Supplemental Table 7.** Kendall’s Tau correlation values by pairing, with P values, not phylogenetically controlled.

| Antimicrobial 1 | Antimicrobial 2 | Kendall_Tau | P_Value |
| --- | --- | --- | --- |
| Luz19 | PAML31.1 | 0.314478114478114 | 0.000698403200632007 |
| Luz19 | OMKO1 | 0.113804713804714 | 0.219874359864308 |
| Luz19 | Amikacin | 0.222846300488227 | 0.0410005705675774 |
| Luz19 | Aztreonam | 0.207789546502013 | 0.056667800520718 |
| Luz19 | Cefepime | 0.400719596884288 | 0.000257843311934169 |
| Luz19 | Ceftazidime | 0.371551330492151 | 0.00070288527649898 |
| Luz19 | Ciprofloxacin | 0.210728934081251 | 0.0567382213751934 |
| Luz19 | Colistin | -0.0415531995559314 | 0.710917113388619 |
| Luz19 | Doripenem | 0.24950672135471 | 0.0221522725024653 |
| Luz19 | Imipenem | 0.0644326986282696 | 0.562349755518756 |
| Luz19 | Levofloxacin | 0.154816264713781 | 0.159990120498132 |
| Luz19 | Meropenem | 0.170136723721774 | 0.114713439992402 |
| Luz19 | Tobramycin | 0.225987299157516 | 0.0385118098256637 |
| PAML31.1 | OMKO1 | 0.139393939393939 | 0.132910025313192 |
| PAML31.1 | Amikacin | 0.278557875610283 | 0.0106369361110816 |
| PAML31.1 | Aztreonam | 0.19535769329249 | 0.073158867321226 |
| PAML31.1 | Cefepime | 0.366346358325365 | 0.000835144661936959 |
| PAML31.1 | Ceftazidime | 0.344496621961169 | 0.00167958949778114 |
| PAML31.1 | Ciprofloxacin | 0.361249601282144 | 0.00108975690758485 |
| PAML31.1 | Colistin | 0.0706404392450834 | 0.528654073421974 |
| PAML31.1 | Doripenem | 0.329059589033023 | 0.00255143924198161 |
| PAML31.1 | Imipenem | 0.319902696698251 | 0.00402180925711214 |
| PAML31.1 | Levofloxacin | 0.254697725819447 | 0.0207984125146413 |
| PAML31.1 | Meropenem | 0.267357708705644 | 0.0131860827519086 |
| PAML31.1 | Tobramycin | 0.469358236711764 | 1.72413981053547E-05 |
| OMKO1 | Amikacin | -0.0233629185995721 | 0.830359558245747 |
| OMKO1 | Aztreonam | 0.019535769329249 | 0.857793357869626 |
| OMKO1 | Cefepime | 0.073269271665073 | 0.504021744094166 |
| OMKO1 | Ceftazidime | 0.0685385949451541 | 0.531935544978916 |
| OMKO1 | Ciprofloxacin | 0.168081411707664 | 0.128581072118439 |
| OMKO1 | Colistin | -0.0332425596447451 | 0.766848475677745 |
| OMKO1 | Doripenem | 0.0126561380397317 | 0.907616778096202 |
| OMKO1 | Imipenem | -0.00791278755084013 | 0.943279295948559 |
| OMKO1 | Levofloxacin | 0.144828118603215 | 0.188693849701051 |
| OMKO1 | Meropenem | 0.0243052462459677 | 0.821717357687815 |
| OMKO1 | Tobramycin | 0.02897273066122 | 0.790775855103335 |
| Amikacin | Aztreonam | 0.432494813542581 | 0.000769741703292617 |
| Amikacin | Cefepime | 0.709731993877846 | 4.08174025794427E-08 |
| Amikacin | Ceftazidime | 0.724433120259706 | 2.1296855450943E-08 |
| Amikacin | Ciprofloxacin | 0.37659765228037 | 0.00389694542228402 |
| Amikacin | Colistin | 0.119212866575011 | 0.367516576388719 |
| Amikacin | Doripenem | 0.612797635822239 | 1.89744266105448E-06 |
| Amikacin | Imipenem | 0.289609561315821 | 0.0272990902331651 |
| Amikacin | Levofloxacin | 0.374850089940042 | 0.00392541888780205 |
| Amikacin | Meropenem | 0.584985934561171 | 4.2388676577588E-06 |
| Amikacin | Tobramycin | 0.579910389719674 | 6.72101566167934E-06 |
| Aztreonam | Cefepime | 0.66320217011254 | 2.91808241406712E-07 |
| Aztreonam | Ceftazidime | 0.65643990505311 | 3.84382209952561E-07 |
| Aztreonam | Ciprofloxacin | 0.310961791316029 | 0.0171324649729665 |
| Aztreonam | Colistin | 0.156164969657582 | 0.237714895159303 |
| Aztreonam | Doripenem | 0.692601760603008 | 7.21816387863572E-08 |
| Aztreonam | Imipenem | 0.199742534108086 | 0.127848259455171 |
| Aztreonam | Levofloxacin | 0.406655026217331 | 0.00175110303146331 |
| Aztreonam | Meropenem | 0.644570436236096 | 4.00130658443757E-07 |
| Aztreonam | Tobramycin | 0.308189495463132 | 0.0166992426112692 |
| Cefepime | Ceftazidime | 0.935195082093422 | 6.4770194351193E-13 |
| Cefepime | Ciprofloxacin | 0.485255744879414 | 0.000217076542940123 |
| Cefepime | Colistin | 0.0558171655583031 | 0.674815025301252 |
| Cefepime | Doripenem | 0.806314659856016 | 4.56726464889063E-10 |
| Cefepime | Imipenem | 0.382644359255739 | 0.0037333242325047 |
| Cefepime | Levofloxacin | 0.483003965931659 | 0.000219595166655178 |
| Cefepime | Meropenem | 0.709802740423545 | 2.86750882383915E-08 |
| Cefepime | Tobramycin | 0.572097530461051 | 1.00206017741719E-05 |
| Ceftazidime | Ciprofloxacin | 0.419955192835313 | 0.00137095157341879 |
| Ceftazidime | Colistin | 0.063995637315692 | 0.630489510764121 |
| Ceftazidime | Doripenem | 0.773610015920683 | 2.22231344831493E-09 |
| Ceftazidime | Imipenem | 0.361807383143103 | 0.00610599078593126 |
| Ceftazidime | Levofloxacin | 0.431382640224006 | 0.000965114929638706 |
| Ceftazidime | Meropenem | 0.679928778958982 | 1.06100161799241E-07 |
| Ceftazidime | Tobramycin | 0.545793907871637 | 2.5111464881584E-05 |
| Ciprofloxacin | Colistin | 0.104491167214653 | 0.436230715856846 |
| Ciprofloxacin | Doripenem | 0.442863877435301 | 0.000689080051526708 |
| Ciprofloxacin | Imipenem | 0.364266230009518 | 0.00620814373720085 |
| Ciprofloxacin | Levofloxacin | 0.879079276192867 | 2.60762685604644E-11 |
| Ciprofloxacin | Meropenem | 0.472853613174189 | 0.000247550464433982 |
| Ciprofloxacin | Tobramycin | 0.663797913347633 | 3.77052560712402E-07 |
| Colistin | Doripenem | 0.0362591356639103 | 0.784043898768489 |
| Colistin | Imipenem | -0.0697529193733527 | 0.605287825353237 |
| Colistin | Levofloxacin | 0.10400628679223 | 0.436591057655158 |
| Colistin | Meropenem | 0.0722121934473044 | 0.580958593199882 |
| Colistin | Tobramycin | -0.0148983820282184 | 0.910463902323082 |
| Doripenem | Imipenem | 0.437042401893306 | 0.000866992915340557 |
| Doripenem | Levofloxacin | 0.527964928777866 | 4.87177191559515E-05 |
| Doripenem | Meropenem | 0.908769458092314 | 9.05409154723235E-13 |
| Doripenem | Tobramycin | 0.59508702457554 | 3.84406246535731E-06 |
| Imipenem | Levofloxacin | 0.458983352530419 | 0.000537179968378109 |
| Imipenem | Meropenem | 0.424620053205624 | 0.00106602897886769 |
| Imipenem | Tobramycin | 0.398805556562867 | 0.00240527953433005 |
| Levofloxacin | Meropenem | 0.559116645339574 | 1.36088233550079E-05 |
| Levofloxacin | Tobramycin | 0.565817806081494 | 1.37971370770347E-05 |
| Meropenem | Tobramycin | 0.587411285410158 | 3.99436198855566E-06 |

**Supplemental Table 8.** Overall Antibiotic and Phage resistance correlation values via four different tests, varying in their approach to phylogenetic correction. Confidence intervals, exact P values, and method of calculating confidence intervals included.

| Test | N | Estimate | CI_lower_95 | CI_upper_95 | P_value | CI_method |
| --- | --- | --- | --- | --- | --- | --- |
| **Simple Spearman**<br>**(no phylogenetic**<br>**correction)** | 55 | 0.409426273013103 | 0.165812177696411 | 0.615950658431408 | 0.00190962693061229 | Nonparametric bootstrap, 2000 resamples (percentile method) |
| **Partial Spearman**<br>**(controlling for**<br>**phylogeny via**<br>**PC1 only, ~49%**<br>**of variance)** | 51 | 0.358279127546917 | 0.0533095839703745 | 0.595592115966338 | 0.0106257727100883 | Nonparametric bootstrap, 2000 resamples (percentile method); 2000/2000 resamples valid |
| **Partial Mantel**<br>**(controlling for**<br>**FULL**<br>**phylogenetic**<br>**distance matrix)** | 51 | 0.181896068983736 | 0.0596546457904211 | 0.418087983017112 | 0.001 | P-value: Mantel permutation test (999 perms). CI: bootstrap over strains, 2000 resamples |
| **PGLS slope with**<br>**Pagel's lambda**<br>**(full-tree**<br>**covariance)** | 51 | 0.218129213253479 | 0.00645451706353828 | 0.429803909443419 | 0.043661413140997 | Wald-type profile CI from PGLS fit (nlme::intervals); P-value from GLS t-test |

**Supplemental Table 9.**
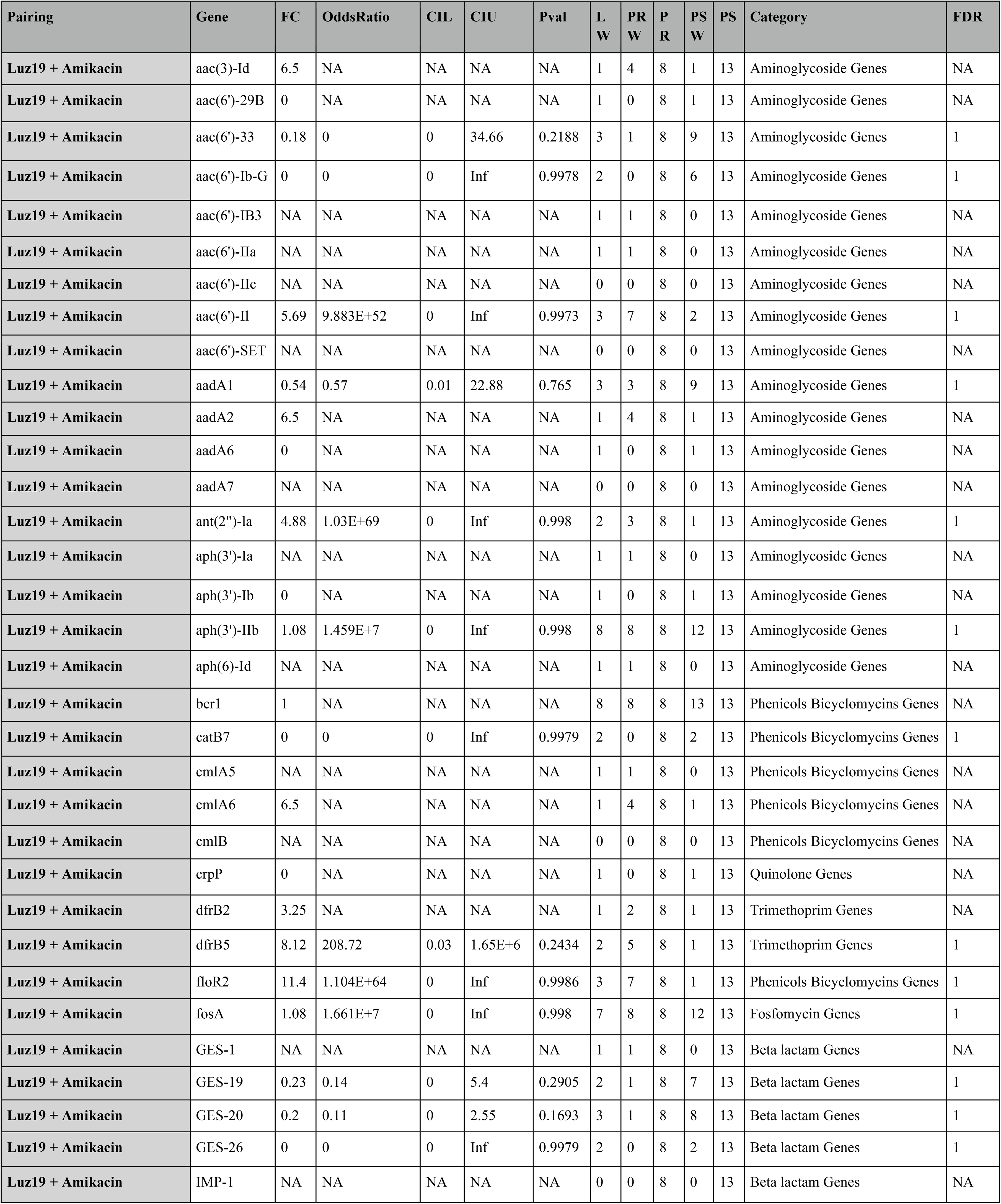
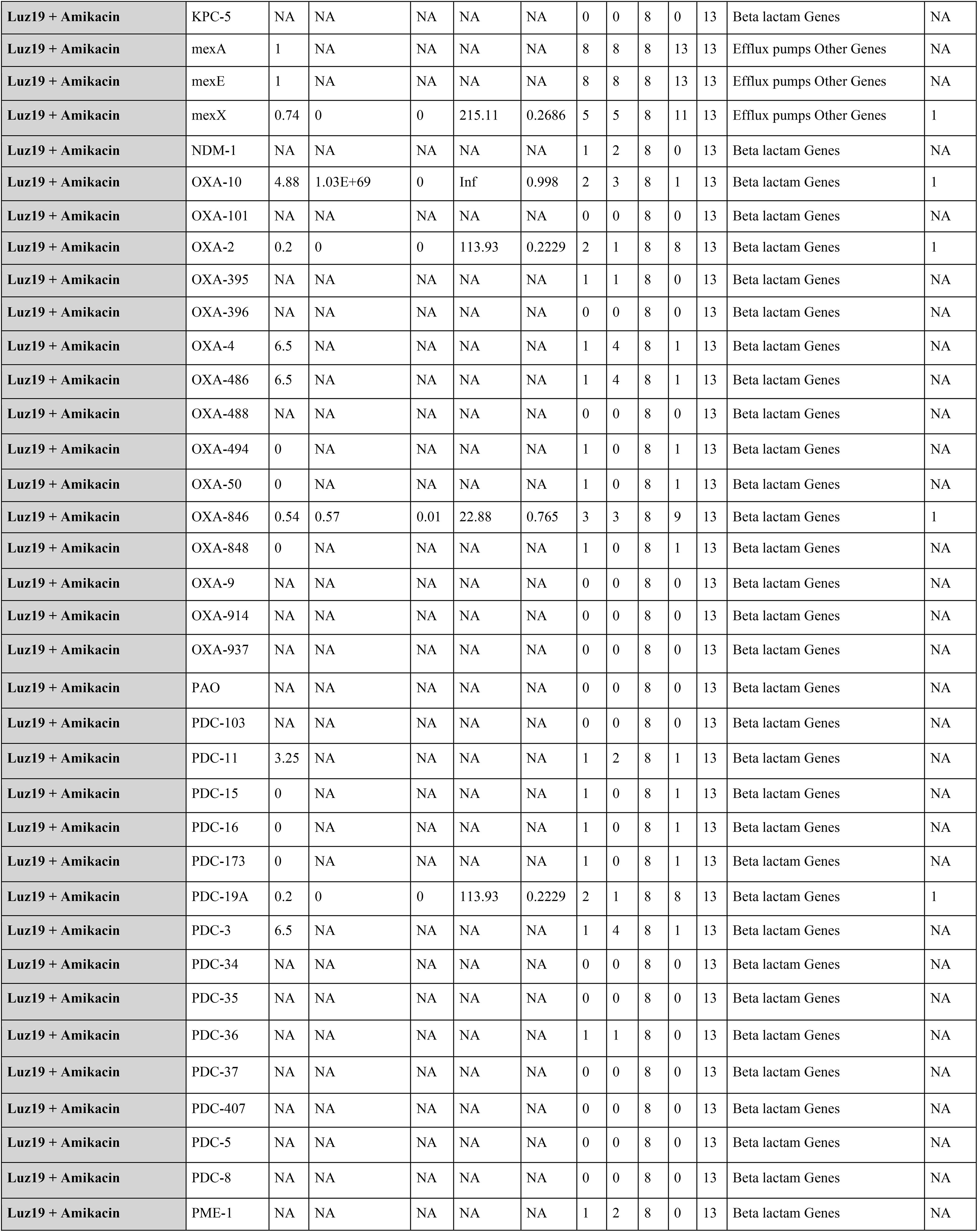
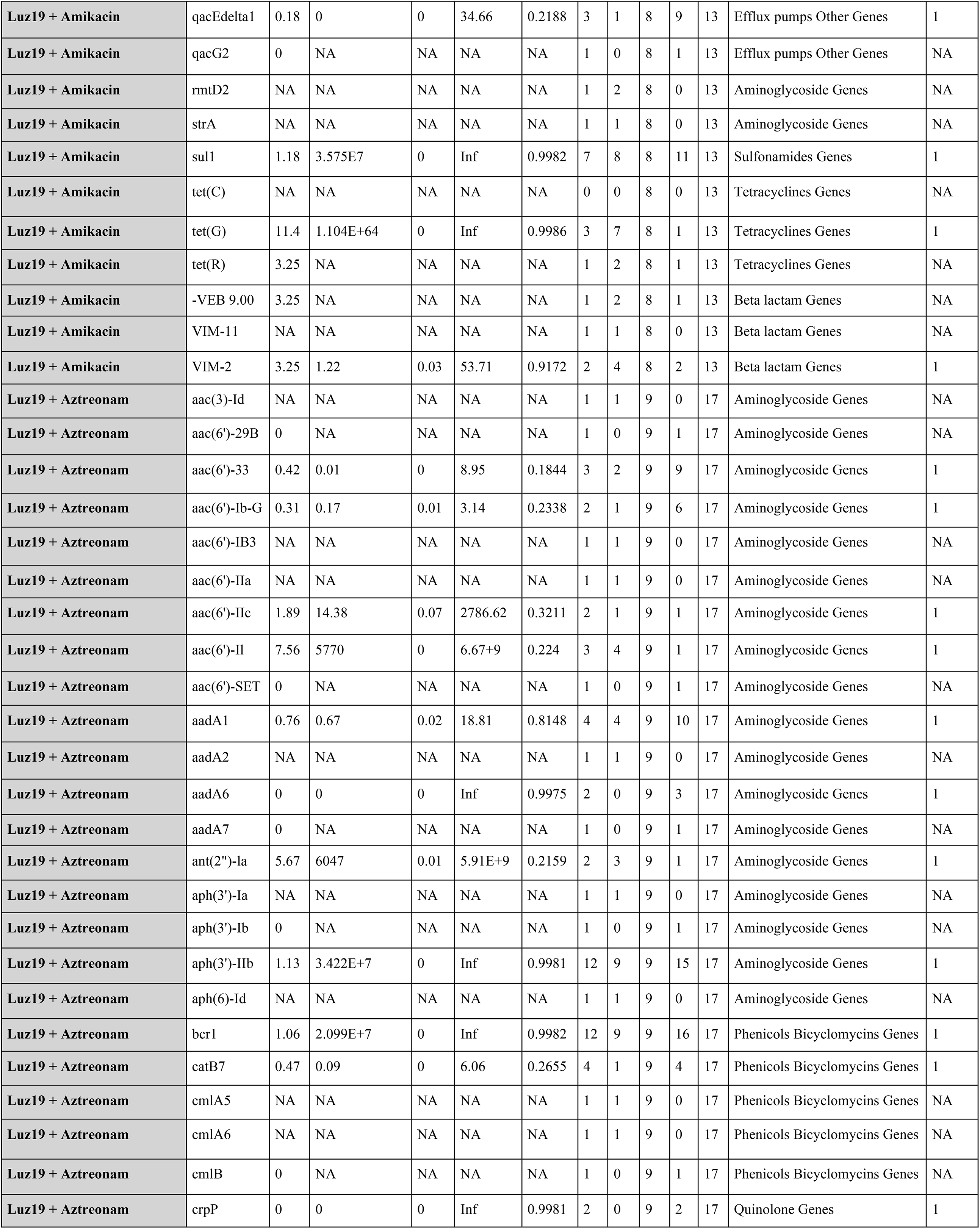
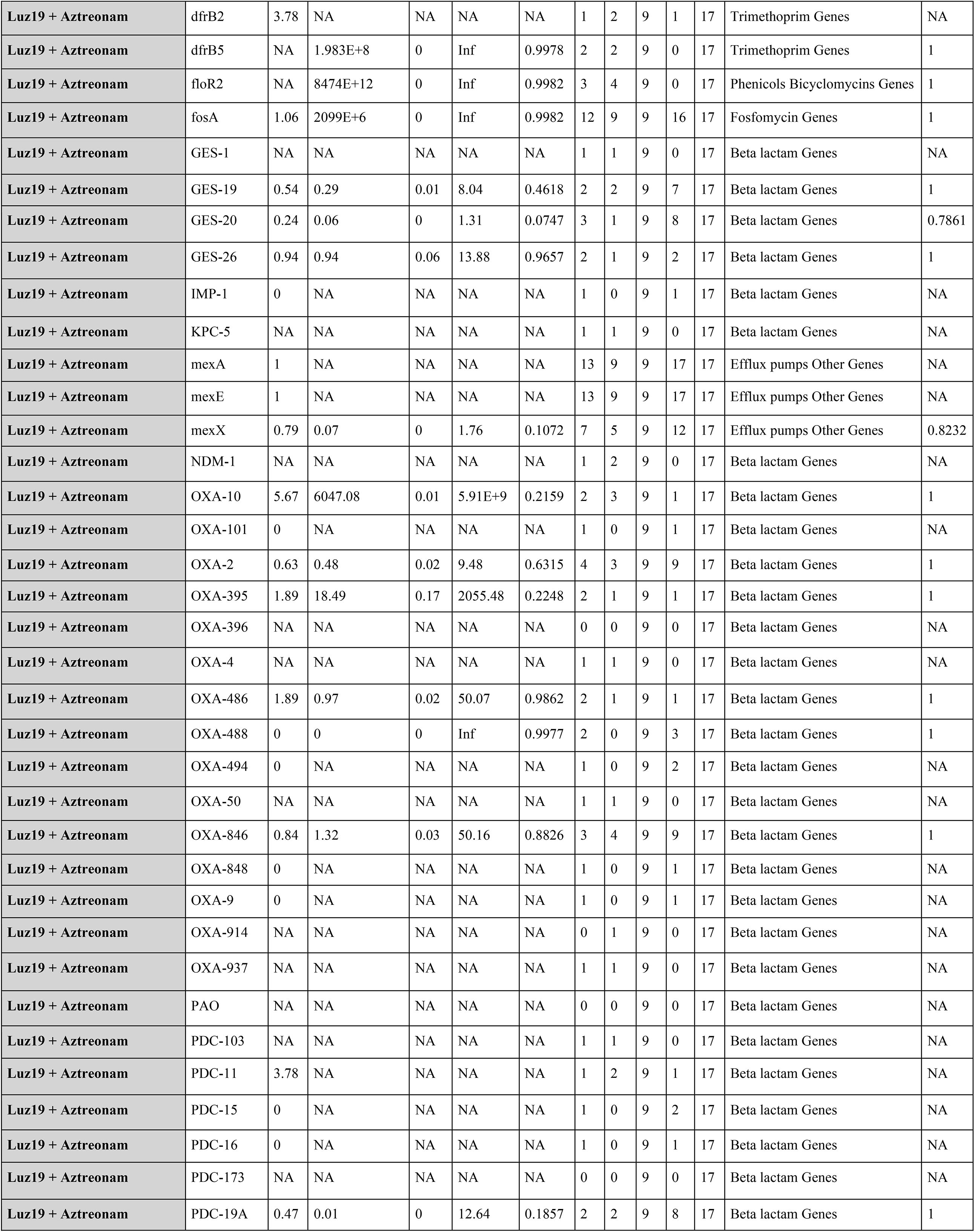
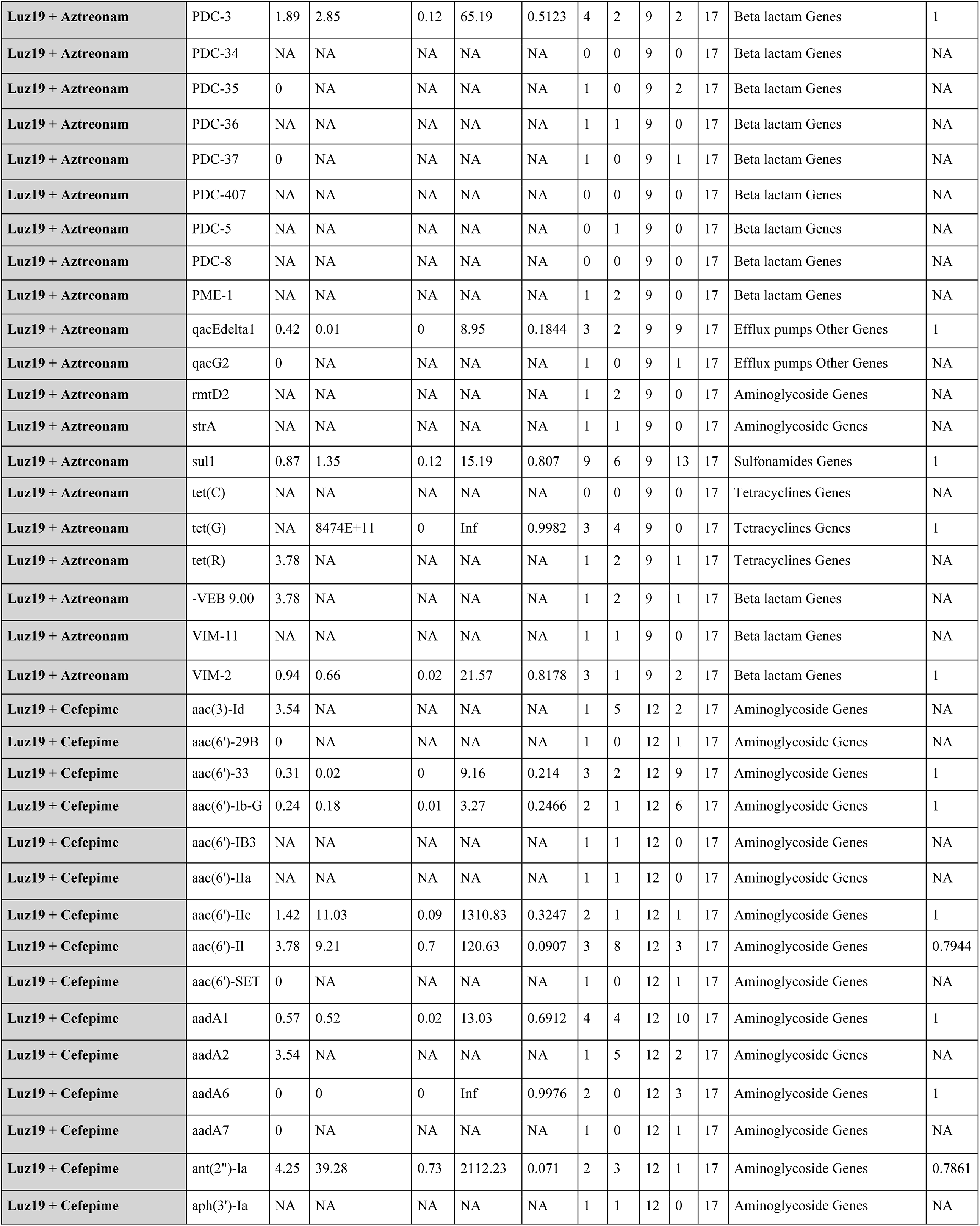
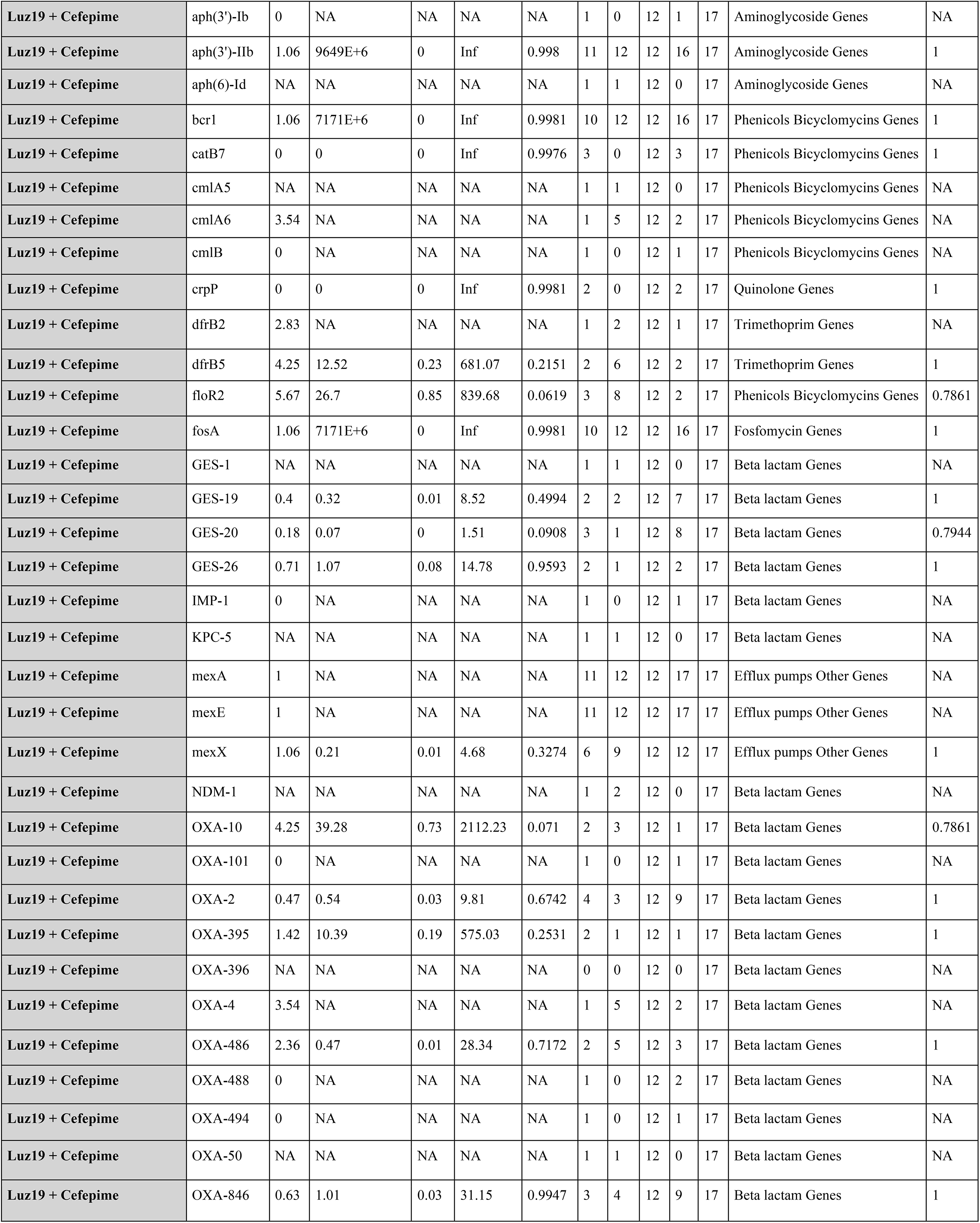
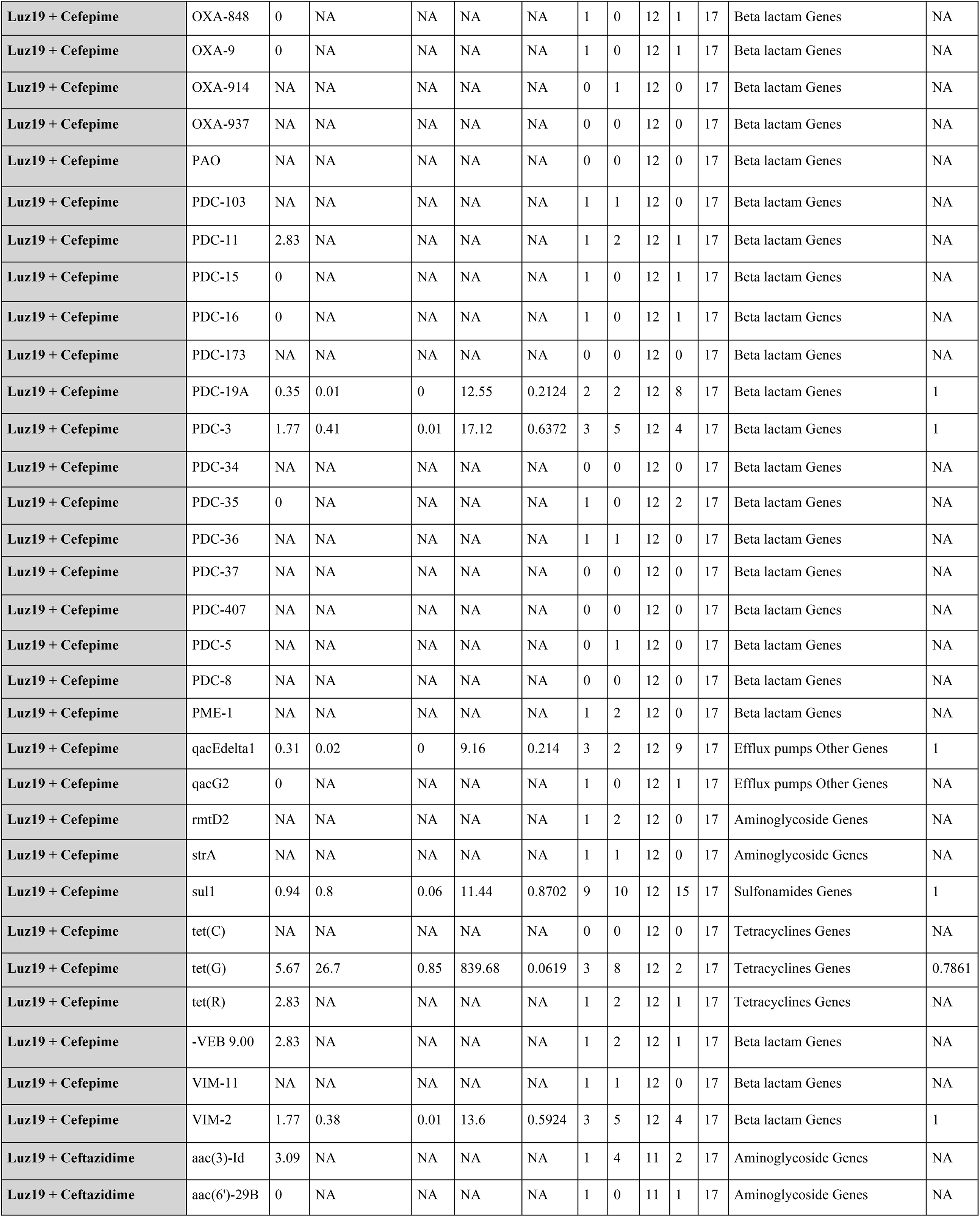
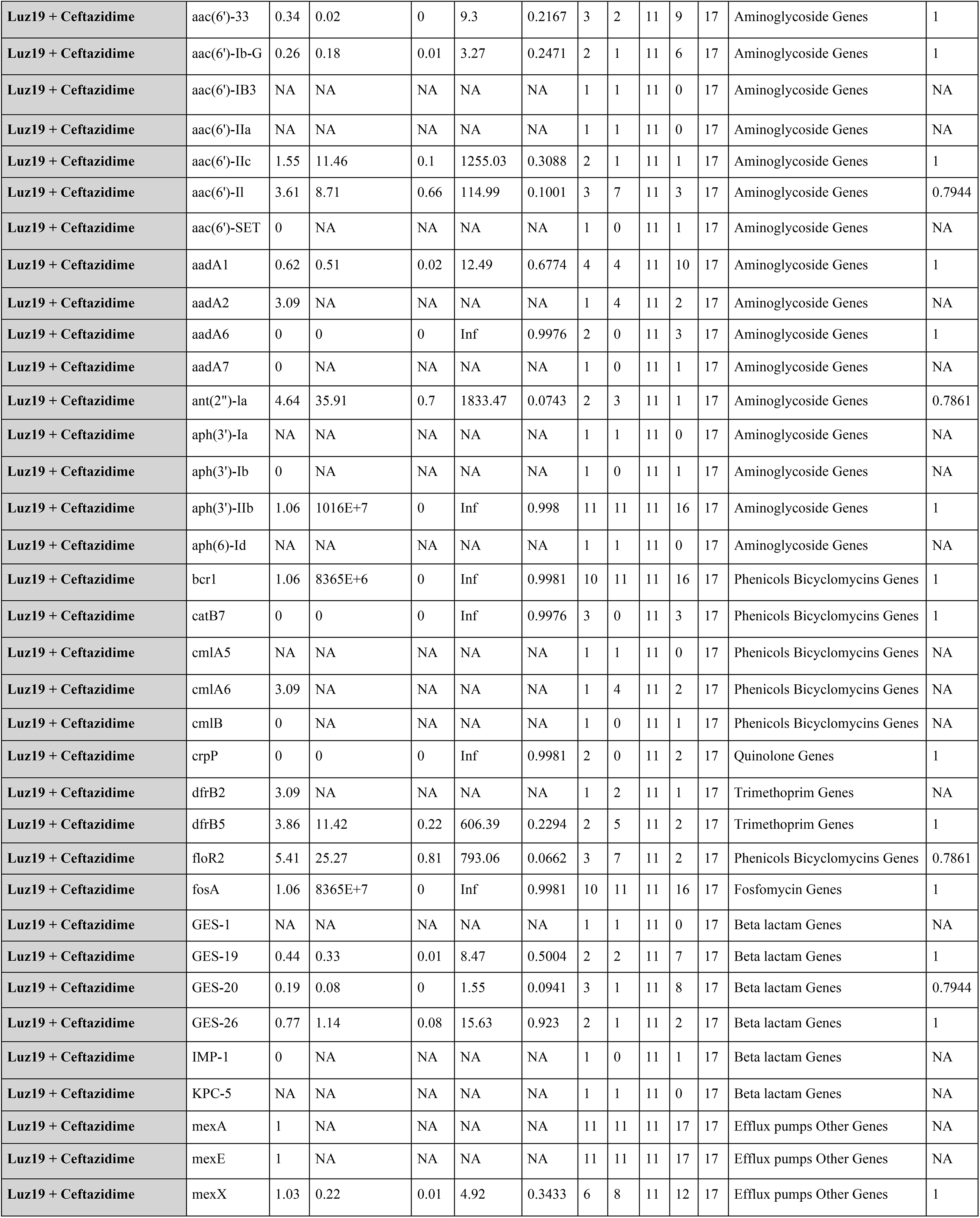
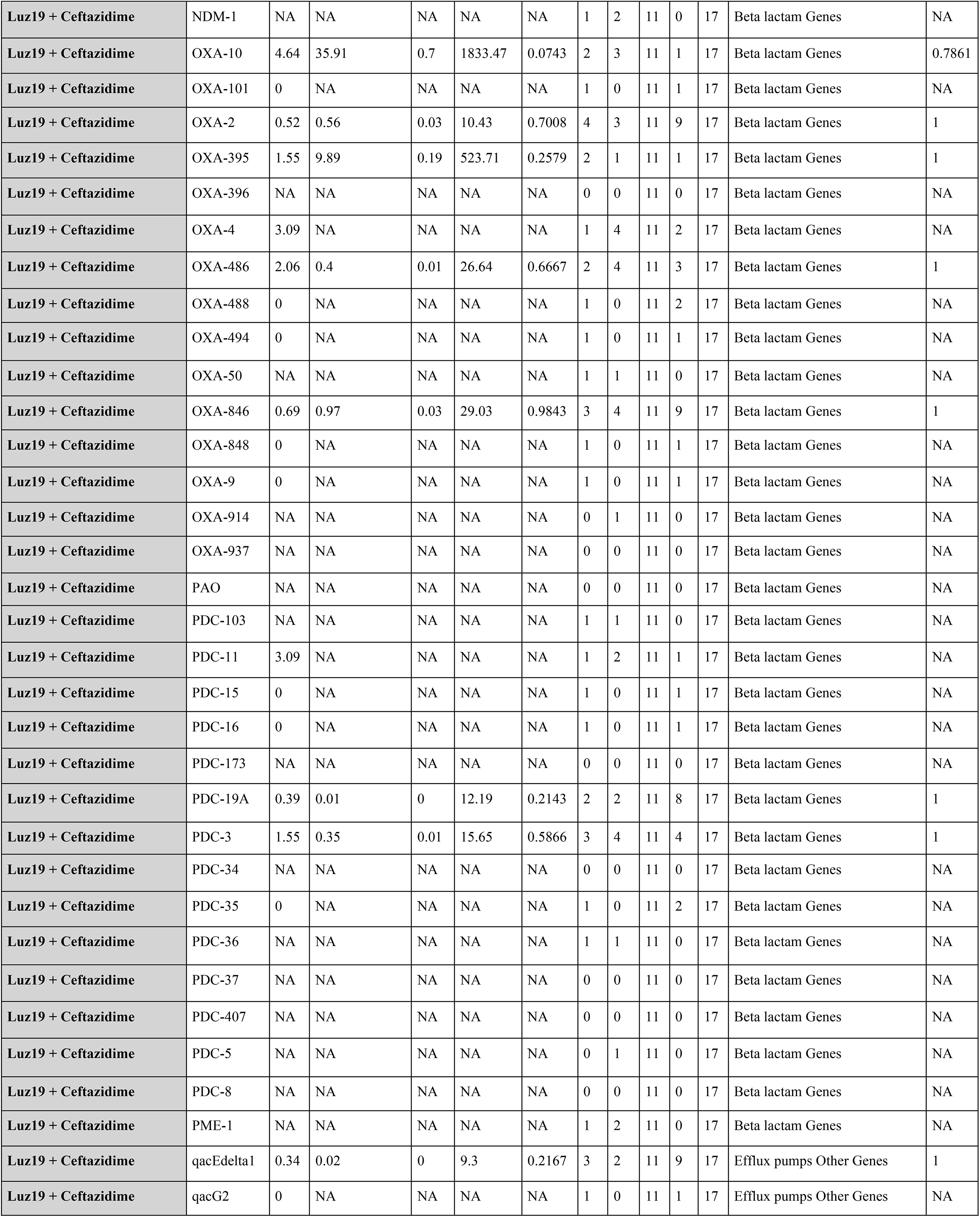
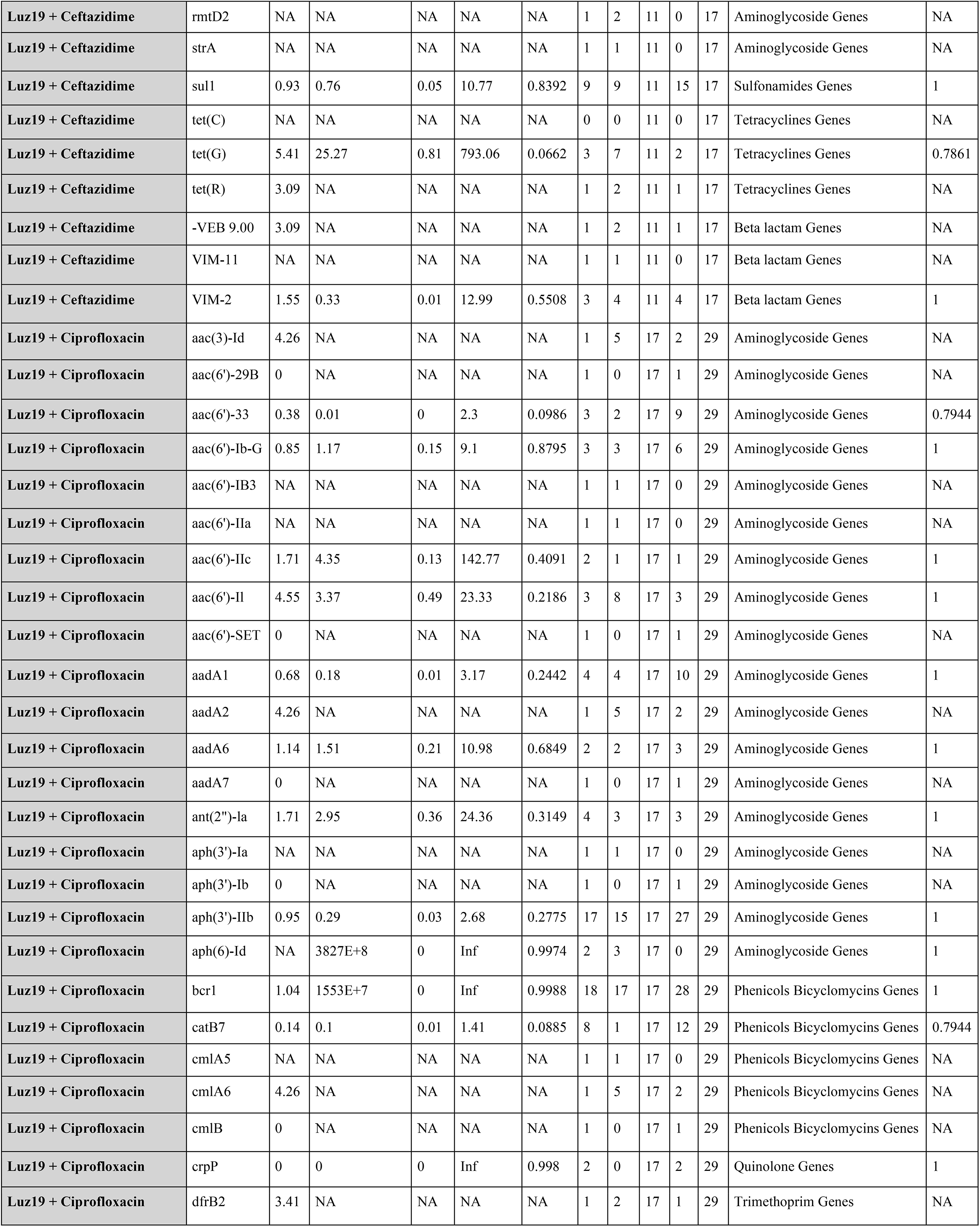
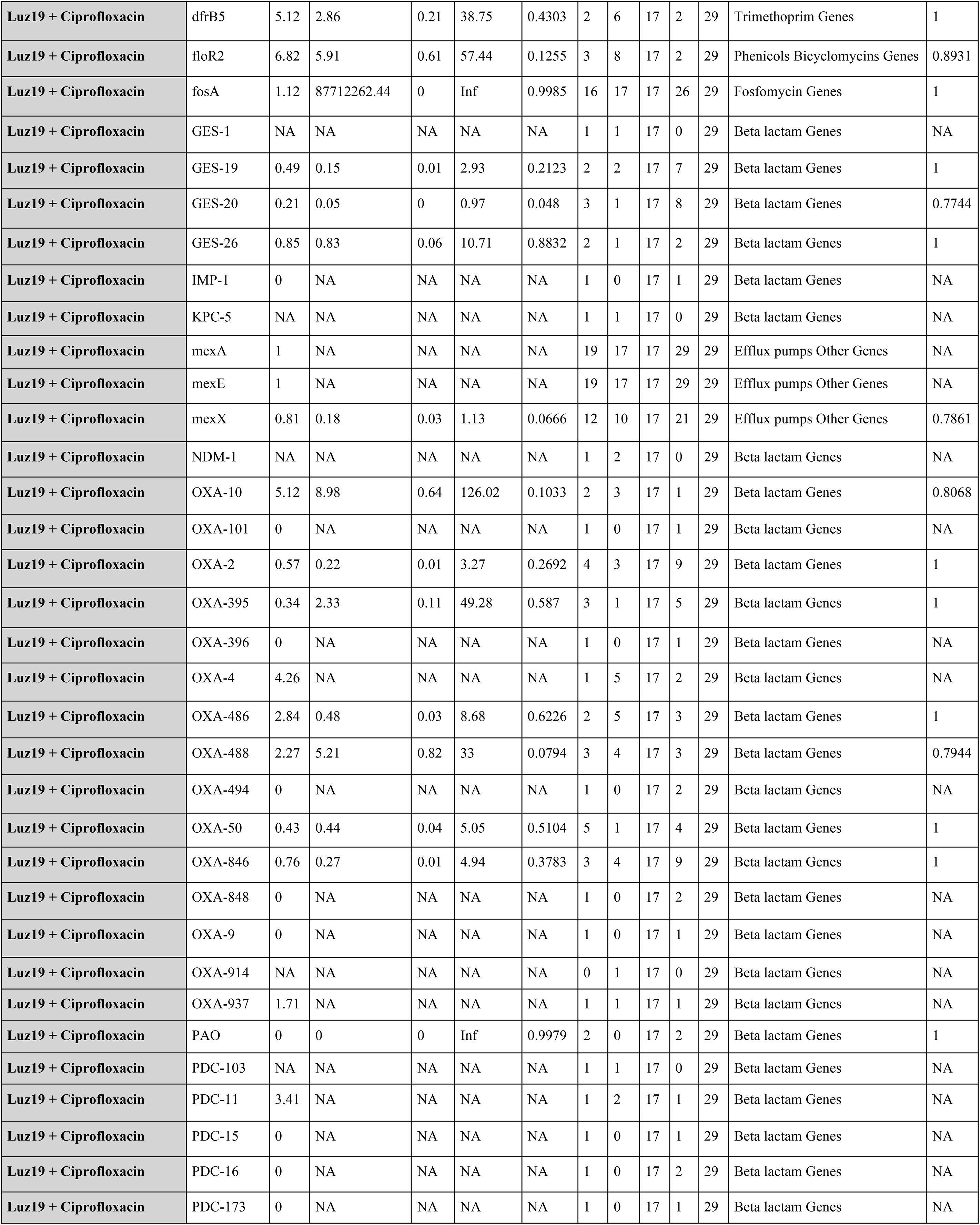
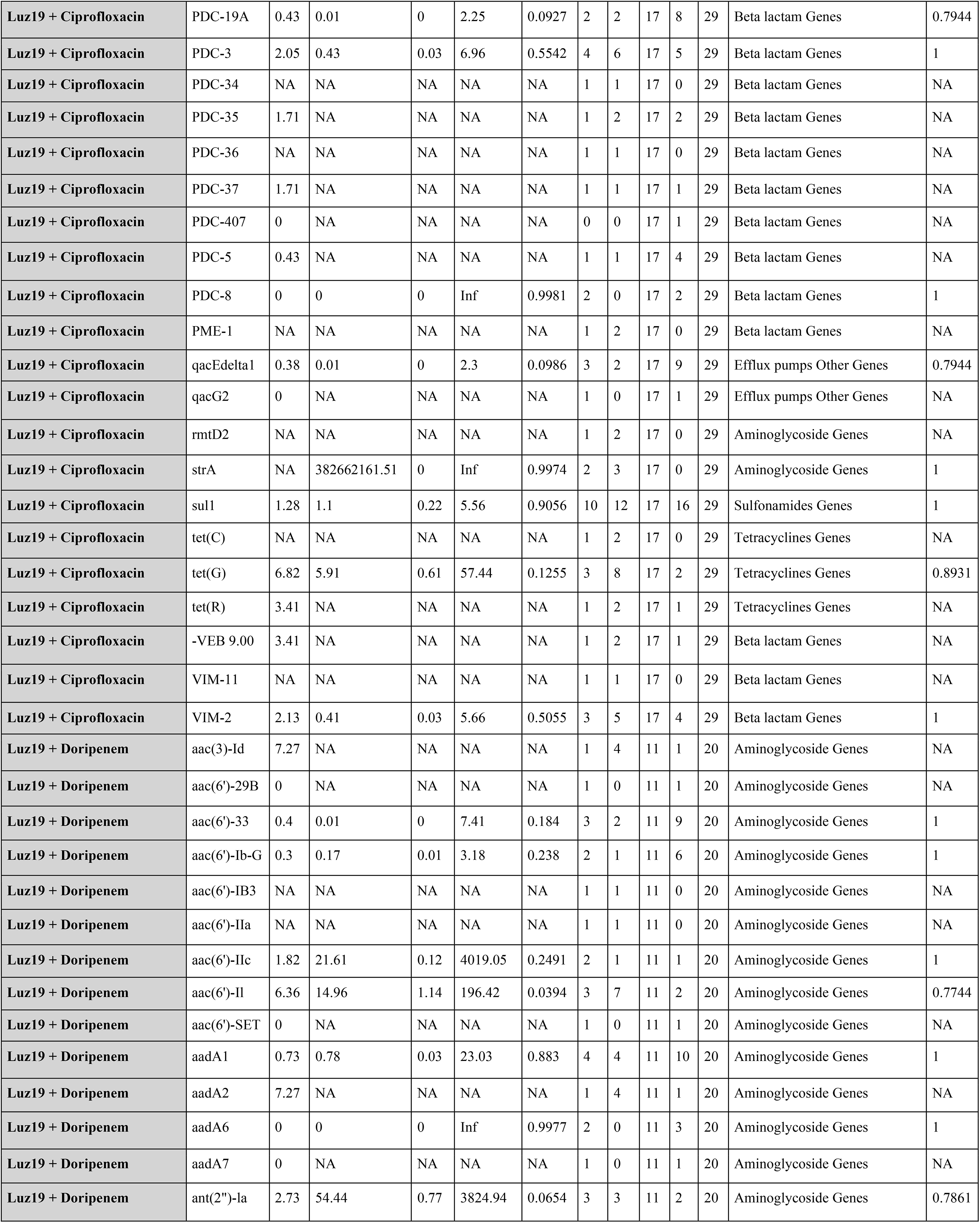
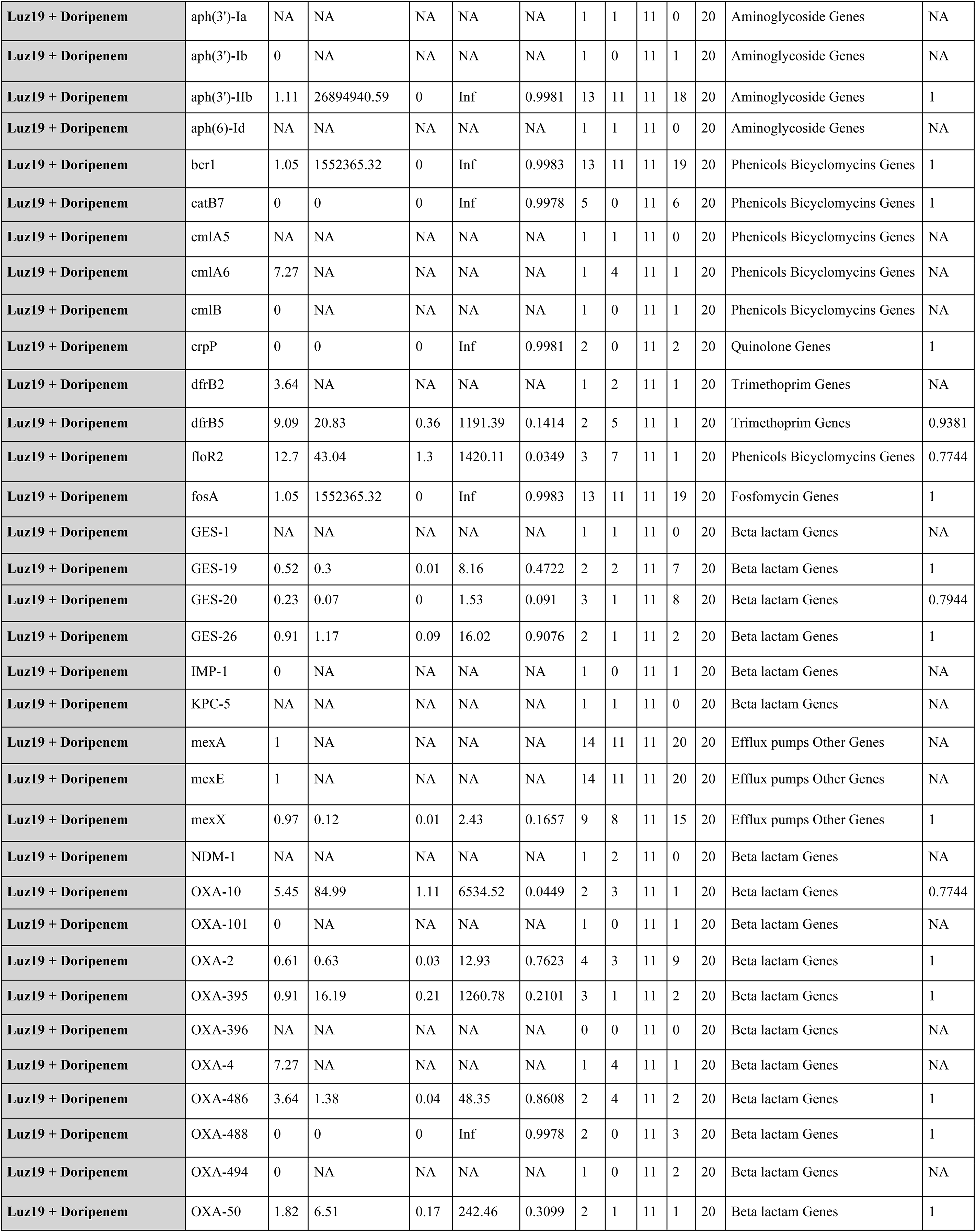
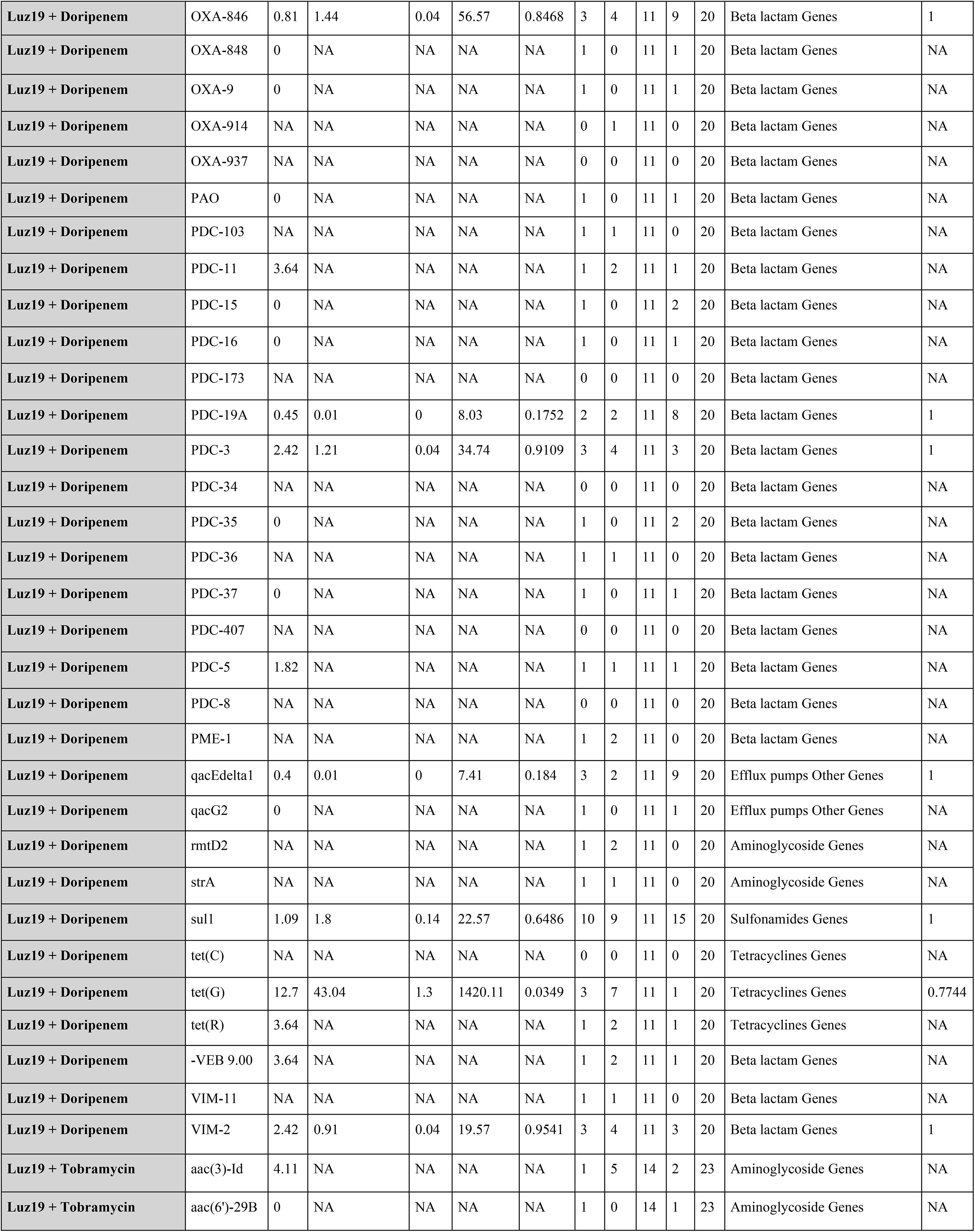
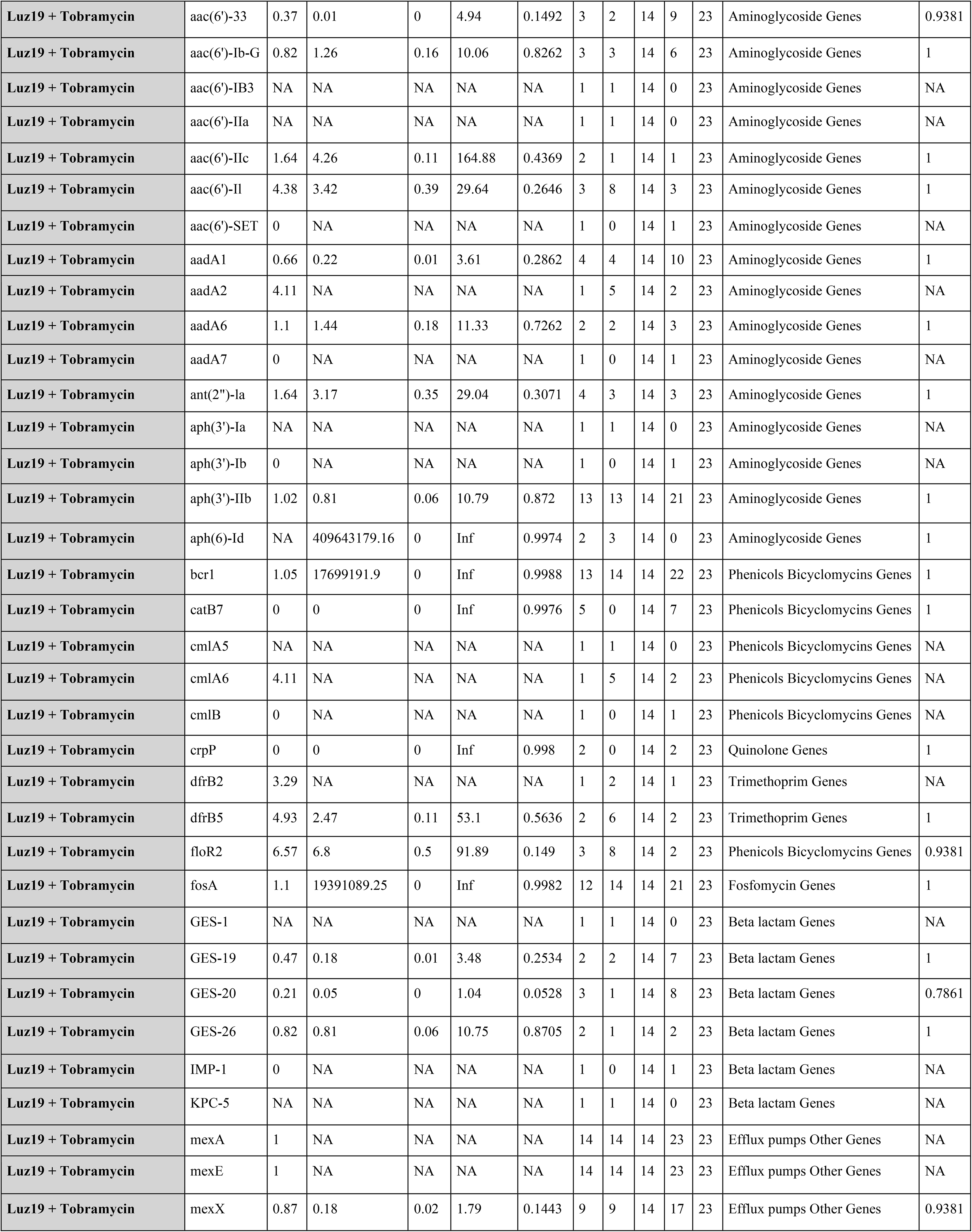
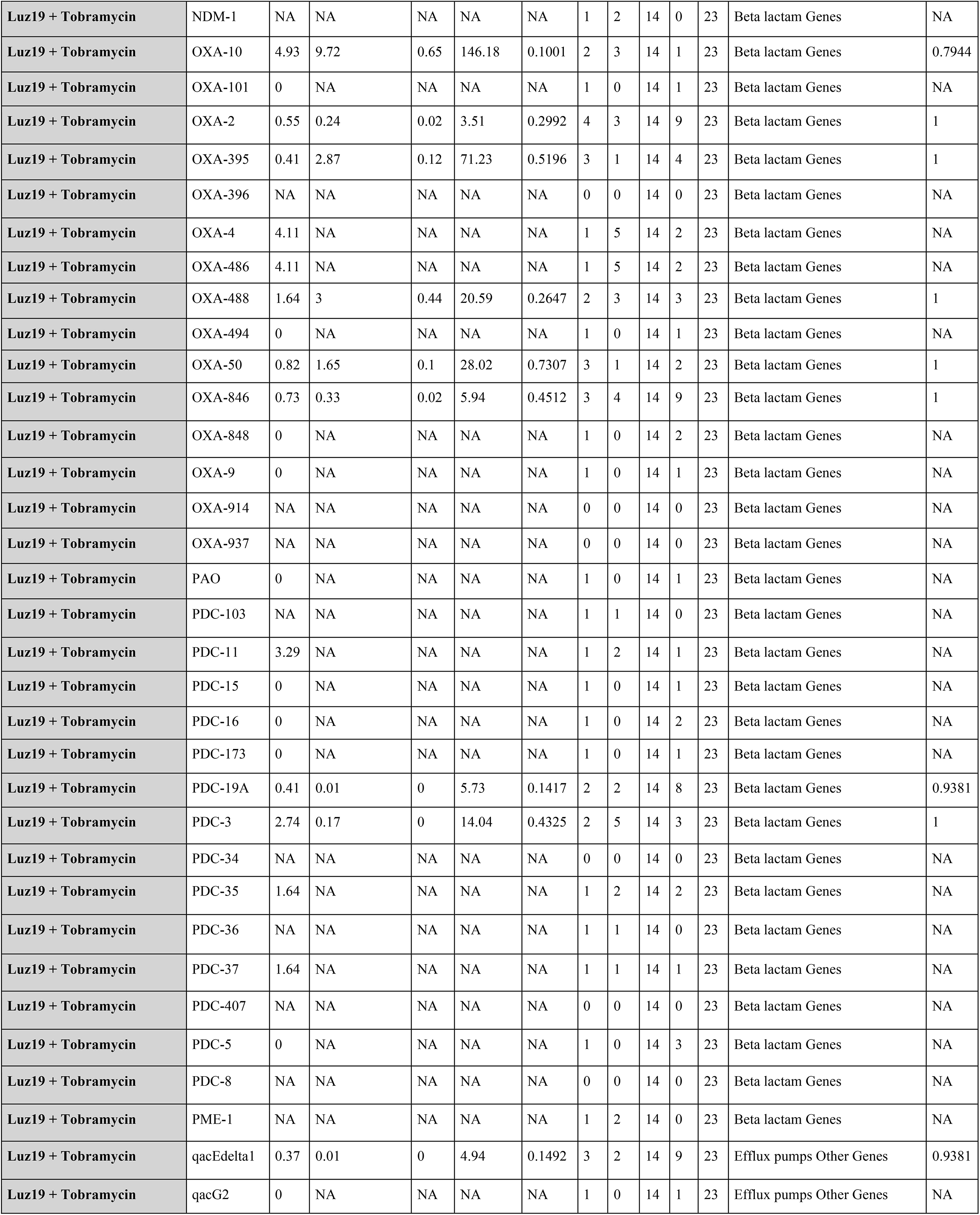
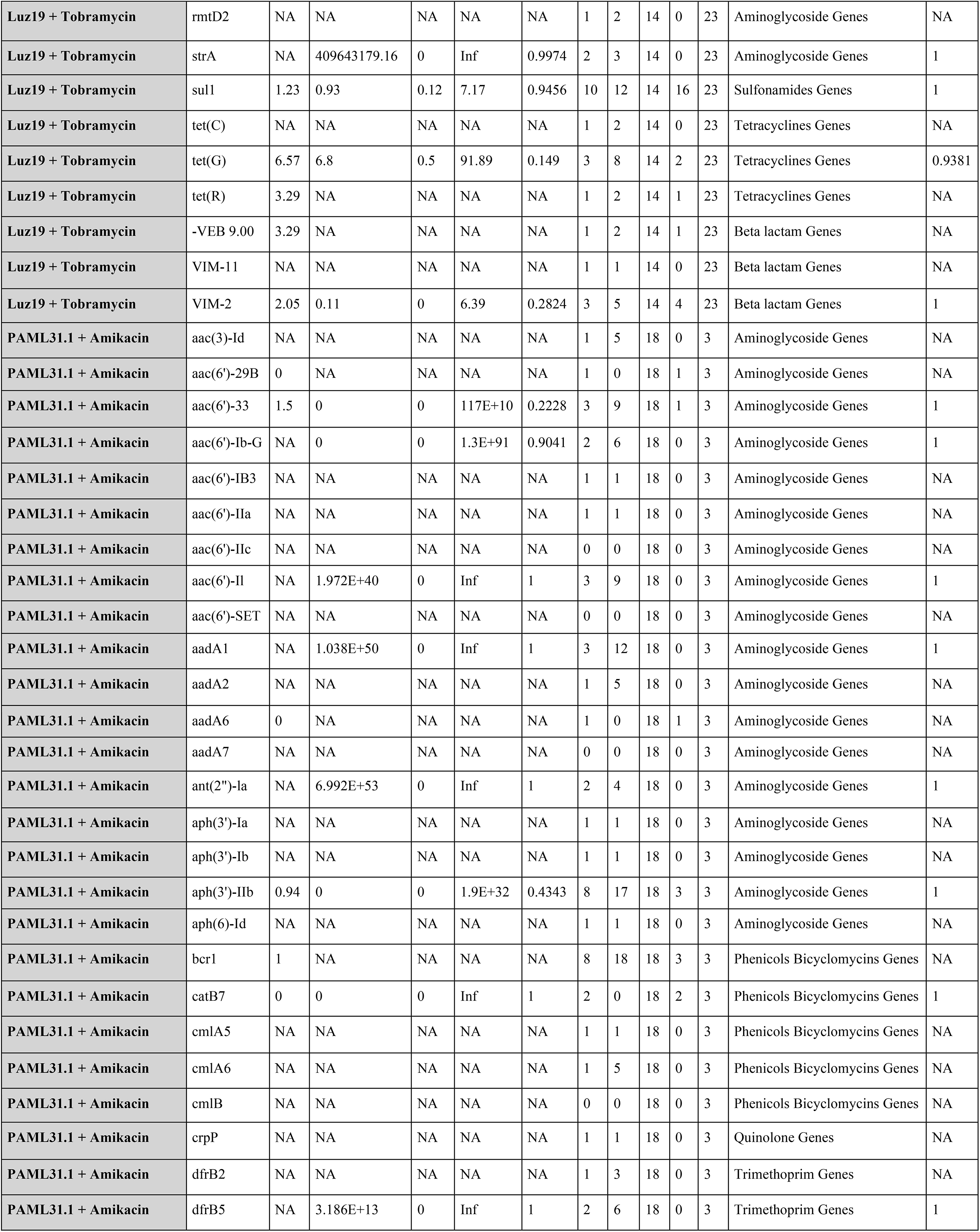
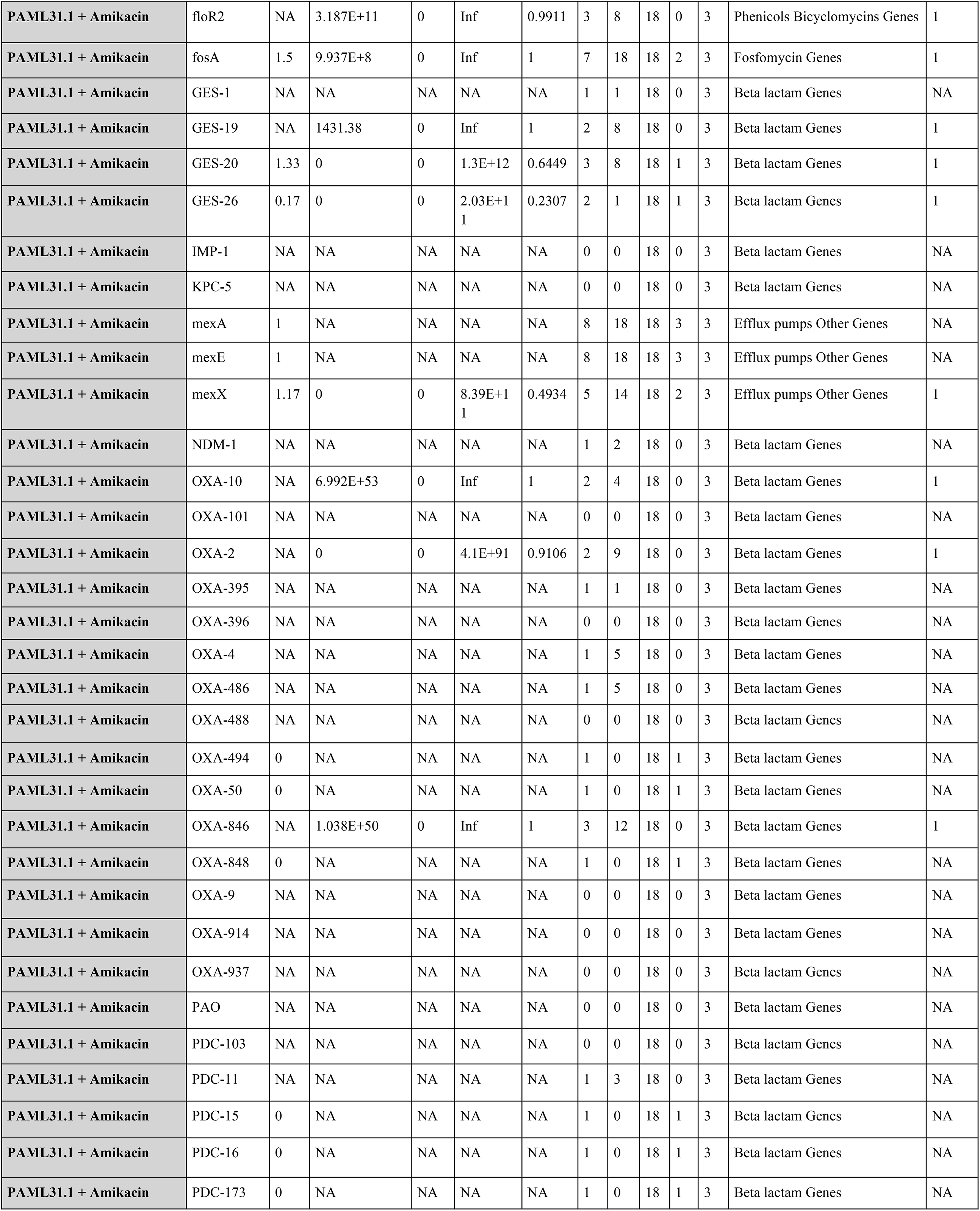
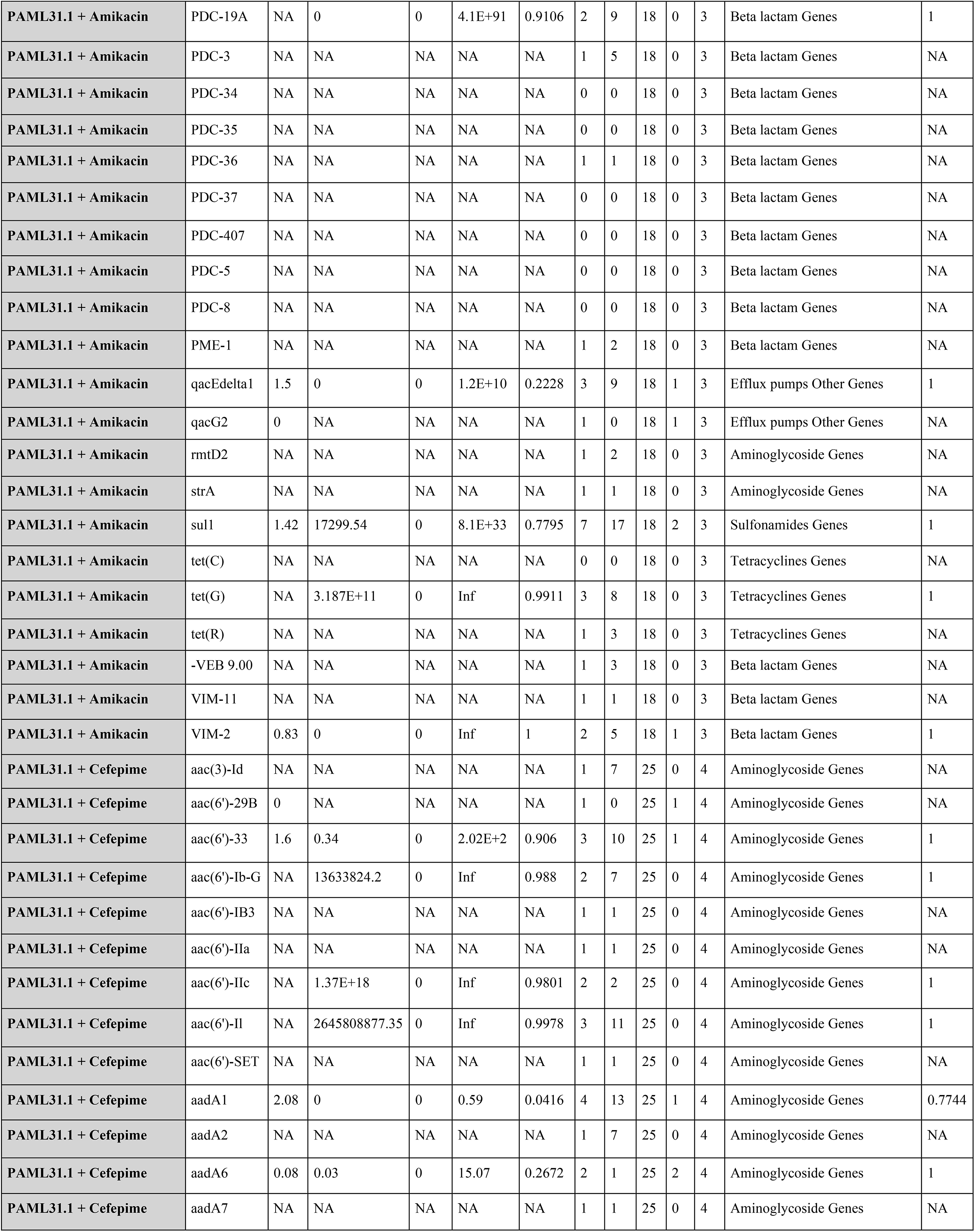
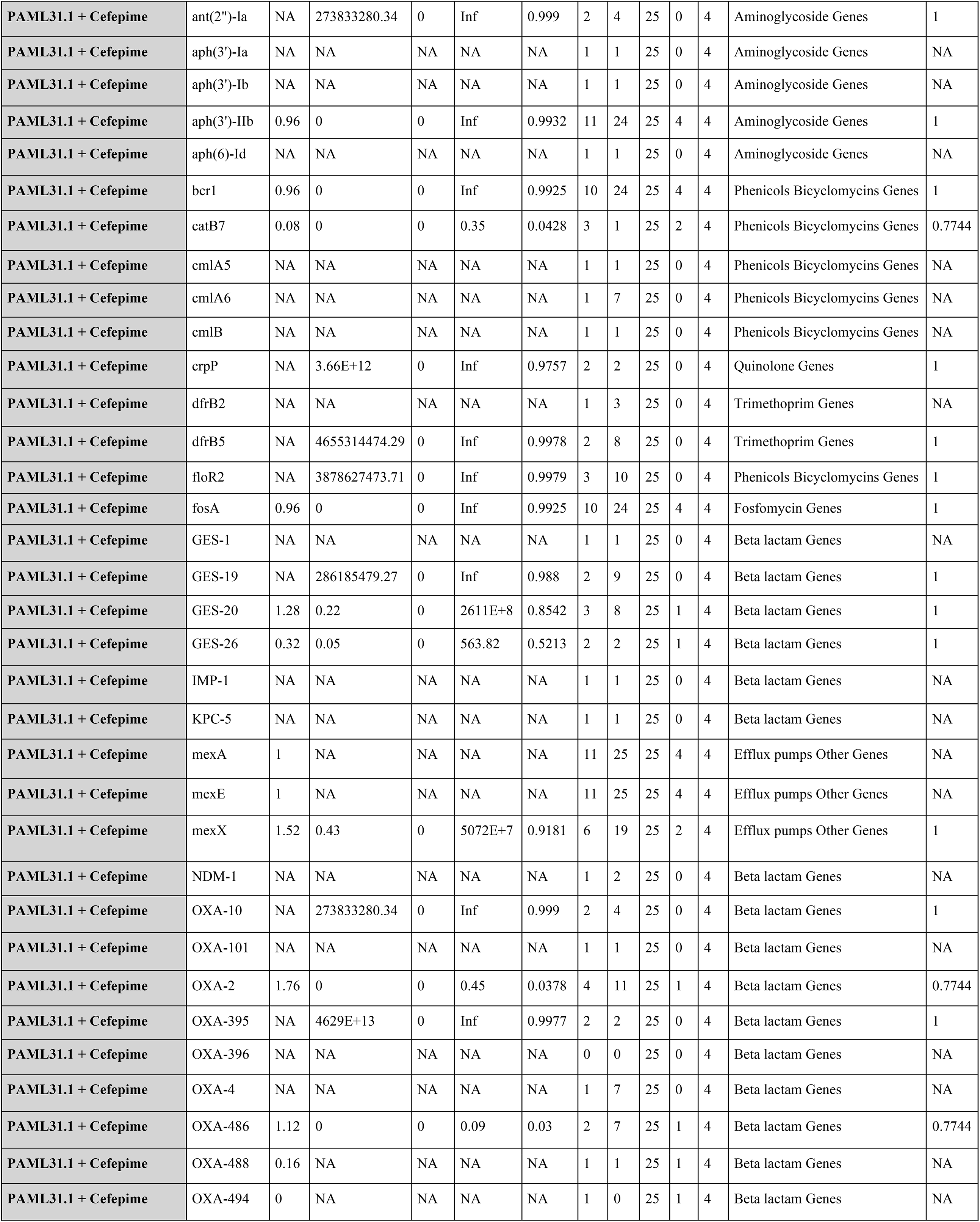
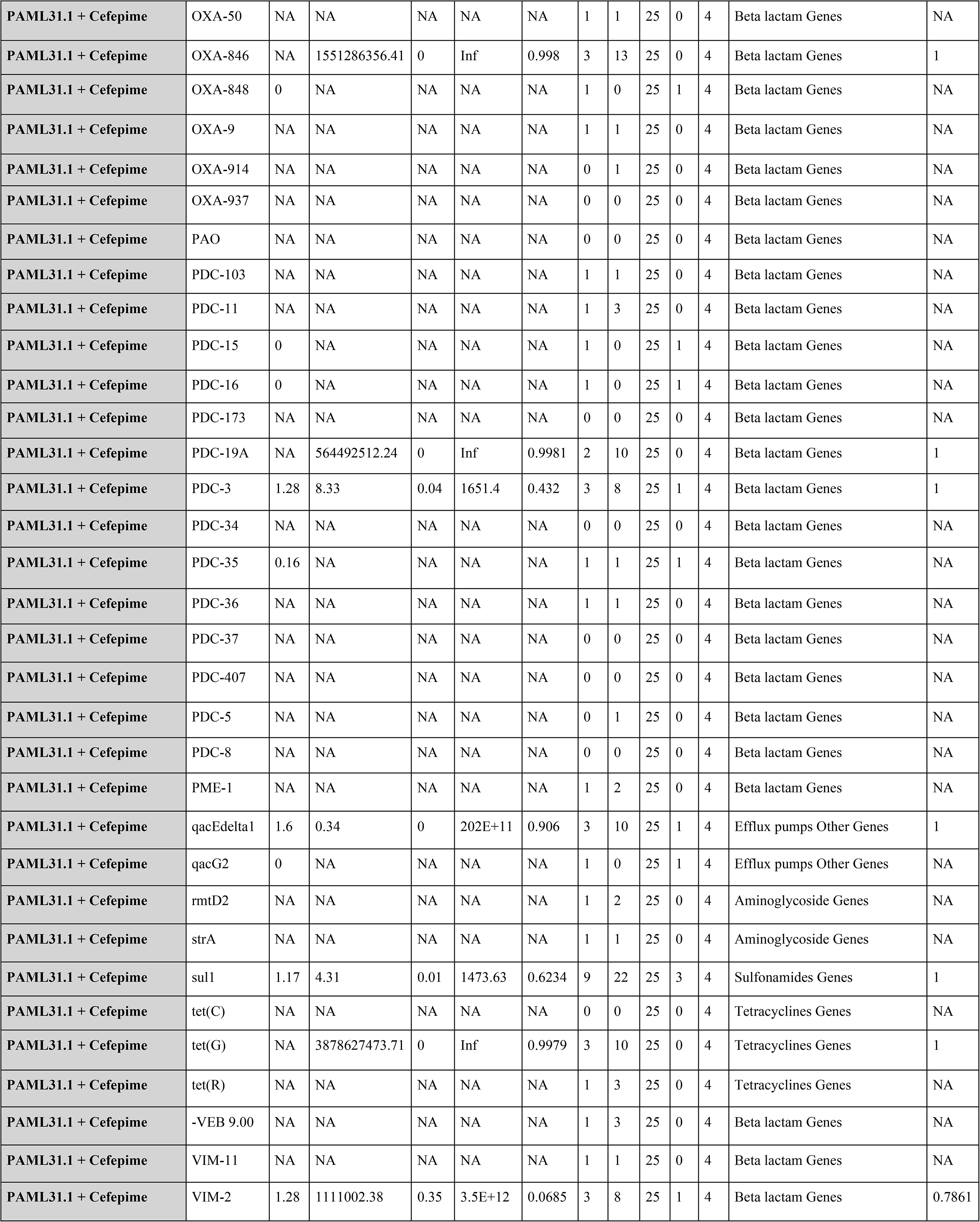
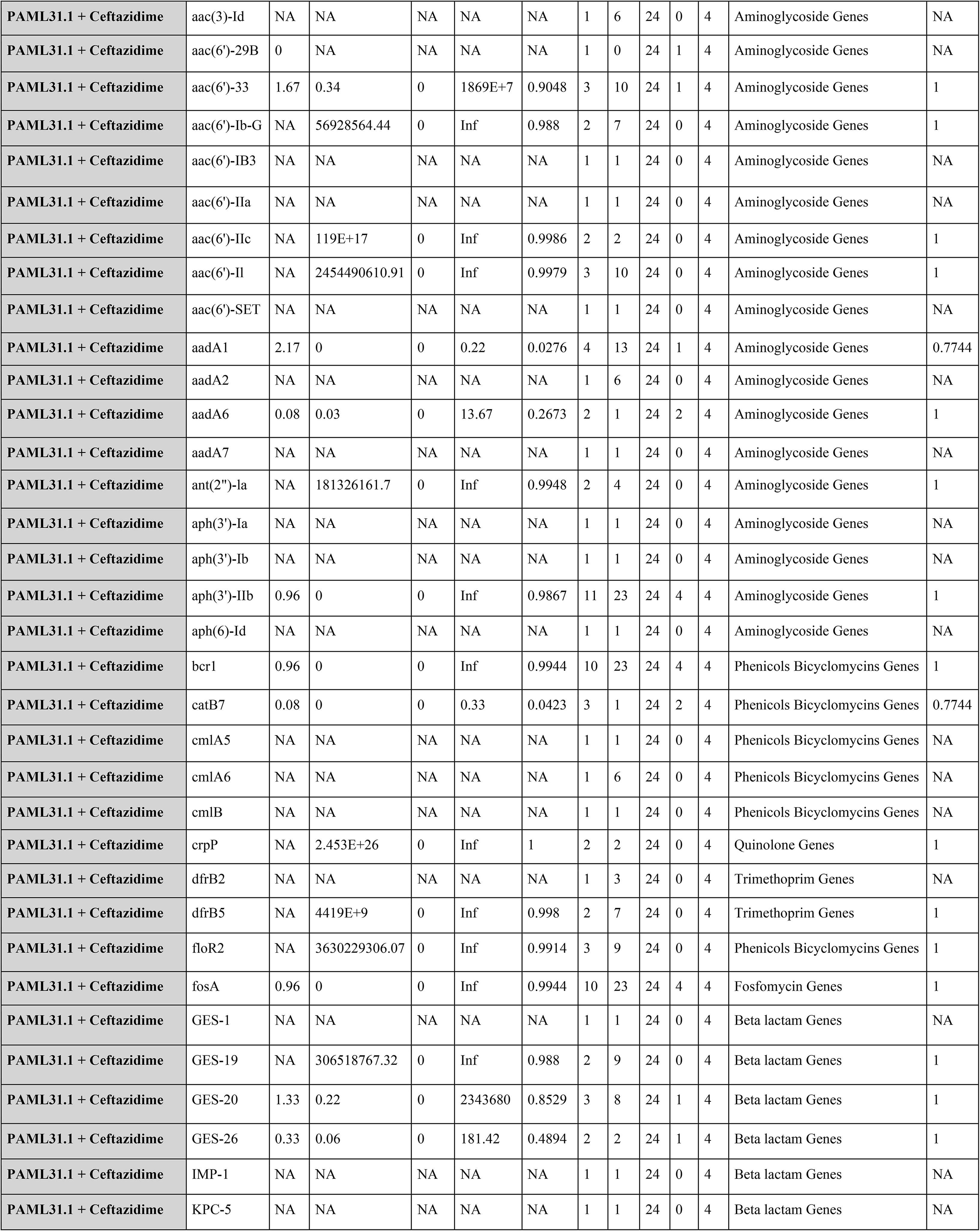
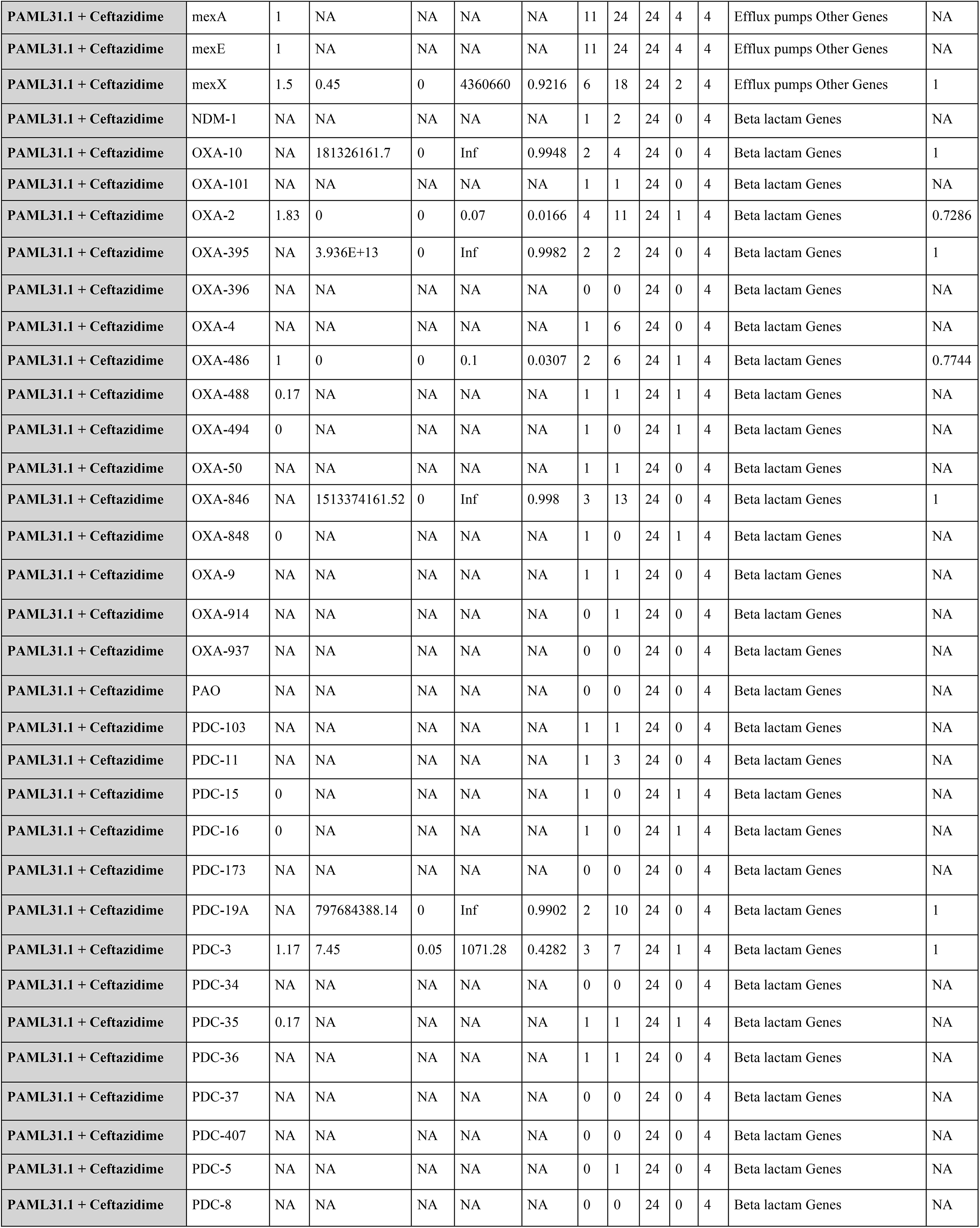
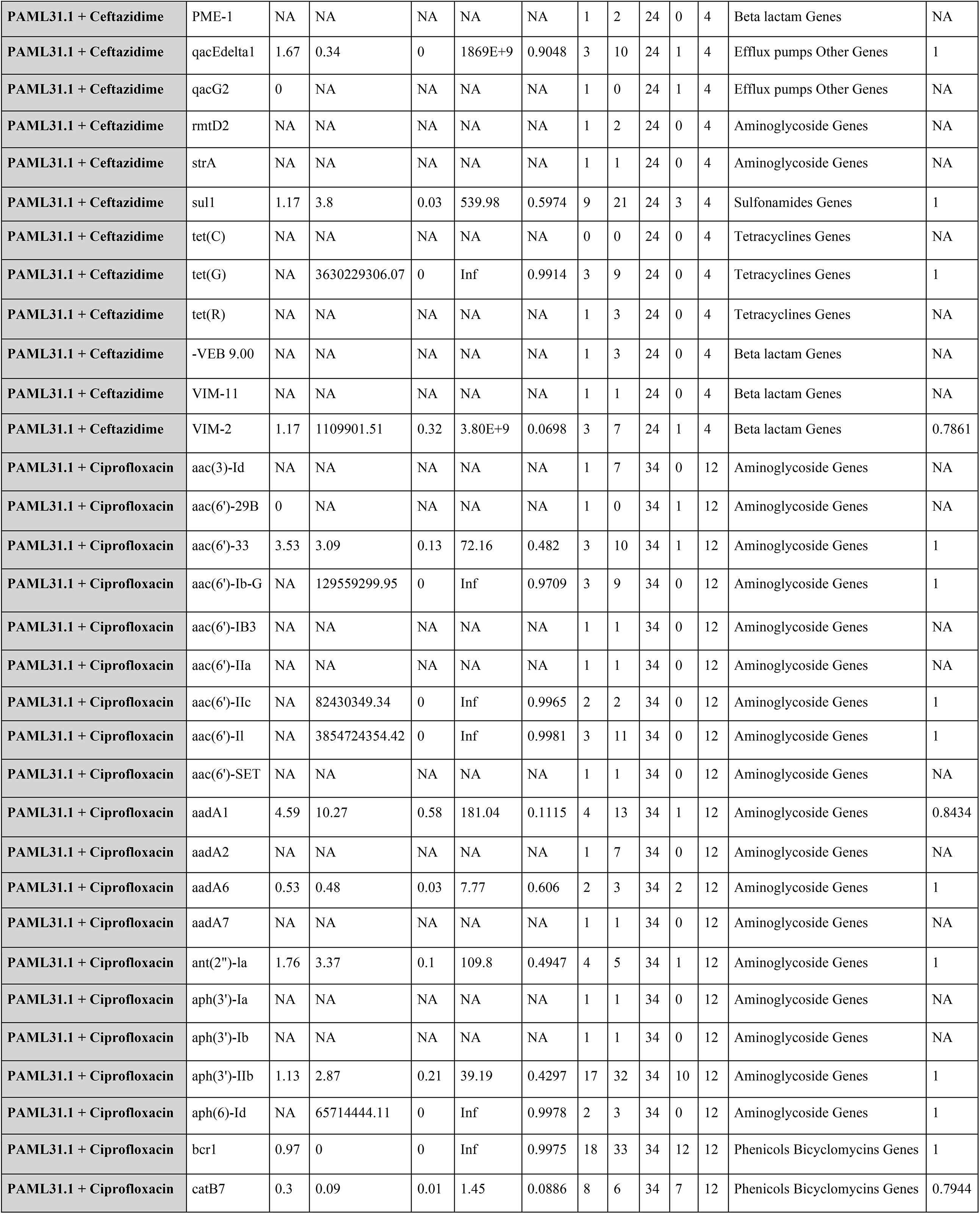
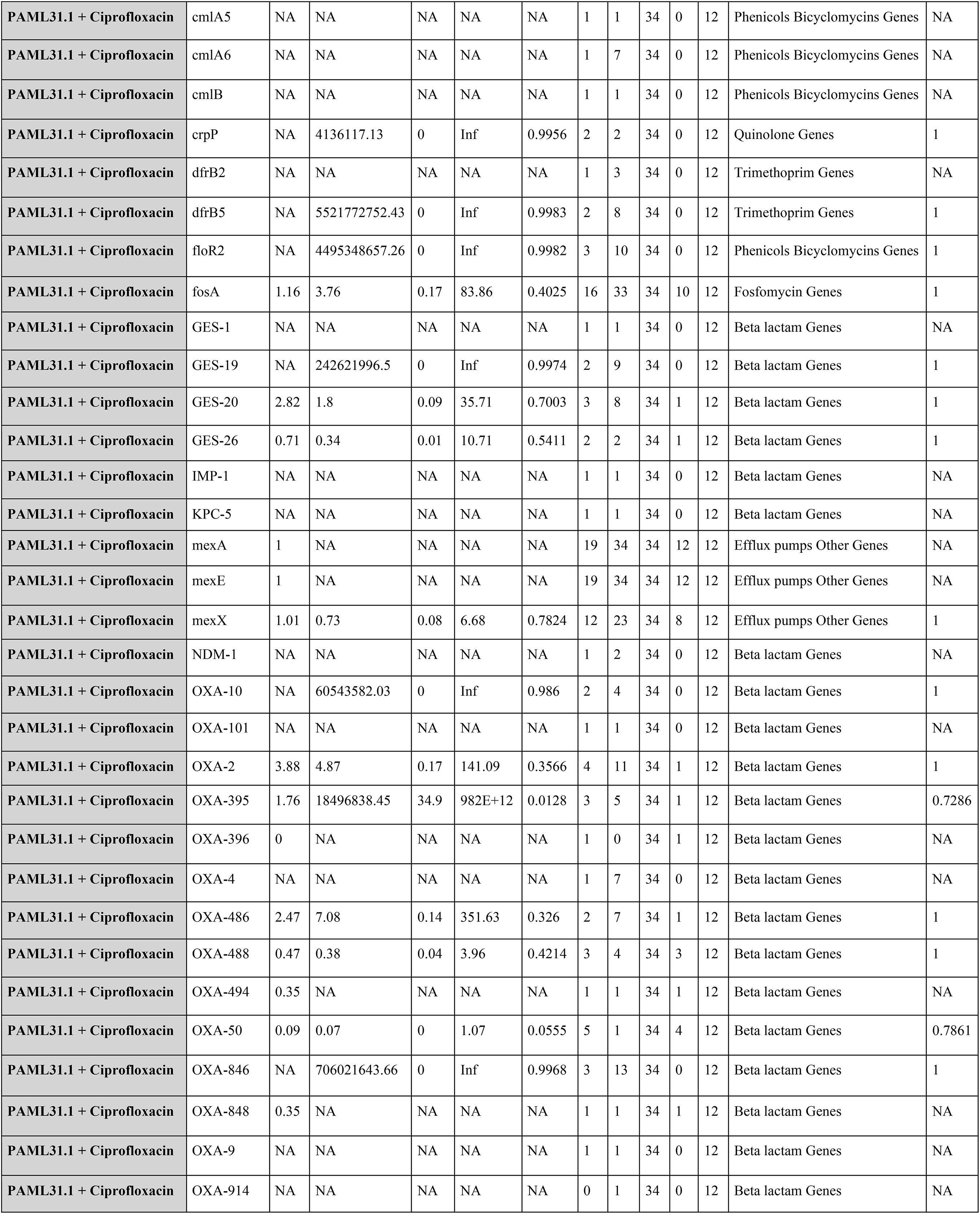
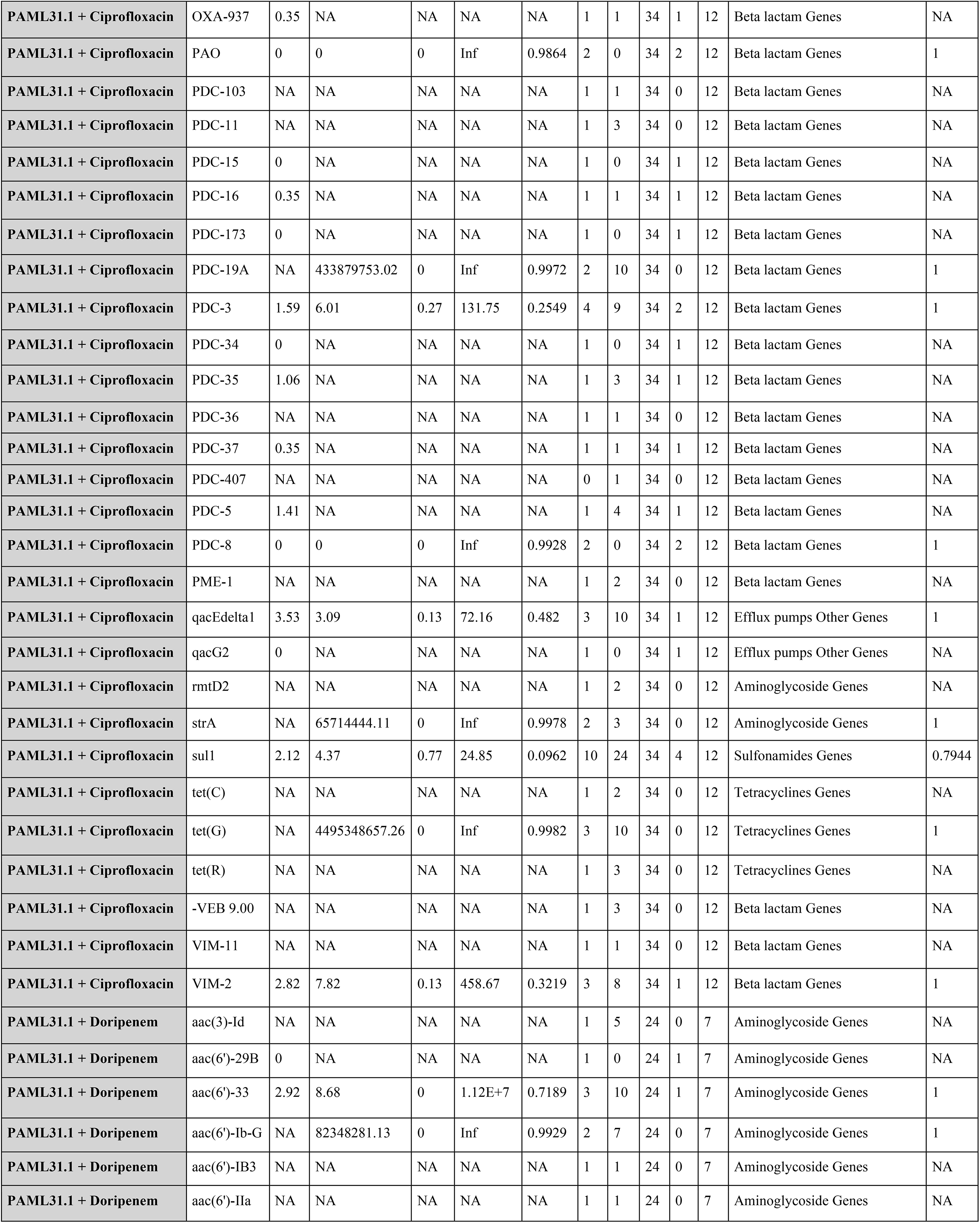
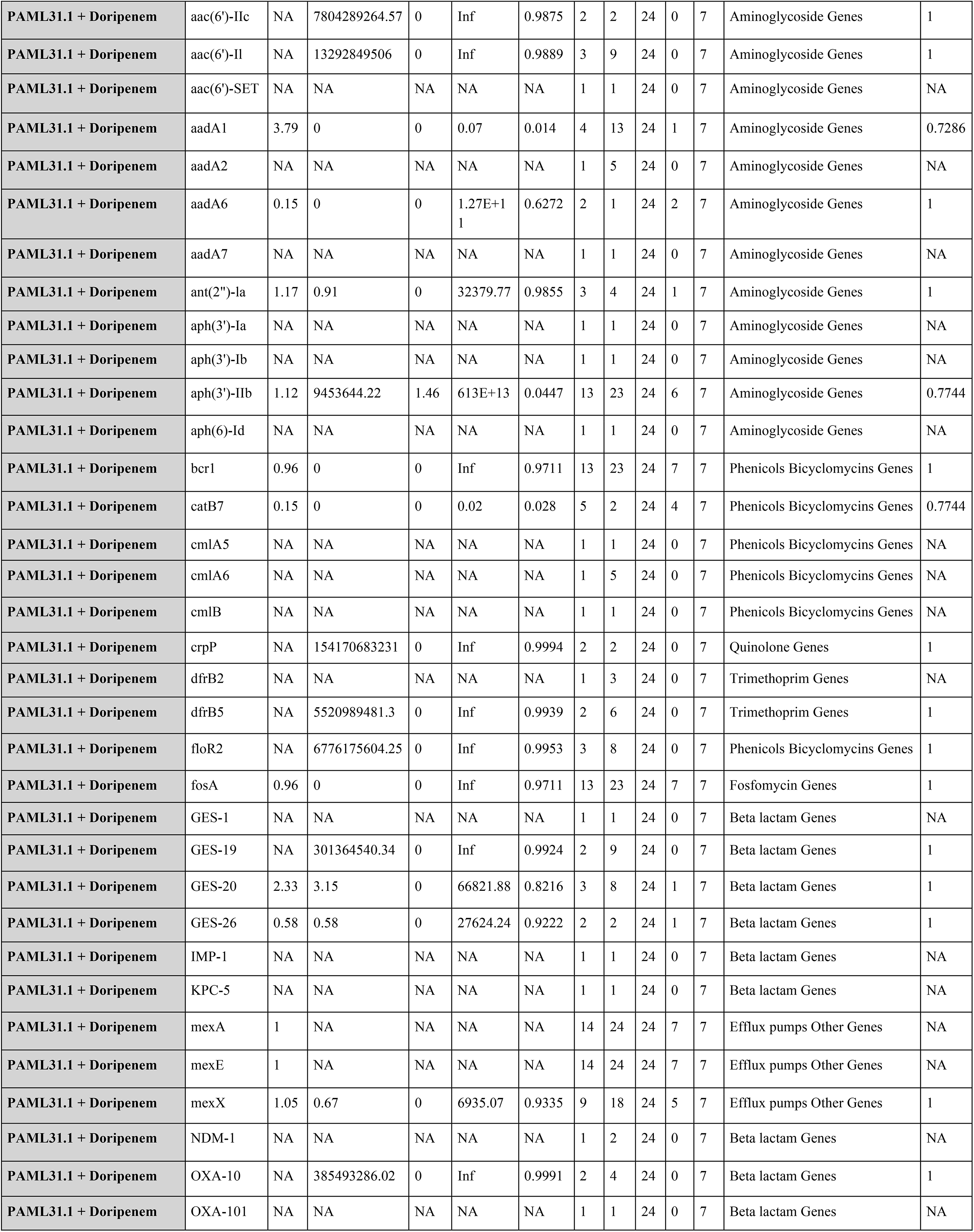
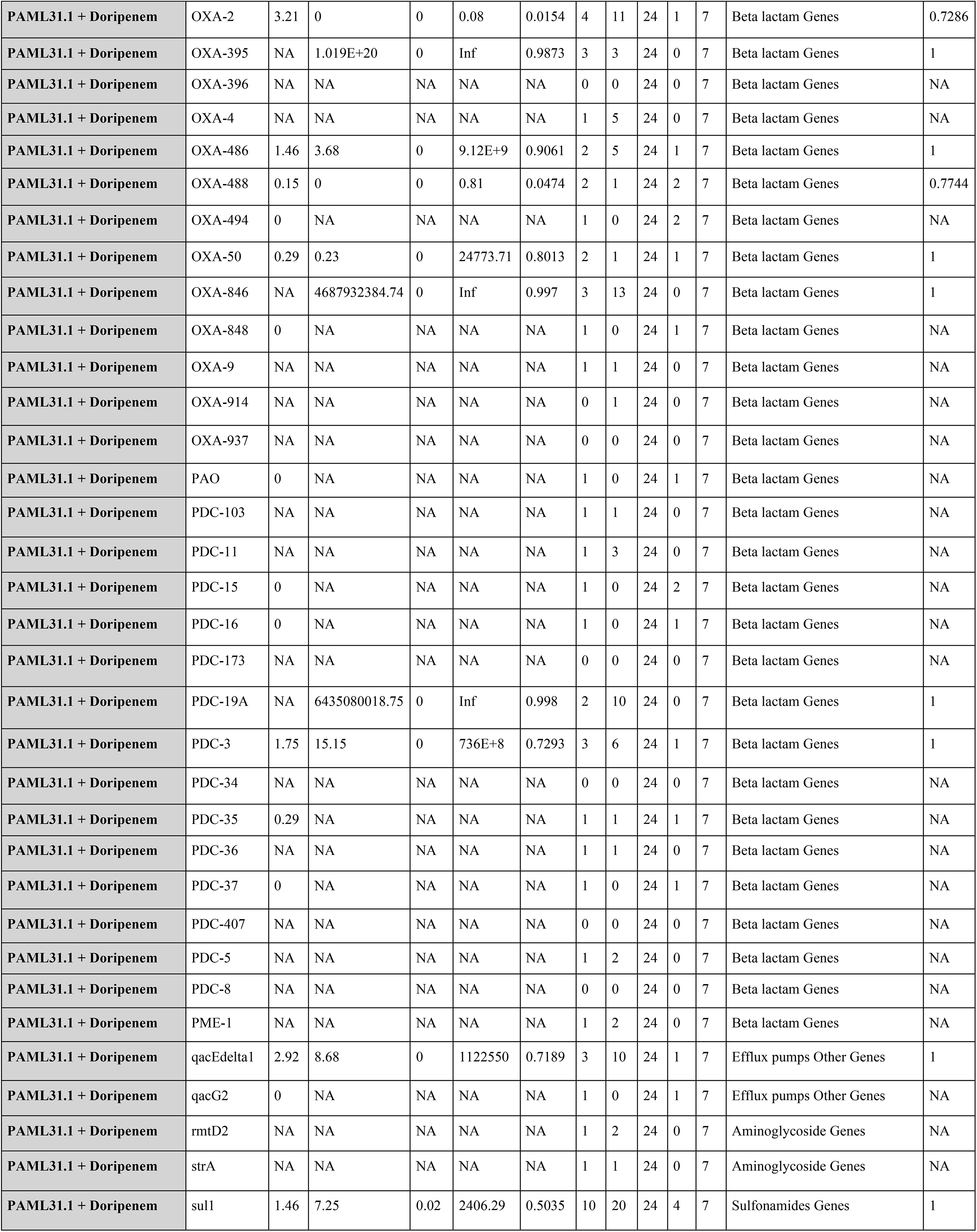
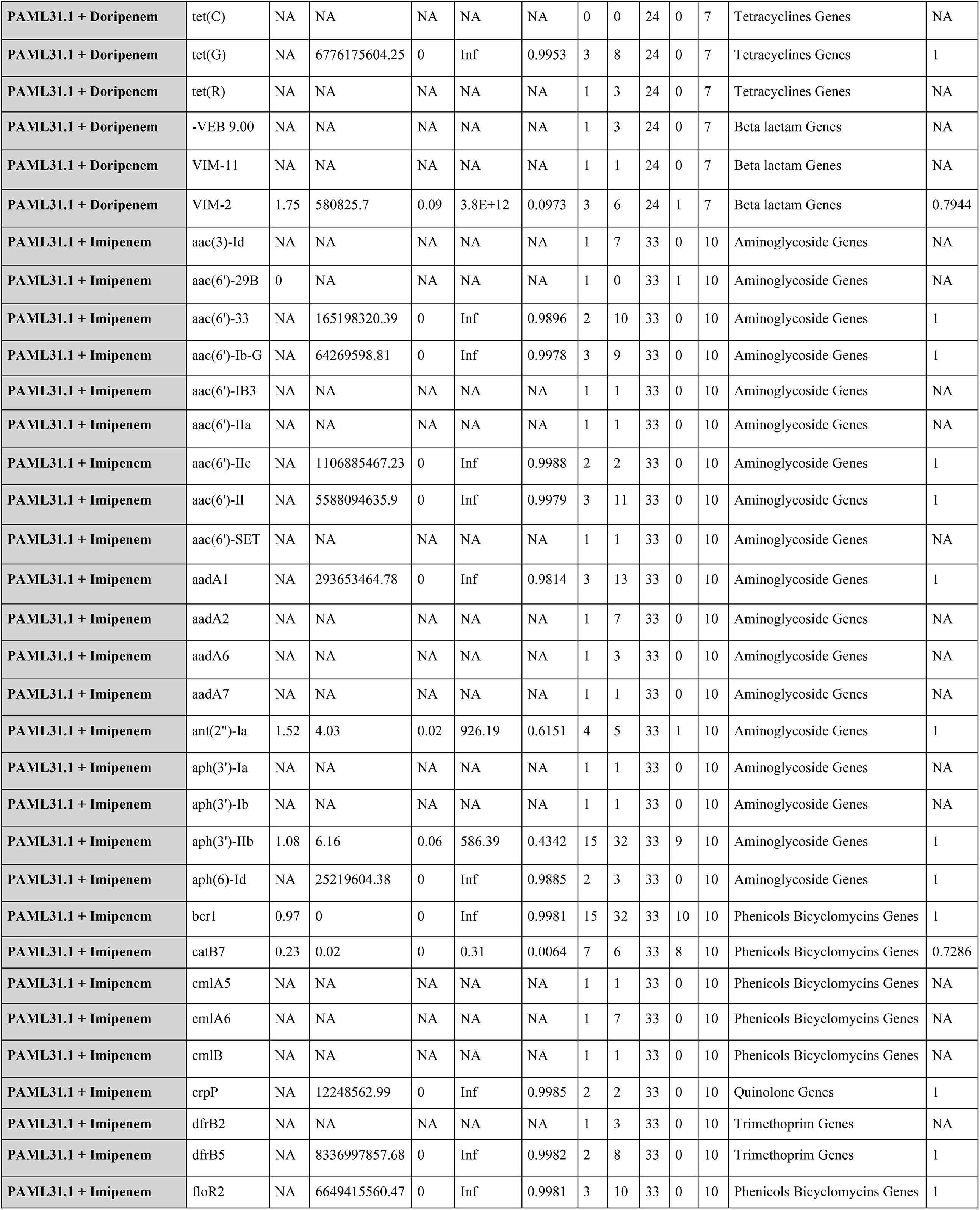
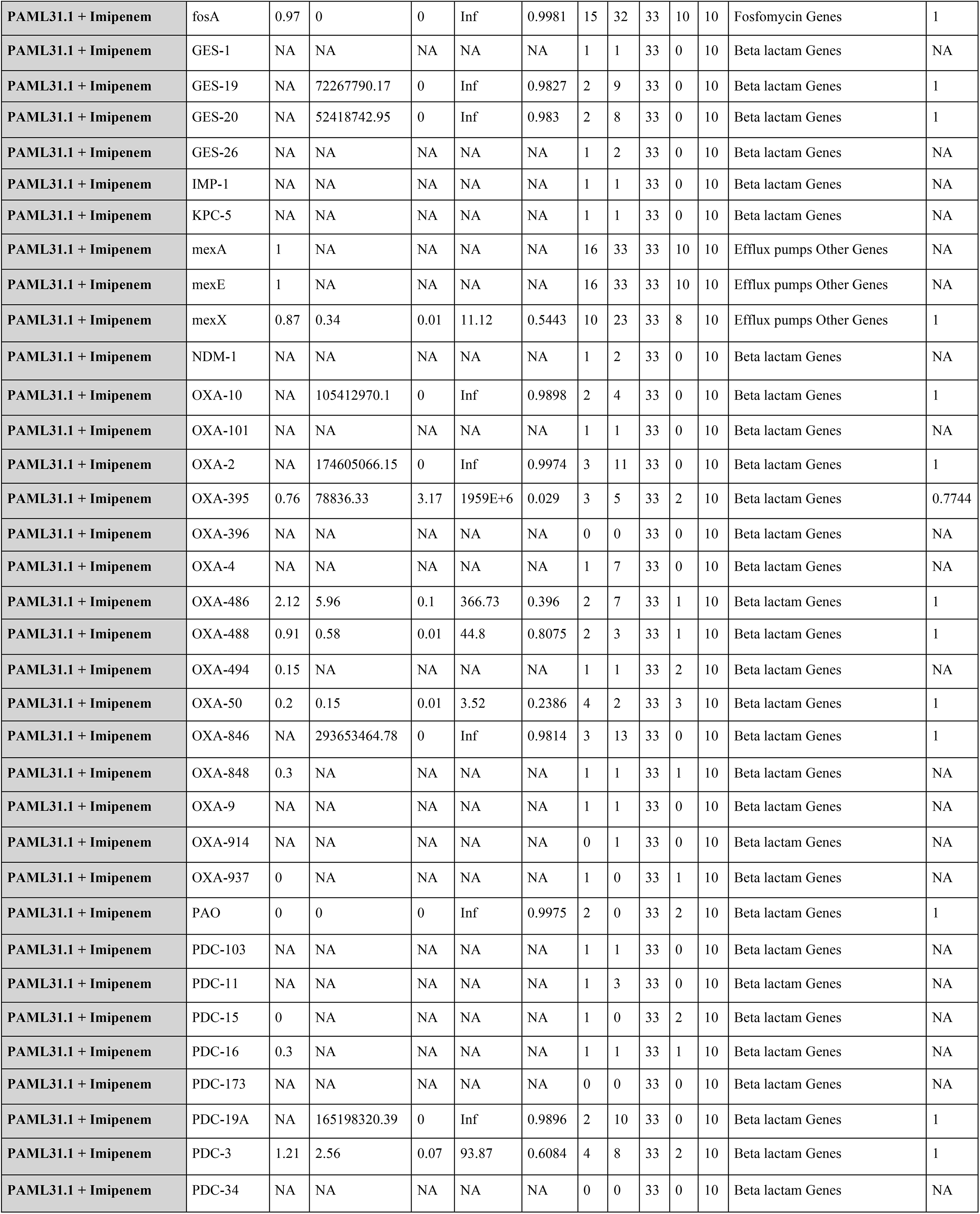
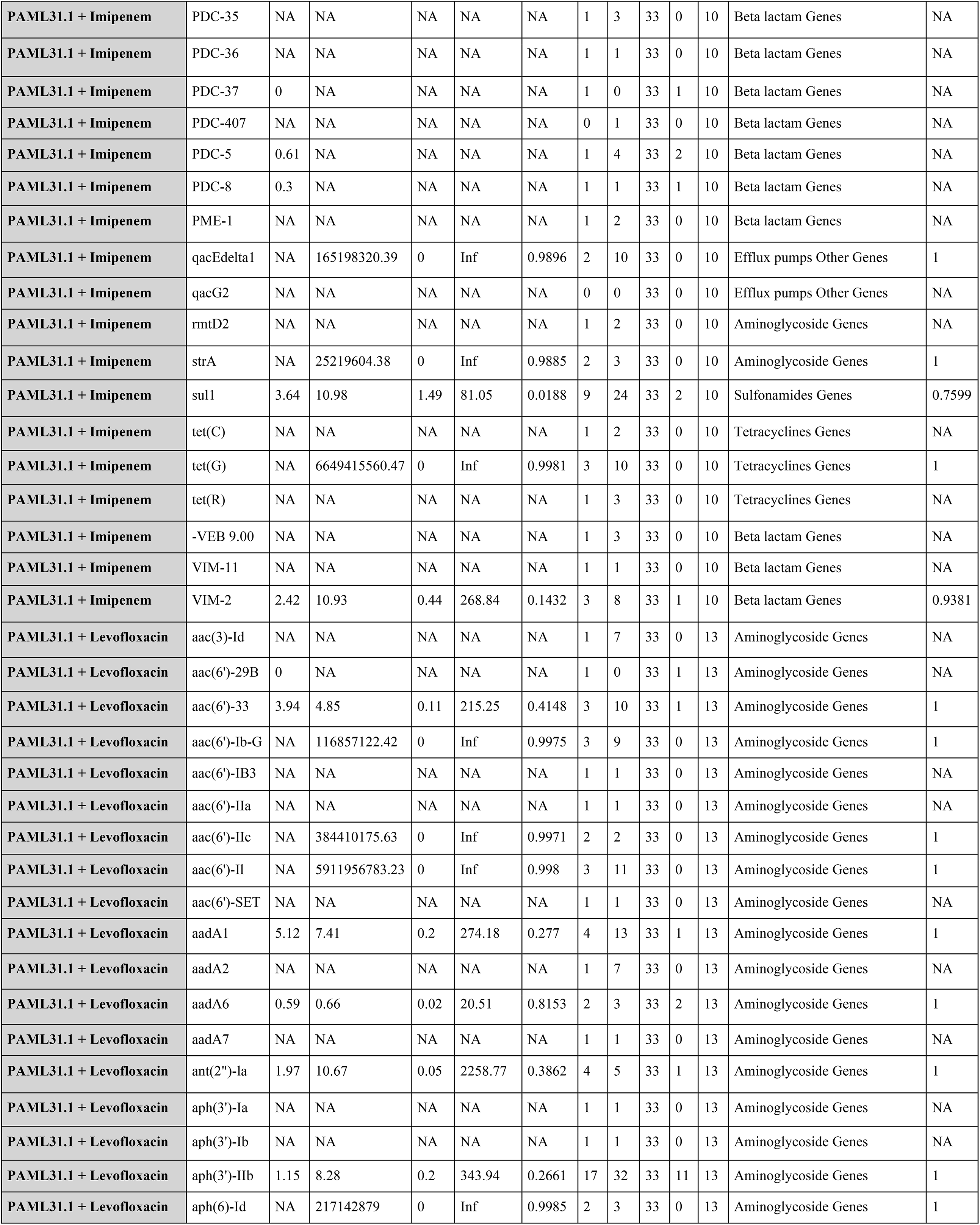
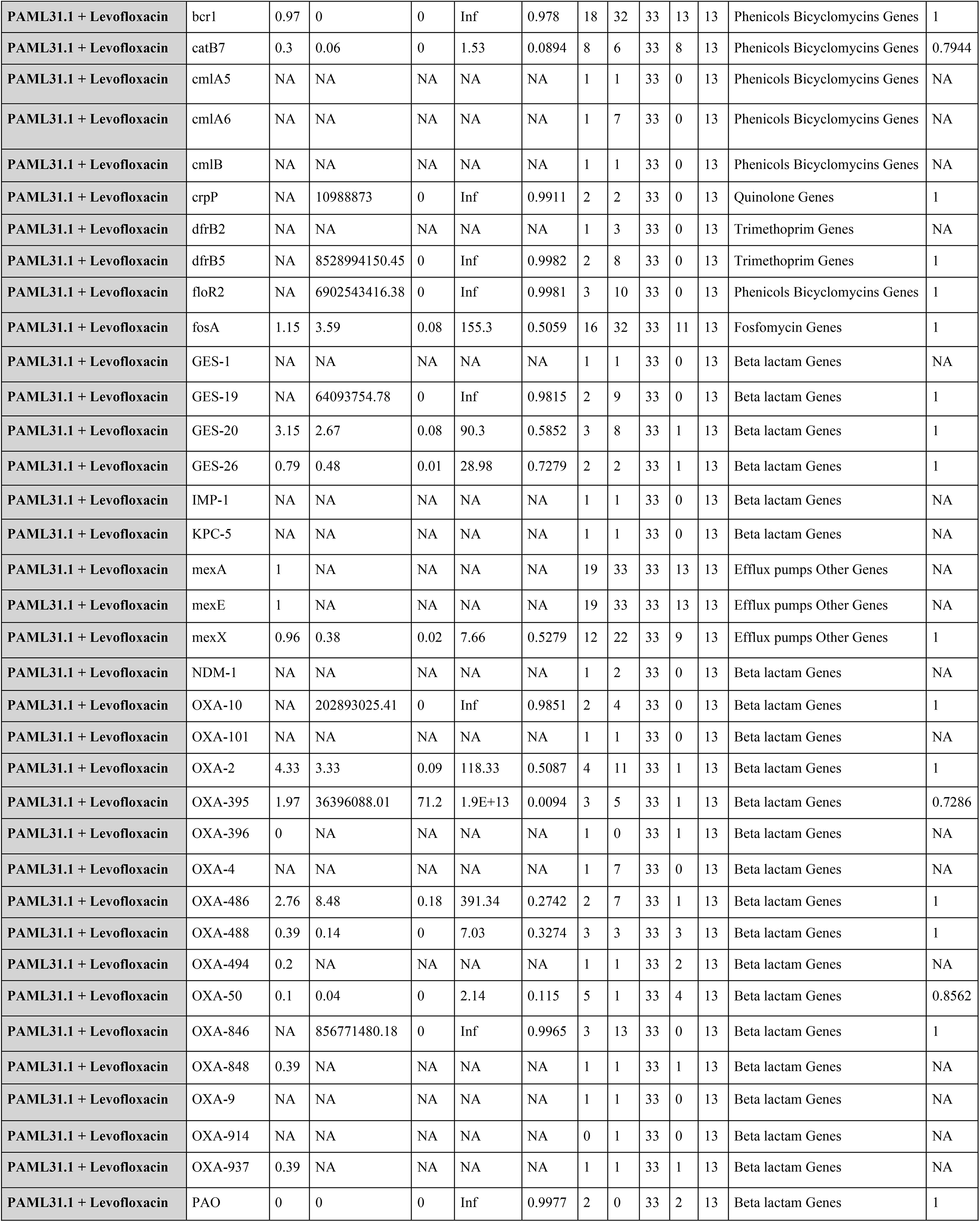
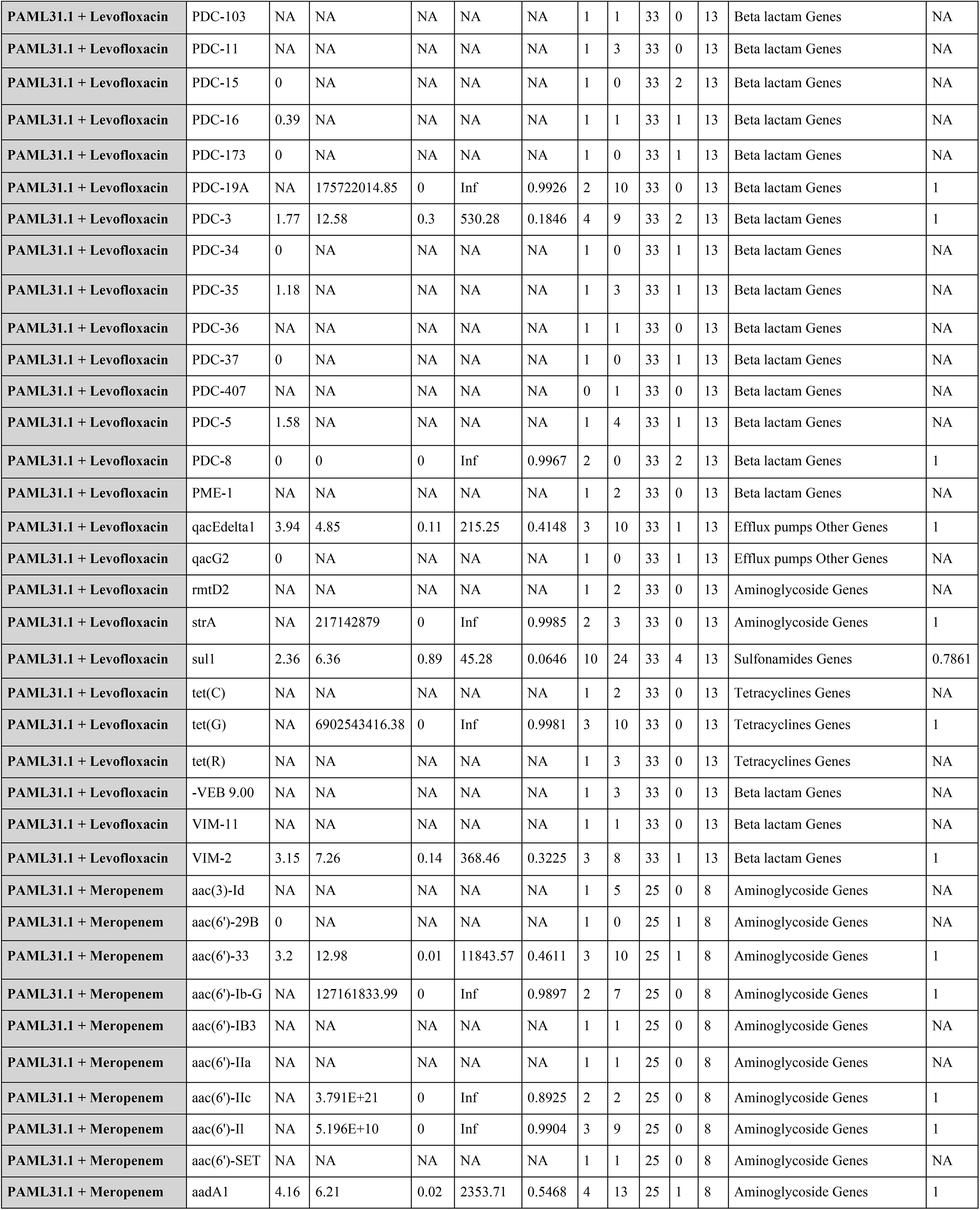
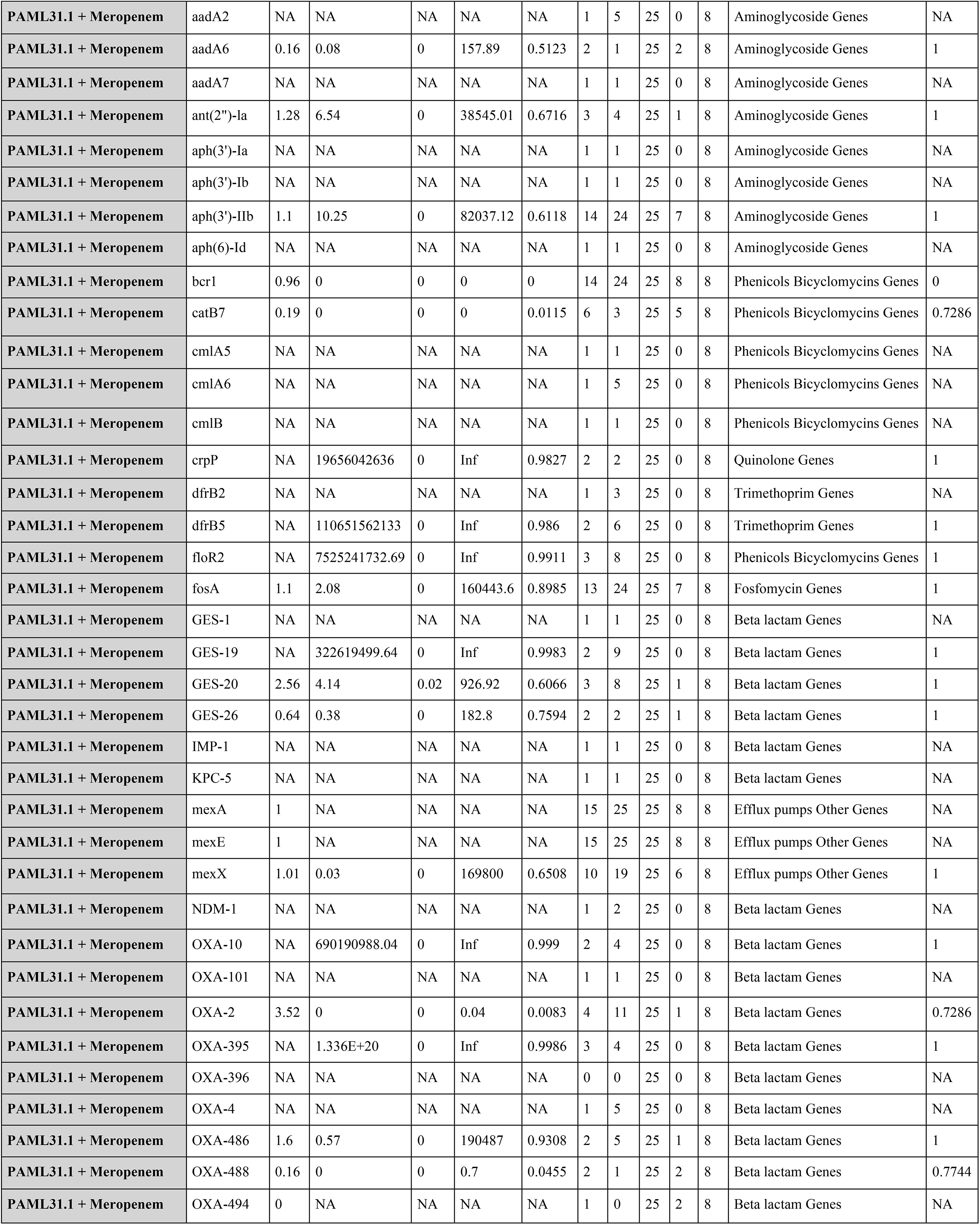
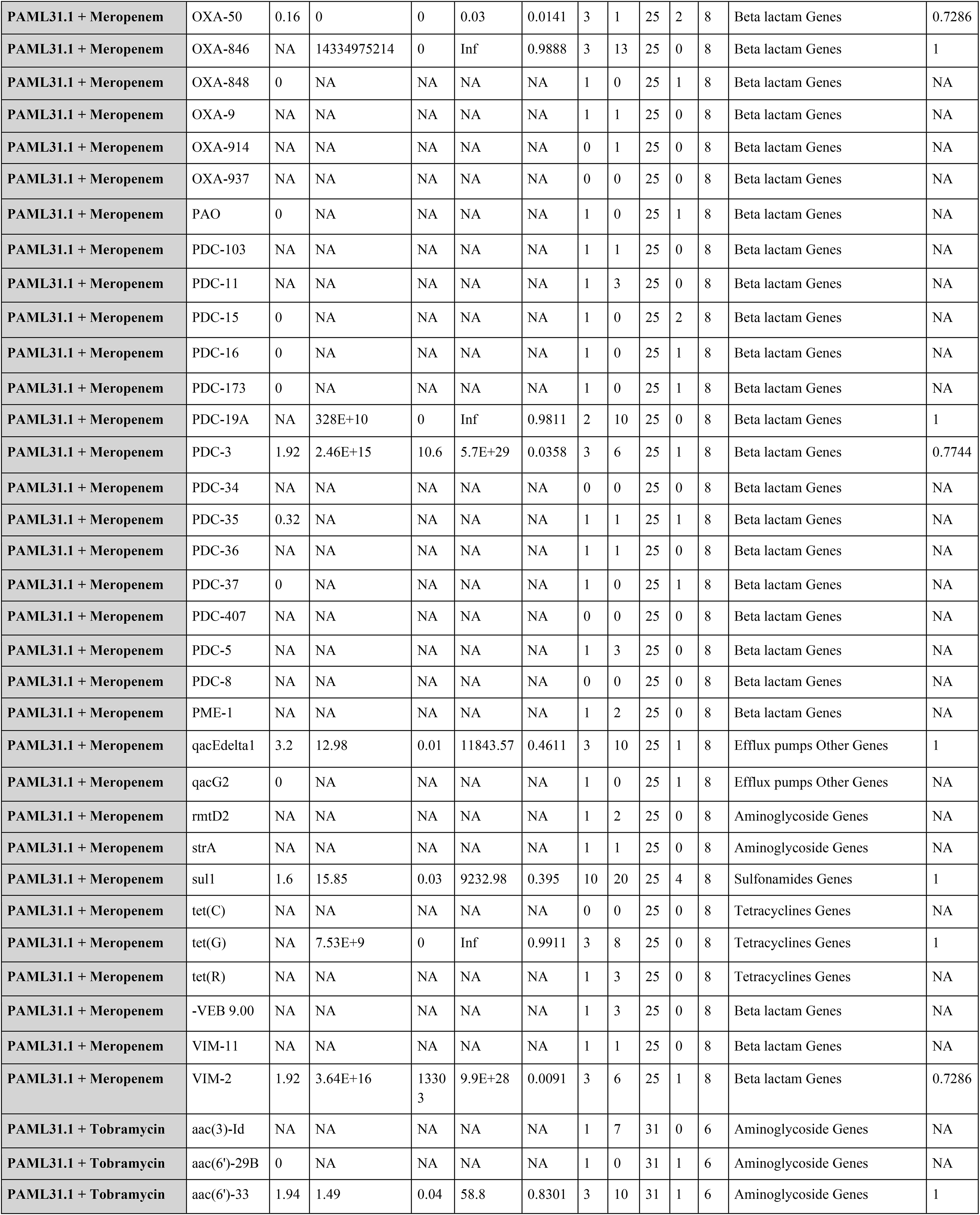
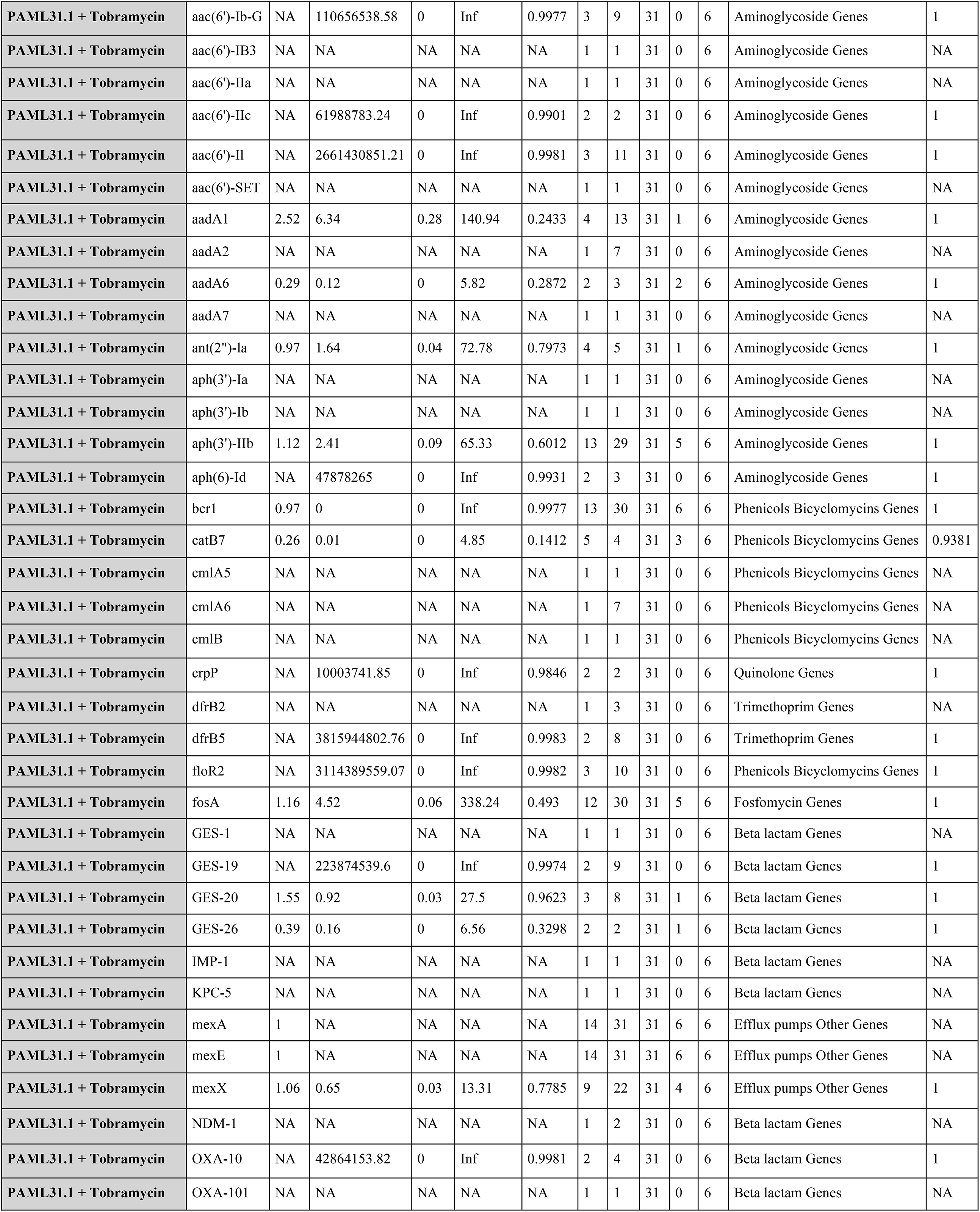
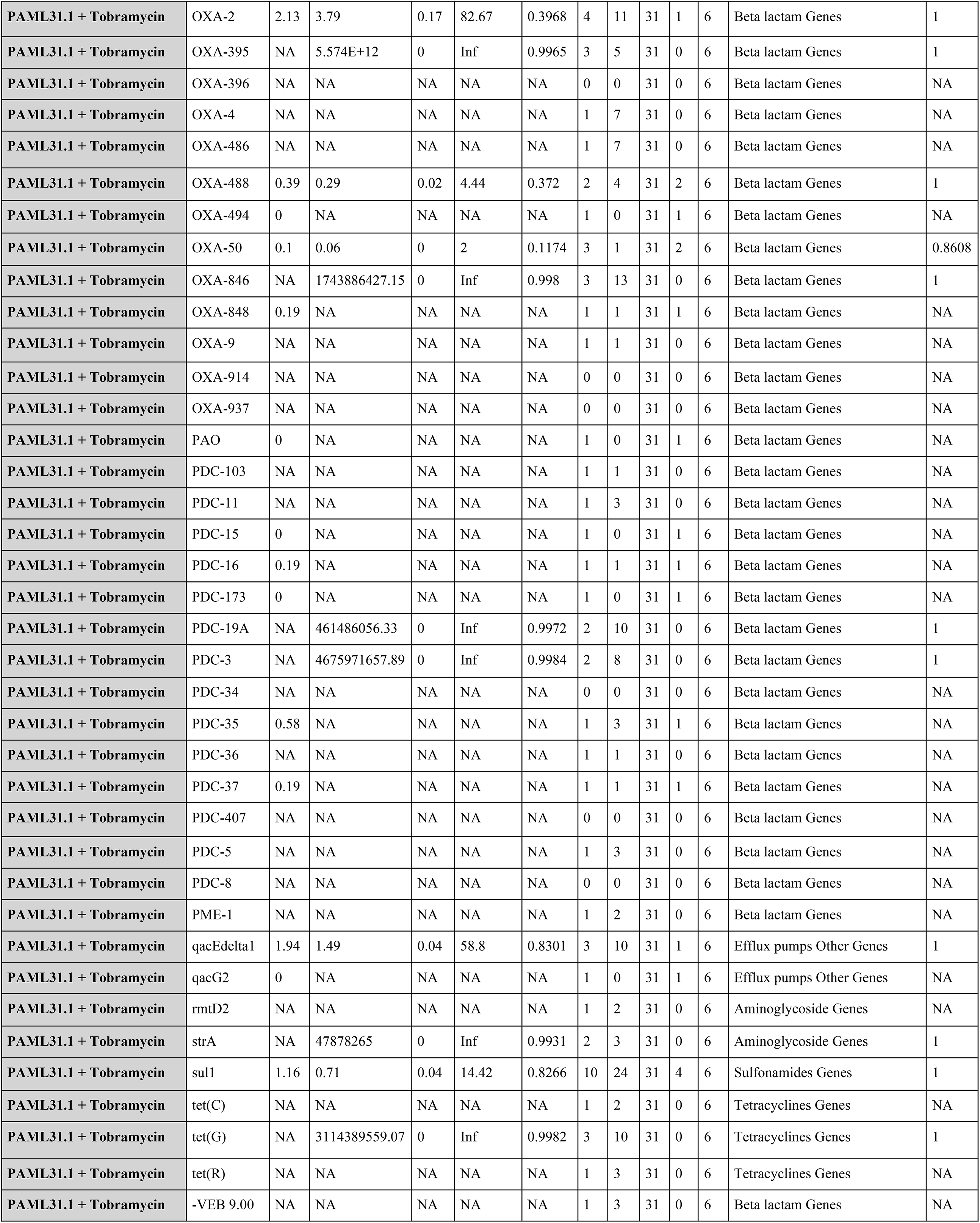
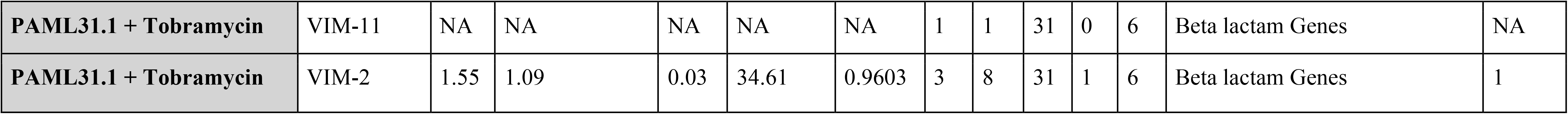
Analysis of gene enrichment/depletion in phage-resistant groups of strains among antibiotic-resistant groups of strains, as reported by a single mixed model adjusting for phylogenetic position and clonal lineage. Pairing, the particular phage and antibiotic being examined; Gene, the resistance gene being examined for presence/absence in various groups of strains; FC, fold change in phage-resistant group compared to phage-susceptible group; Odds Ratio, the ratio of the odds of being phage-resistant versus phage-susceptible within the antibiotic-resistant group of strains after controlling for phylogenetic position and clonal lineage; CIL, 95% Wald Confidence interval lower bound; CIU: 95% Wald Confidence interval upper bound; Pval: unadjusted p value; LW, total number of lineages with the gene; PRW, number of phage resistant strains that carry the gene (out of the antibiotic resistant strains); PR, number of phage resistant strains total (out of the antibiotic resistant strains); PSW, number of phage susceptible strains that carry the gene (out of the antibiotic resistant strains); PS, number of phage susceptible strains total (out of the antibiotic resistant strains); Category, refers to type of resistance conferred by a particular gene; FDR, FDR-adjusted P value. If only 1 lineage carries the gene, statistical analysis throws NA because the gene effect is perfectly confounded by the lineage effect.

